# Mouse Single-Cell Long-Read Splicing Atlas by Ouro-Seq

**DOI:** 10.1101/2025.01.17.633678

**Authors:** Hyunsu An, Sin Young Choi, Jawoon Yi, Chaemin Lim, So-I Shin, Sera Oh, Rosalind H. Lee, Jihwan Park

## Abstract

Alternative splicing creates cellular and phenotypic diversity; thus, characterizing splicing patterns across diverse cell types is essential for a comprehensive understanding of these cell types. However, existing full-length single-cell RNA-sequencing methods suffer from multiple technical limitations, such as the pervasive presence of reverse transcription artifacts across genes. Here, we introduce Ouro-Seq, a novel single-cell long-read sequencing framework with a versatile artifact removal mechanism for characterizing full-length mRNA species expressed by individual cells. We also developed a computational pipeline that identifies genuine full-length cDNAs and normalizes full-length mRNA size distributions for integration. Using Ouro-Seq, we constructed the Mouse Single-Cell Long-Read Splicing Atlas by collecting full-length transcriptomes of 103,304 nuclei or cells from 12 major organs and tissues of adult mice. Our atlas comprehensively characterizes alternative splicing, alternative promoter usage, and alternative polyadenylation events across cell types, revealing previously unappreciated functional heterogeneity among individual cells. Different cell types exhibit largely distinct expression patterns of various isoforms, corresponding to their specific morphologies and biological functions. Our splicing atlas and generalized long-read frameworks will empower researchers to fully interpret biological variation across full-length transcriptomes of single cells.

## INTRODUCTION

Through alternative splicing^1^, a human genome expresses at least several hundreds^2^ of thousands of unique mRNA isoforms from ∼20,000 genes, each of which possibly has a distinct biological function (e.g., encoding a functionally distinct protein variant). However, despite the strong evidence that these different isoforms will display significant functional differences^3^, most alternative transcripts remain poorly understood^4^. The functional annotation of these mRNA species would enable a deeper understanding of the functional heterogeneity of cells^5^ and provide new ideas for more efficient characterization of the complete repertoire of mRNA species expressed in the genome. Thus, mapping the expression patterns of individual mRNA isoforms across different cell types is the first crucial step in identifying their functions, which in turn provides a better understanding of the cell types. Additionally, cell type-specific isoform usage information could aid in identifying disease-associated splicing variants, variant prioritization for diagnosing^6^, understanding rare genetic diseases^7^, and more sensitive detection of cancer neoantigens^8^.

Studying transcriptomes at a single-cell resolution has revolutionized many fields in biological sciences, unmasking previously unknown cellular heterogeneity in various organs and tissues^9–12^. However, the fragmentation steps for short-read sequencing strongly limit the characterization of full-length transcriptomes in single cells (**Supplementary Note 1**). For example, the Smart-seq2 protocol allows analyzing splice junctions across the entire transcript length via short-read sequencing; however, the full-length transcript structures have to be inferred^13^ from these short-reads with the help of transcript assembly tools, which has been known to assemble only 20%–40% of the human transcriptome^14,15^. High-throughput single-cell RNA-sequencing (scRNA-seq) methods that utilize massively parallel cell barcoding generate a pooled full-length cDNA library as an intermediate product after reverse transcription^16^ (**Supplementary Note 1**). With the recent advent of long-read sequencing technologies, increasing efforts have been made to analyze single-cell full-length cDNA libraries without fragmentation, significantly improving our understanding of complex alternative splicing behaviors in individual cells^17–21^. However, these efforts revealed significant limitations in current long-read scRNA-seq technologies, requiring solutions for broader adoption in biomedical research. First, a significantly large yet highly variable portion (50%–98%) of molecules in a single-cell full-length cDNA library comprise reverse transcription (RT) and PCR artifacts^20–22^, which are truncated fragments of the original full-length mRNA molecule. These artifacts lack cell barcodes and cannot be analyzed further, reducing the overall sequencing efficiency and significantly confounding downstream analyses such as isoform quantification, data integration, and *de novo* transcriptome assembly. Another critical limitation is the bias toward shorter molecules (<2 kbp) introduced during the artifact removal process. Lastly, the long-read scRNA-seq library often contains a high amount of mitochondrial genome (mtRNA) and ribosomal RNA (rRNA)-derived cDNA molecules, significantly reducing the sequencing efficiency and resulting in inadequate sequencing coverages for other protein-coding genes of interest.

To address these issues, we developed Ouro-Seq, a novel long-read scRNA-seq framework designed to eliminate barcoding artifacts robustly across a wide range of cDNA sizes. This framework comprises both experimental and computational modules, including Ouro-Deplete and Ouro-Enrich for depleting and enriching the cDNAs derived from selected cells and transcripts in the long-read scRNA-seq library, respectively, Ouro-SizeSelect for robust enrichment of longer cDNAs from the library, and Ouro-Tools for integrative analysis of multiple long-read scRNA-seq data. Using Ouro-seq, we constructed the mouse single-cell splicing atlas at the resolution of individual full-length mRNA species, addressing the urgent need to understand biology at the resolution of individual cells and individual mRNA species.

## RESULTS

### Ouro-Seq, an efficient long-read scRNA-seq method

Since long-read sequencing technologies have been applied to analyze single-cell cDNA libraries, few methods have been developed to deplete cell barcode-free artifact molecules: biotin selection-based^20,23^ and asymmetric PCR-based^21^ methods. However, these methods display a strong length bias, preferring smaller molecules < 1 kbp. Consequently, previous long-read scRNA-seq studies often reported a molecule size distribution centered at ∼700 bp, with no or only a small fraction of molecules larger than 2 kbp^24–26^, leading to poor characterization of more than half of the human transcriptome^27,28^ (**Extended Data Figure 1a, b**). Specifically, biotin selection-based methods preferentially select smaller molecules because of their lower steric hindrance and faster diffusion rate. Additionally, asymmetric PCR-based methods exponentially increase the proportion of the specific type of reverse transcription artifacts containing cell barcodes at both ends (i.e., cell barcode-doublet artifacts, **Supplementary Note 2**).

Therefore, we sought to develop a novel artifact removal method for high-throughput long-read scRNA-seq. As both types of barcoding artifacts have either two 5’ or two 3’ PCR handles (i.e., two “tails” or two “heads”), respectively, we employed a selective self-cyclization reaction that only circularizes DNA molecules containing both 3’ and 5’ PCR handles followed by an exonuclease treatment to deplete both types of artifacts (**Figure 1a**). We thus developed a novel long-read scRNA-seq framework termed “Ouro-Seq” that removes both cell barcode-free and cell barcode-doublet artifacts without length bias (**Methods**). Another problem of long-read scRNA-seq is the presence of less informative yet highly abundant cDNA molecules derived from mitochondrial DNA (mtDNA) transcripts and ribosomal RNAs (rRNAs), which often constitute 30%–70% of a single-cell full-length cDNA library. To address this problem, we developed Ouro-Deplete (**Figure 1a**), which has an additional CRISPR-Cas9-based *in-vitro* digestion step to linearize circularized unwanted molecules before removing any linear DNA molecules in the library (**Methods**).

**Figure 1.**
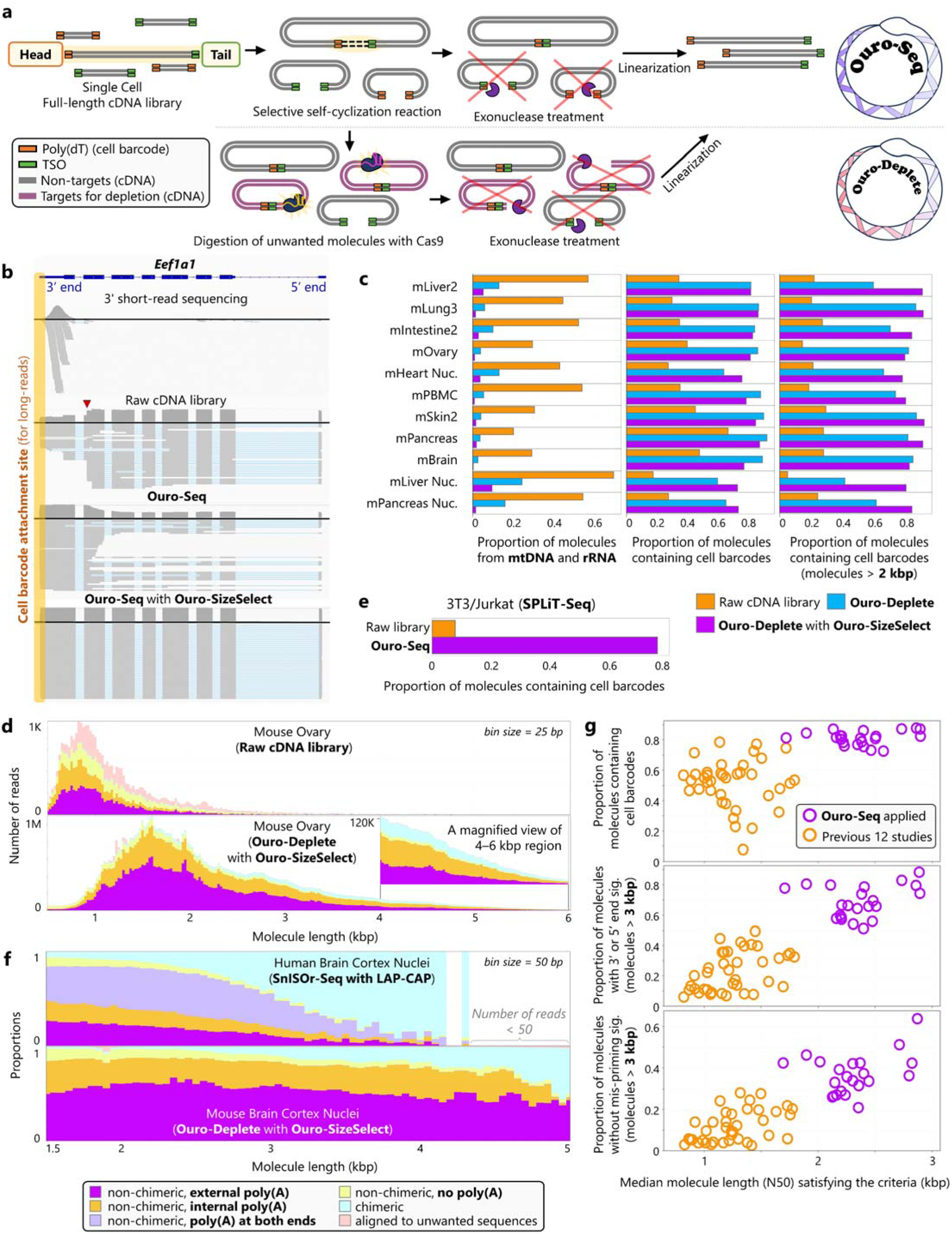
Comprehensive single-cell full-length transcriptome profiling using Ouro-Seq. **a**, Schematic representations of the Ouro-Seq and Ouro-Deplete work processes. (Upper panel) Ouro-Seq utilizes a selective self-cyclization reaction with minimal length-bias to efficiently deplete cell barcode-doublet (two “heads”) and cell barcode-free artifacts (two “tails”). (Lower panel) In addition to the comprehensive artifact removal capability of Ouro-Seq, Ouro-Deplete has the capability to deplete less informative yet highly abundant cDNA molecules in the library. **b**, Read coverage patterns of *Eef1a1* from the short-read sequencing library and long-read sequencing libraries from various stages of our long-read scRNA-seq library preparation process (sample# mLung3). The read coverage pattern of *Eef1a1* provides a good proxy for measuring how effectively cell barcode-free artifact molecules were depleted in the library. The location of a possible mispriming site responsible for the generation of reverse transcription artifacts is indicated with an arrowhead (“ACATGGG,” chr9:78,478,991–78,478,997 of *GRCm38*). **c**, Ouro-Deplete and, subsequently, Ouro-SizeSelect were applied to 11 single-cell and single-nucleus cDNA libraries prepared from eight mouse organs (**Extended Data Table 1**) to evaluate their robustness. **d**, **f**, Ouro-Seq enables efficient artifact removal across a wide molecule size range with minimal length bias. Colors represent different classes of cDNA molecules. **d**, Read counts before (above) and after (below) applying Ouro-Seq (Ouro-Deplete with Ouro-SizeSelect) (sample# mOvary). **f**, Proportions after applying an asymmetric PCR-based artifact depletion method (SnISOr-Seq with LAP-CAP, above) or Ouro-Seq (below). **e**, Ouro-Seq is compatible with various single-cell capture platforms, including the combinatorial indexing-based scRNA-seq technology SPLiT-Seq. **g**, Benchmark results of various single-cell long-read methods. cDNA molecules without the signatures of 5’ (template-switching oligo) or 3’ (poly(dT) primer) mispriming events are considered to have valid 5’ or 3’ cDNA ends, respectively.

Because of the length biases of mRNA capture and PCR amplification, a cDNA size distribution of a single-cell cDNA library often differs significantly from the expected mRNA size distribution^28^ (**Extended Data Figure 1a**). Therefore, we developed a precipitation-based DNA size selection method, Ouro-SizeSelect, which can robustly enrich up to 6 kbp of cDNA molecules from a single-cell full-length cDNA library (**Methods**). Ouro-SizeSelect employs a sequence of precipitation-based size selection steps (**Extended Data Figure 1c**), each of which gently pushes the size distribution of a library toward longer DNA molecules. When we used Ouro-Seq and Ouro-SizeSelect to deplete artifacts derived from reverse transcription and PCR and enrich longer cDNA molecules (2–6 kbp), their effects on the sequencing read coverage were apparent for genes generating a high proportion of artifact molecules and expressing long mRNA transcripts, respectively. For example, *Eef1a1* expresses a 1.8 kbp-long transcript containing several template-switching-oligo (TSO)-binding sites that reverse transcriptases can utilize to generate cell barcode-free artifact molecules (the most prominent site is ACATGGG, indicated by the red arrowhead in **Figure 1b**). Before applying Ouro-Seq, 65% of sequencing reads aligned to *Eef1a1* were cell barcode-free artifact molecules. After applying Ouro-Seq, virtually all sequencing reads aligned to *Eef1a1* contained cell barcodes, but the majority (56%) of reads captured truncated mRNAs (∼600 bp) before Ouro-SizeSelect. Interestingly, the truncation positions were clustered within ∼200 bp downstream of the TSO-binding sites (**Figure 1b**), which was consistently observed across genes containing TSO-binding sites, suggesting possible mechanisms by which 5’-truncated cDNAs are primarily generated from these genes. As expected, after applying Ouro-SizeSelect, all sequencing reads aligned to *Eef1a1* were cell barcode-containing, full-length *Eef1a1* transcripts (**Figure 1b**), indicating that cDNA molecules of longer lengths are appropriately represented in the resulting library.

In raw single-cell cDNA libraries of various biological samples, the percentages of cDNA molecules from rRNAs and mtDNA were 30%–60% across the samples, significantly reducing the sequencing output for informative cDNAs (**Figure 1c**). After applying Ouro-Deplete (**Methods**) and Ouro-SizeSelect, the percentage dropped to negligible levels (<2%) for most samples (**Figure 1c** and **Extended Data Table 1**). As previously reported^21^, the percentage of cell barcode-containing, usable reads in raw libraries is 25%–50%, and even lower (15%–25%) when considering only cDNA molecules longer than 2 kbp (**Figure 1c**). However, Ouro-Deplete with Ouro-SizeSelect significantly improved the proportions of barcoded reads (>90%) across the entire size range of cDNA molecules (**Figure 1c, d** and **Extended Data Figure 1d**). We also applied Ouro-Seq to a single-cell cDNA library prepared using another scRNA-seq technology, SPLiT-seq (**Methods**). Ouro-Seq efficiently depleted cell barcode-free artifacts, increasing the fraction of barcoded reads in the library by an order of magnitude (**Figure 1e**). Overall, these findings indicate that Ouro-Seq is a flexible and robust method for the comprehensive characterization of biologically informative, full-length cDNA molecules across a wide size range, including most of the known mRNA transcripts in humans^27^ (55,294/61,292 for protein-coding transcripts and 256,967/265,480 for all transcripts, using Ensembl 105) at the single-cell level.

### Benchmarking reveals the unique strength of Ouro-Seq

We compared Ouro-Seq (i.e., Ouro-Deplete with Ouro-SizeSelect) with previously reported long-read scRNA-seq methods, covering 16 studies published since 2017 (**Methods**). Ouro-Seq successfully depleted cell barcode-doublet artifacts to a negligible fraction (0.05%–0.1%) of the library (baseline, 0.5%–1%). When using the asymmetric PCR-based method^21^, more than half of the cell barcode-containing molecules larger than 2 kbp were cell barcode-doublet artifacts (**Figure 1f** and **Extended Data Figure 2a**, “non-chimeric, poly(A) at both ends”) (see **Supplementary Note 2** for more details). In addition, the asymmetric PCR-based method displayed a significant bias toward shorter molecules, reducing the fraction of molecules longer than 2 kbp from 10% to 3% (**Extended Data Figure 2b**). The biotinylated cDNA pull-down method^26^ effectively depleted cell barcode-free artifacts like the asymmetric PCR method but showed an even stronger bias toward shorter molecules (**Extended Data Figure 3a**). After applying the biotin selection-based method, 15%–25% of molecules in the size range of 2–3 kbp contained a cell barcode, which was significantly lower than the percentage (∼90%) obtained with Ouro-Seq (**Extended Data Figure 3b)**. Furthermore, the biotin selection-based method was ineffective at removing cell barcode-doublet artifacts, accounting for ∼1% after applying the method, unchanged from the baseline level.

Finally, we compared the cell barcode-free artifact depletion efficiency and the fraction of truncated cDNAs of various long-read scRNA-seq methods, covering 60 datasets in total (**Figure 1g** and **Extended Data Table 2**). For the 38 datasets from the previous studies, the fraction of cell barcode-containing molecules varied widely from 8% to 78% and was generally lower than that of the Ouro-Seq libraries (75%–90%). Next, we searched for reverse transcription signatures indicating mispriming events. For example, a genome-encoded poly(A) tract in a cDNA molecule indicates that the molecule was generated through a mispriming event at an internal poly(A) tract of an mRNA transcript. Further, if a 5’ end of a cDNA molecule represents a genuine 5’ end of an mRNA molecule *in vitro*, the three consecutive guanosine nucleotides at the 5’ end of the cDNA (originating from the 3’ end of the TSO) are not encoded in the genome (i.e., the “external Gs” signature, **Extended Data Figure 1d**). Surprisingly, we found that the fraction of molecules without any mispriming signature, possibly representing *in-vivo* full-length mRNA transcripts, was significantly higher in the Ouro-Seq libraries, particularly for longer cDNA molecules (>3 kbp) (**Figure 1g**). However, in samples processed using other artifact removal methods, the fraction of molecules without mispriming events decreased rapidly as molecule length increased (“non-chimeric, external poly(A), and external Gs” class in **Extended Data Figure 3a** and **Extended Data Figure 2a**, respectively). In contrast, for samples processed using Ouro-Seq, the fraction of molecules without a mispriming signature remained unchanged across the entire size range covered by Ouro-Seq. These observations suggest that the strong length bias of the previous methods inadvertently enriched truncated cDNAs generated by mispriming events, which are significantly shorter than the full-length cDNAs representing the original mRNA transcripts.

### A novel framework for target-enriched long-read scRNA-seq

Existing target enrichment methods for full-length cDNAs, such as hybridization capture-based methods^29,30^, require at least an overnight reaction (∼12 h) and generally have a high per-sample cost. While the alternative, PCR-based methods^31^ require much less time (∼1 h), the throughput is substantially lower (up to dozens of genes per reaction), and the full-length information encoded in the targets is inevitably lost in the PCR products. The flexibility of our framework enabled us to develop a novel CRISPR-based target enrichment method, Ouro-Enrich (**Figure 2a**), for efficient and rapid (∼2 h) enrichment of target full-length cDNA molecules in a high-throughput and highly cost-effective manner (**Methods**). Ouro-Enrich replaces the final step of Ouro-Seq, i.e., the linearization of all circularized cDNA molecules, with CRISPR-Cas9-based selective linearization of target molecules. As a result, only target molecules are linearized, which can be directly ligated with long-read sequencing adapters or Y-shaped adapters for additional PCR amplification. As a proof-of-concept experiment, when we enriched full-length cDNA molecules of 11 mitochondrial DNA-encoded genes from a single-cell cDNA library (**Methods**), the fraction of mtDNA-encoded transcripts increased more than 20-fold, from 4% to 95% (**Figure 2b**). When we applied Ouro-Enrich for cDNA molecules covering 13 single-nucleotide variants (SNVs) of interest to a single-cell cDNA library^32^ of a patient with acute myeloid leukemia (**Methods**), we similarly achieved more than 20-fold enrichment for the target SNVs (**Figure 2c**).

**Figure 2.**
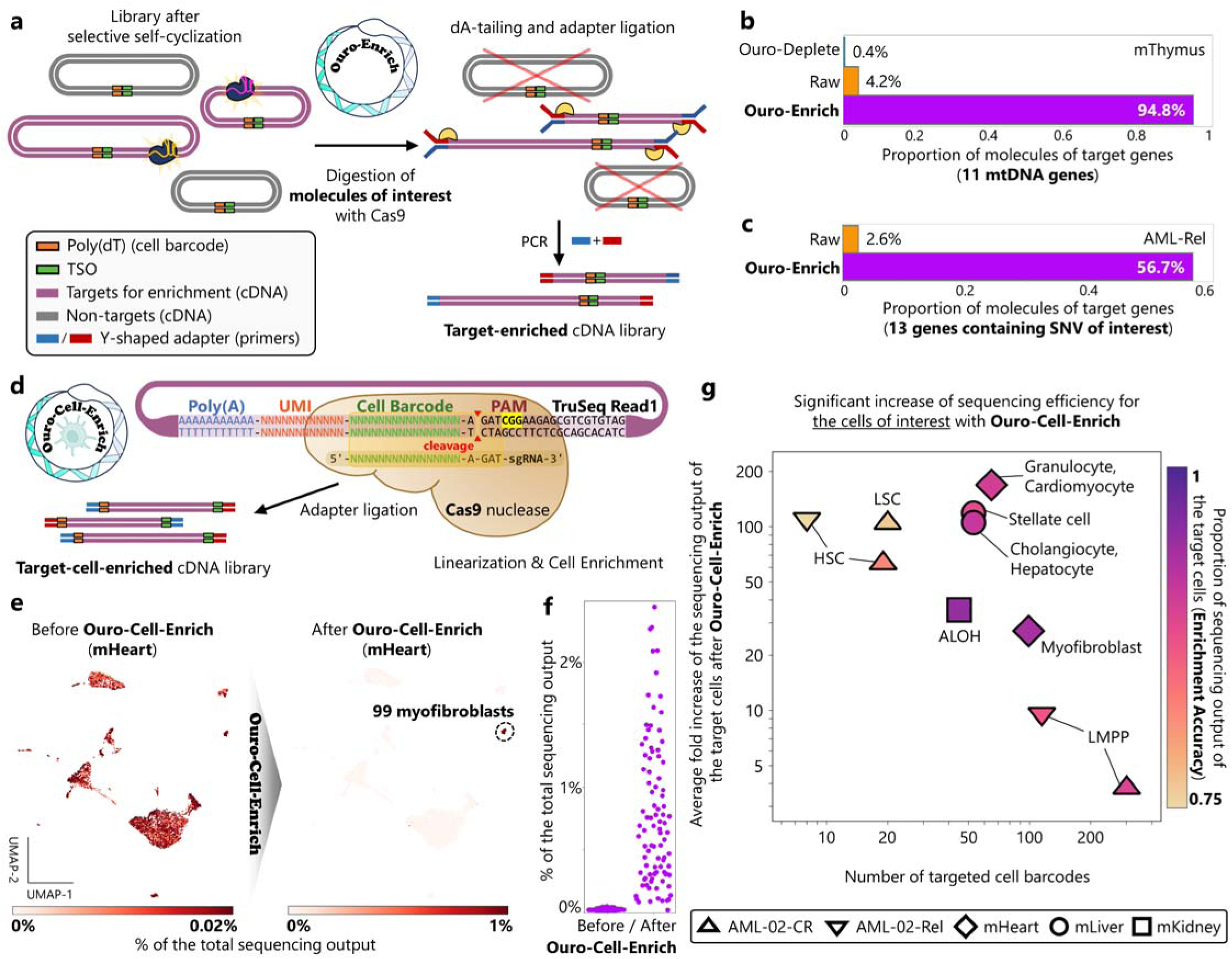
A versatile long-read enrichment framework, Ouro-Enrich. **a**, Schematic representation of Ouro-Enrich targeted enrichment. **b**, A single-cell cDNA library containing the lowest fraction of mtDNA-derived transcripts was selected (sample# mThymus), and Ouro-Enrich was applied for targeted enrichment of 11 mtDNA-derived transcripts. **c**, Ouro-Enrich was applied to a long-read scRNA-seq library of a patient with relapsed acute myeloid leukemia (sample# AML-02-Rel) to enrich cDNA molecules containing single-nucleotide variants in 13 genes of interest. **d**, High-throughput targeted cell enrichment with Ouro-Cell-Enrich for 10x Genomics scRNA-seq and spatial transcriptomics libraries as an exemplary implementation. **e**, **f**, Percentages of total long-read sequencing output of individual cells before (left) and after (right) applying Ouro-Cell-Enrich for the enrichment of 99 myofibroblasts from a pooled long-read scRNA-seq library of ∼7,000 cells from the mouse heart, visualized with (**e**, UMAP embeddings; detailed cell type annotations can be found in **Extended Data Figure 4**) and without (**f**, strip plots) non-target cells. **g**, Results of various Ouro-Cell-Enrich experiments with targeted cell barcode numbers varying from 8 to 301 cells. PAM, protospacer adjacent motif; UMI, unique molecular identifier; HSC, hematopoietic stem cell; LSC, leukemic stem cell; ALOH, epithelial cell of the ascending limb of the loop of Henle; LMPP, lympho-myeloid primed progenitor cell.

A high-throughput single-cell cDNA library typically contains cDNA molecules of hundreds to even millions of cells^33^. However, because of the relatively high per-read cost of long-read sequencing, it is often more reasonable to focus the analysis on a subset of the cells. Therefore, we devised Ouro-Cell-Enrich (**Figure 2d**), a specific implementation of Ouro-Enrich, to confidently enrich full-length cDNAs of particular cells of interest in a massively parallel manner using the CRISPR-Cas9 system (**Methods**). The CRISPR-Cas9 system recognizes target sites of 17–20 nucleotides in length^34^, which is close to the typical length of a cell barcode sequence (10–16 bp)^16^ in scRNA-seq libraries. By selecting an appropriate protospacer adjacent motif sequence around the cell barcode sequence (e.g., in a TruSeq Read1 adapter sequence), the entire length of the cell barcode sequence (16 bp) can be fully recognized by the CRISPR-Cas9 system, allowing accurate target cell barcode enrichment (**Figure 2d**). By applying Ouro-Cell-Enrich, we successfully enriched full-length cDNA molecules of various cell types of interest, enabling selective analysis of the full-length transcriptomes of the selected cells (**Figure 2e–g**). Initially, we enriched cDNAs of 99 cardiac myofibroblasts from a long-read scRNA-seq library of ∼7,000 cells isolated from a mouse heart (**Methods**). Before target cell enrichment, the cDNA molecules of the cardiac myofibroblasts accounted for only 2.3% of the total long-read sequencing output (**Extended Data Figure 4** and **Figure 2e**). However, after applying Ouro-Cell-Enrich, the sequencing output for the 99 cardiac myofibroblasts significantly increased to 94%, achieving a nearly 40-fold increase in sequencing coverage for the target cells (**Figure 2e, f**). We subsequently applied Ouro-Cell-Enrich for nine other cell types of interest, with varying numbers of cells (8–301 cells), in four long-read scRNA-seq libraries, targeting a total of 778 cells from 26,197 annotated cells across the libraries (**Methods**). We achieved consistently accurate enrichment of the target cells across the samples, regardless of the number of target cells, with an average enrichment accuracy of ∼90% (**Figure 2g**). In summary, Ouro-Enrich and Ouro-Cell-Enrich provide convenient and accessible toolkits with unique capabilities (**Supplementary Methods**) to the research community for analyzing specific transcripts or cells of interest using long-read sequencing.

### Ouro-Tools minimizes technical variations *in silico*

To comprehensively characterize single-cell full-length cDNA libraries, we developed Ouro-Tools (**Figure 3a**), a general computational framework for processing and analyzing multi-sample long-read scRNA-seq data (**Methods**). Compared with existing long-read scRNA-seq pipelines^20^, Ouro-Tools has several unique features. For example, by analyzing the patterns of non-templated nucleotide additions during reverse transcription, Ouro-Tools identifies *de novo* transcription start sites^35^ (TSSs). Moreover, Ouro-Tools performs *in-silico* mRNA size distribution normalization of each long-read sequencing experiment while integrating multiple datasets. To our knowledge, Ouro-Tools is the first bioinformatics framework that corrects technical variation in the mRNA size distribution for differential transcript usage (DTU) analysis. Additionally, Ouro-Tools supports single-cell analysis of insufficiently characterized genomic loci, including transcribed cis-regulatory elements (tCREs) and transcripts derived from transposable elements (TEs) (**Figure 3a**). Further, Ouro-Tools provides a quick and comprehensive quality control report, including the sequence length distributions of cell barcode-free and -doublet artifacts, PCR chimeric molecules, and molecules generated by mispriming events (examples are shown in **Extended Data Figures 2–3**).

**Figure 3.**
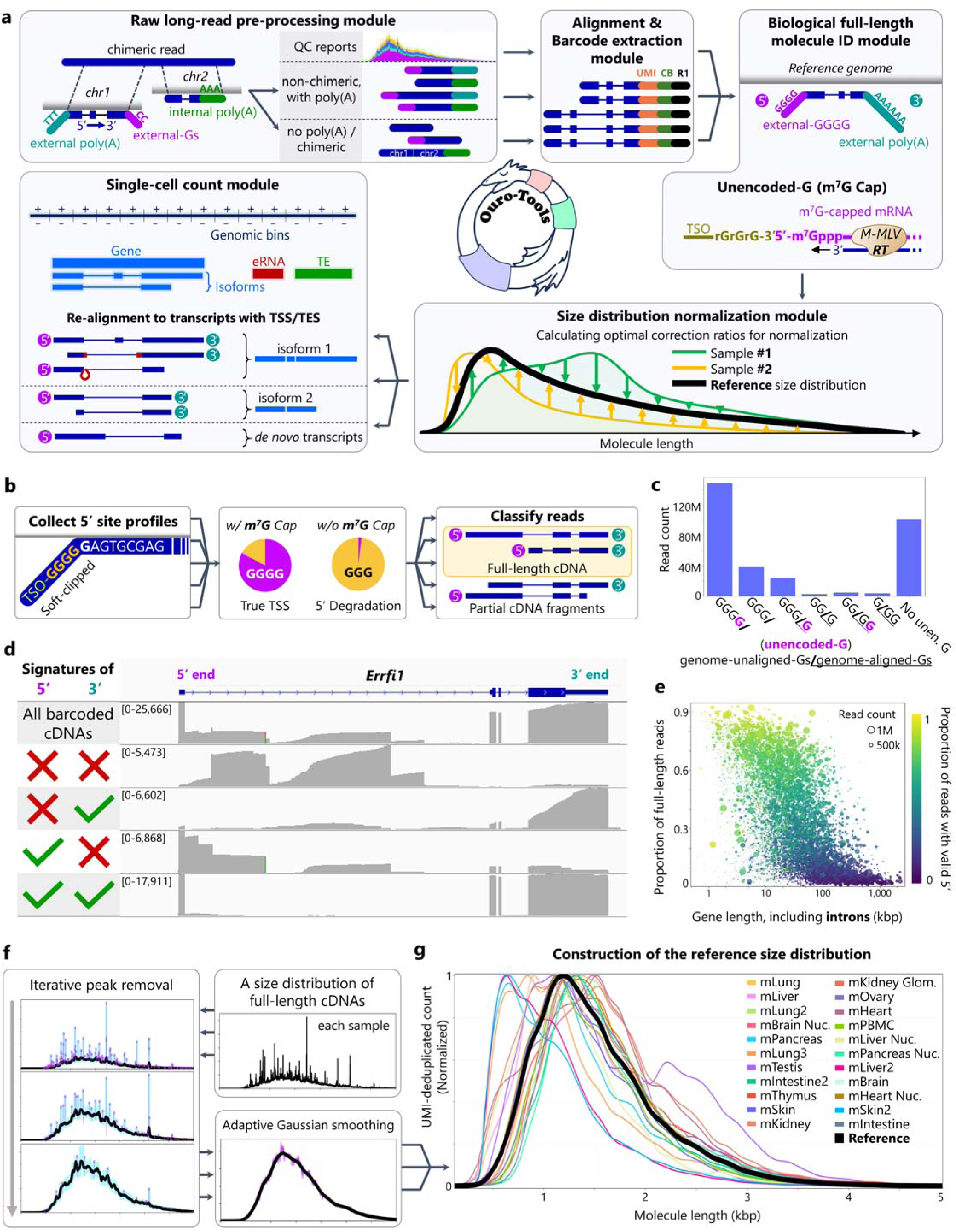
Comprehensive analysis of full-length transcriptomes at the single-cell level with Ouro-Tools. **a**, A brief overview of Ouro-Tools. The Ouro-Tools pipeline is designed to minimize technical variations specific to long-read RNA-seq protocols *in silico*, such as reverse transcription (RT) and PCR artifacts, truncated cDNA molecules, and a variation of read length distributions across different long-read sequencing experiments. **b**, Full-length identification pipeline of Ouro-Tools. The pipeline collects soft-clipped G homopolymer lengths (or C homopolymer lengths for reverse complemented reads) across all samples, creating a histogram (a 5’-site profile) of the soft-clipped G/C homopolymer lengths for each candidate TSS across the genome. **c**, Numbers of reads assigned to different 5’-site classes across 22 samples from 12 mouse organs and tissues. **d**, The Ouro-Tools pipeline enables *in-silico* filtering of truncated cDNA molecules, exemplified by the filtered reads representing *in-vivo* full-length mRNAs transcribed from *Errfi1*. **e**, Ouro-Seq characterizes the majority of mouse genes at full length. **f**, The underlying mRNA size distribution of each sample is estimated through iterative peak removal and adaptive smoothing steps. **g**, To adequately represent all mRNA transcripts profiled in all samples (**Extended Data Table 1**), the reference mRNA size distribution is constructed, which is subsequently utilized to export a size distribution-normalized count matrix for each sample. eRNA, enhancer RNA; TE, transposable element-derived transcript; unen. G, unencoded-G.

Currently, all scRNA-seq platforms that generate a full-length cDNA library utilize Moloney murine leukemia virus (M-MLV) reverse transcriptase for reverse transcription, which adds one “non-templated” cytidine for mRNAs containing the m7G cap^36^ (**Figure 3a**), which has been utilized to identify locations of *de novo* TSSs^35,37^ using short-read sequencing. However, for long-read sequencing, no such bioinformatic method is available. Therefore, we implemented the biological full-length identification module of Ouro-Tools (**Methods**). When we applied the pipeline to 22 long-read scRNA-seq datasets, 50%–90% of cDNAs that started from genuine TSSs contained an unencoded G. In contrast, the fraction of cDNAs with an unencoded G was significantly lower (<5%) for artifactual TSSs of cDNAs generated by mispriming events and partially degraded, uncapped mRNAs (**Figure 3b**). Across the Ouro-Seq datasets, more than half of the barcoded cDNA molecules were identified as originating from genuine TSSs (**Methods**), totaling 177 million cDNAs after unique molecular identifier (UMI) deduplication (**Figure 3c**). Subsequently, the pipeline examines each properly barcoded cDNA molecule to determine whether the molecule is a full-length cDNA molecule containing both an unencoded G and a poly(A) tail sequence that is not encoded in the genome. Interestingly, signatures of intact 5’ and 3’ ends were sufficient for the clear separation of full-length cDNA molecules from truncated cDNA molecules (**Figure 3d** and **Extended Data Figure 5a, b**), which likely originated from partially degraded mRNA molecules with 5’ truncations^38^, lariat intronic RNAs^39^ containing mispriming sites, or the full-length cDNAs themselves through mispriming events. As expected, the proportion of full-length cDNA molecules varied more widely for larger genes, particularly genes longer than 20 kbp (**Figure 3e**), because of the higher probability of having mispriming sites in their longer intronic regions. However, compared to other barcoding artifact removal methods^21,26,40^, Ouro-Seq displayed a superior performance for identifying *in-vivo* full-length cDNAs across all protein-coding genes (**Extended Data Figure 5c, d** and **Supplementary Table 4**), indicating that Ouro-Seq is an excellent tool for characterizing full-length mRNA species expressed by individual single cells.

Despite the observation that the underlying mRNA size distributions of most organs and tissues are nearly identical (**Extended Data Figure 5e** and **Supplementary Methods**), the average read length in long-read RNA-seq experiments varies considerably across protocols and sample types^41,42^. During the benchmarking of long-read scRNA-seq methods, we noticed a significant variation in average cDNA molecule size among the samples (varying from 800 bp to 1,700 bp), even when they were processed using the same method (e.g., 800–1,250 bp for SnISOr-Seq, **Extended Data Table 2**). Such variation in read length distributions can introduce a strong batch effect that prevents accurate differential transcript usage (DTU) detection across samples. For example, consider a gene with two isoforms of 1 kbp and 2 kbp in length that are expressed at the same level in all cell types. However, without adjustment for read length distribution, the detected ratio of the two expressed isoforms will be the ratio of the relative frequencies of the reads of 1 kbp and 2 kbp in length in a read length distribution of each sample, regardless of the true ratio. Ouro-Seq can minimize the difference in cDNA size distributions among samples by pooling the amounts of Ouro-SizeSelect-applied libraries of different peak cDNA sizes (**Extended Data Figure 1c**) with a fixed ratio for every sample (**Methods**). However, we still observed subtle differences in cDNA size distributions among the datasets, which may have been introduced by DNA quantification errors from varying amounts of PCR bubbles and the long-read sequencing processes themselves (**Supplementary Methods**). The first step of the size distribution normalization module of Ouro-Tools is identifying underlying mRNA size distributions of individual samples by iteratively removing highly abundant mRNA species, which manifest as “peaks” in a full-length cDNA size distribution of each sample (**Figure 3f**). To faithfully represent low-coverage regions of an mRNA size distribution while effectively removing local variation in high-coverage regions, we implemented an adaptive smoothing algorithm that utilizes a series of Gaussian filters with different standard deviation (σ) values (**Figure 3f** and **Supplementary Methods**). Finally, a reference mRNA size distribution of the samples is constructed as the geometric mean of the underlying mRNA size distributions of the samples (**Figure 3g**). Optimal correction ratios are thus determined for each sample using a grid search algorithm, which is subsequently utilized by the single-cell count module of Ouro-Tools to construct a size distribution-normalized count matrix (**Figure 3a**). Overall, Ouro-Tools is designed to minimize the technical variation specific to long-read scRNA-seq protocols, enabling accurate integration of datasets and sensitive detection of alternative splicing events.

### Single-cell long-read splicing atlas of *Mus musculus*

We utilized Ouro-Seq to compile a “Mouse Single-Cell Long-Read Splicing Atlas” comprising the full-length mRNA transcripts expressed by more than 100,000 single cells collected from 12 organs and tissues of *M. musculus* (**Figure 4a**). To characterize various cell types, including those that are too large (e.g., cardiomyocytes) or too fragile (e.g., neurons) for tissue dissociation, we utilized both single-nucleus and single-cell RNA-seq. To counter the strong bias of scRNA-seq data toward immune cell types, we additionally performed leukocyte depletion to preferentially characterize parenchymal cell types from various organs (**Supplementary Methods**). We collected 22 samples from the brain, heart, kidneys, intestine, liver, lungs, pancreas, skin, mammary glands, thymus, blood, testes, and ovaries (**Extended Data Table 1**). After Ouro-Seq, we successfully processed and integrated the long-read scRNA-seq datasets using Ouro-Tools (**Methods**). In total, we generated 3.4 trillion base pairs of Nanopore long-read sequencing data, 81% of which were of non-chimeric, properly barcoded genome-aligned cDNA molecules, more than half of which were longer than 2 kbp (**Extended Data Table 1**). Using the *in-vivo* full-length mRNA identification module of Ouro-Tools, we detected 217,046 and 161,220 previously annotated splice junctions and exons across 21,425 genes, 45,437 transcribed regulatory elements, and 30,754 transcribed transposable elements, representing a variety of full-length mRNA transcripts expressed by the cells captured in the atlas (**Supplementary Table 5**). Overall, we identified 215 cell types across the tissues, which were annotated using Cell Ontology to facilitate^43^ an exploration of our single-cell full-length transcriptomic atlas (**Methods**).

**Figure 4.**
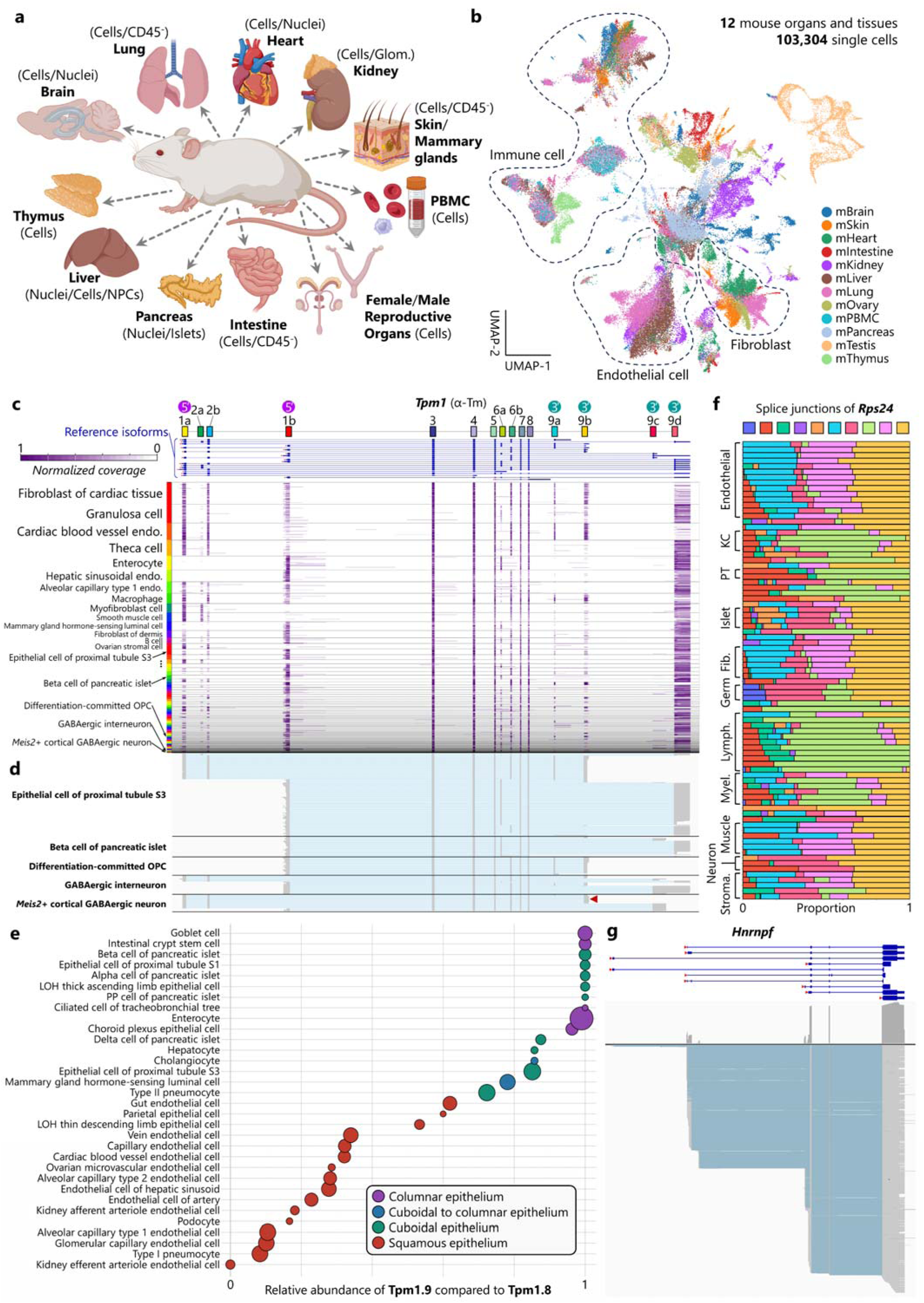
Overview of the Mouse Single-Cell Long-Read Splicing Atlas. **a**, Using Ouro-Seq and Ouro-Tools, a total of 12 organs and tissues from female and male mice were analyzed (created with BioRender.com accessed on 10 January 2024), compiling the Mouse Single-Cell Long-Read Splicing Atlas comprising full-length mRNA transcripts of single cells as-is, without fragmentation. **b**, UMAP representation of the atlas (n = 90,456 individual cells). Colors represent organs and tissues. In line with previous observations^12,75^, we observed moderate to high organ-specificity of the cell types resident in various tissues and organs, such as fibroblasts, macrophages, smooth muscle cells, and endothelial cells. **c**, Coverage patterns of full-length cDNAs aligned to *Tpm1* at the resolution of individual cells. Through combinations of alternative promoter usage, alternative polyadenylation, and alternative splicing, *Tpm1* generates 13 known protein isoforms (**Extended Data Figure 6**). Cell type labels based on Cell Ontology are shown. **d**, Individual full-length *Tpm1* transcripts for the selected cell types. The arrowhead indicates *de novo* Tpm1 isoforms in *Meis2*+ GABAergic neurons. **e**, Relative abundance of Tpm1.9 compared to Tpm1.8 across diverse epithelial cell types, calculated as Tpm1.9/(Tpm1.8 + Tpm1.9). **f**, Differential splice junction usage of *Rps24* across cell types (detailed cell type names can be found in **Extended Data Figure 7b**). **g**, Individual full-length *Hnrnpf* transcripts. NPC, hepatic non-parenchymal cell; OPC, oligodendrocyte precursor cell; LOH, the loop of Henle of nephron; KC, keratinocyte; PT, proximal tubule epithelial cell; Fib., fibroblast; Lymph., lymphoid cell; Myel., myeloid cell; Stroma., stromal cells.

To illustrate the complex landscape of cell type-specific transcript usage patterns, we explored transcript usage patterns of *Tpm1* (tropomyosin 1), which is ubiquitously expressed and encodes at least 13 distinct known protein isoforms^44^ through transcripts ranging in size from 1 to 3 kbp (**Extended Data Figure 6a**), across various cell types (**Figure 4c, d**). The variety of tropomyosins supports various cellular functions involving actin filaments^44^. Interestingly, we found that different cell types display largely mutually exclusive expression patterns of various isoforms based on their morphologies and biological functions. For example, simple squamous epithelial cells and other epithelial cells of flattened appearance (e.g., type I pneumocytes, endothelial cells, and podocytes) predominantly express Tpm1.8 (**Figure 4e** and **Extended Data Figure 6b**). In contrast, simple cuboidal and columnar epithelium cells (e.g., type II pneumocytes, enterocytes, goblet cells, and beta cells) predominantly express Tpm1.9. As expected, muscle-like cell types, such as vascular smooth muscle cells, pericytes, myofibroblasts, and extraglomerular mesangial cells, primarily express muscle tropomyosins (Tpms1.1–1.4), whereas non-muscle contractile cell types, such as fibroblasts across various organs, podocytes^45^, and ovarian theca cells, highly express stress fiber-associated tropomyosins (Tpms1.6 and Tpm1.7) (**Extended Data Figure 6c**). Furthermore, we observed a wide variety of expression patterns of *de novo Tpm1* transcripts with novel splice junctions^44^ in various cell types (**Supplementary Note 3**), such as oligodendrocyte precursor cells (OPCs), differentiation-committed oligodendrocyte precursor cells, mature oligodendrocytes, and *Meis2*+ GABAergic neurons (**Figure 4d**).

As another example^46,47^, we analyzed differential transcript usage of *Rps24* (ribosomal protein S24) transcripts across cell types. Previously, differentially spliced isoforms of *Rps24* have been associated with ribosomal specialization under specific conditions, such as oncogenesis^48^ or hypoxia^49^. However, the annotated mRNA transcripts of *Rps24* vary in size over an order of magnitude, from 468 bp to 4,059 bp, which poses a unique challenge for detecting differential transcript usage across experiments. As expected, cell type-specific expression patterns of *Rps24* transcripts of various lengths were fully captured in our atlas, including a 2.5 kbp-long *de novo Rps24* transcript that utilizes a unique combination of existing splice sites (**Extended Data Figure 7a**), which we visualized as proportions of splice junction usage across cell types (**Figure 4f** and **Extended Data Figure 7b**). We thus found that each cell type displays a characteristic *Rps24* transcript usage pattern associated mainly with its biological function. For example, *Rps24* transcript usage patterns were highly similar among various stromal cell types, including fibroblasts, myofibroblasts, pericytes, and smooth muscle cells from multiple organs and tissues. The complex alternative splicing patterns of *Cd47*^11^ were also captured in our single-cell splicing atlas (**Extended Data Figure 8a, b**). Moreover, using Ouro-Tools, our atlas accurately captured usage patterns of transcription start sites (TSSs) or transcription end sites (TESs) at the resolution of individual cells. For example, we observed that the majority of cell types utilize more than one promoter of *Hnrnpf*, a universally expressed gene encoding an RNA-binding protein that regulates alternative splicing^50^ and translation efficiency^51^, resulting in various cell type-specific alternative promoter and polyadenylation site usage patterns (**Figure 4g** and **Extended Data Figure 9**). Altogether, our Mouse Single-Cell Long-Read Splicing Atlas comprehensively characterized alternative splicing, alternative promoter usage, and alternative polyadenylation events across cell types.

### Deciphering functional heterogeneity of single cells using isoform information

Next, we analyzed the full-length transcriptomes of individual organs and tissues to identify various cell type-specific alternative splicing events. We developed the differential transcript usage detection module of Ouro-Tools that adaptively generates varying numbers of pseudo-bulk samples^52^ for each gene of interest based on the expression levels of the gene before applying conventional differential transcript usage analysis methods (**Supplementary Methods**). The number of cell type-specific transcripts varied across cell types for each organ and tissue (**Figure 5a** and **Extended Data Figures 10–20**). For instance, we verified the known alternative promoter usage of the claudin gene *Cldn10* (**Figure 5b**), which encodes tight junction proteins Cldn10a^53^ and Cldn10b^54^, which are responsible for paracellular Na^+^ and Cl^−^ transport in the proximal tubule and thick ascending limb (TAL), respectively, in the kidneys. Distinct ion permeabilities in different parts of the TAL have been attributed to spatially distributed mosaic patterns of tight junctions along the TAL^54^ (**Supplementary Note 4**). To better understand the cellular heterogeneity in the TAL^54^, we investigated transcript usage patterns of TAL cells based on *Cldn* expression patterns. Subclustering of TAL cells revealed four cell populations: *Cldn10*+ TAL, *Cldn16*+ TAL, *Cldn10*+ *Cldn16*+ TAL, and macular densa cells (**Figure 5c, d**). However, unique marker genes could not be confidently identified for the *Cldn10*+ *Cldn16*+ TAL population (**Extended Data Figure 21a**). To exclude the possibility that this population is a technical artifact consisting of doublets from other subpopulations of TAL, we analyzed differential transcript usage among the TAL cell populations. Interestingly, we observed that each cell population exclusively expresses a distinct isoform of *Slc12a1* in full length, linking a functionally distinct protein isoform of *Slc12a1* to each cell population (**Figure 5e–h**). *Slc12a1* encodes NKCC2, a kidney-specific Na-K-Cl cotransporter. Through alternative splicing, resulting in the inclusion of one of three mutually exclusive cassette exons (A, B, and F) encoding a transmembrane domain that interacts with the substrates, *Slc12a1* generates three functionally distinct protein isoforms: NKCC2-B, NKCC2-A, and NKCC2-F^55^. Each NKCC2 isoform has different binding affinities for Na^+^, K^−^, and Cl^− 56^ and is expressed in a different region of the TAL^55^. Additionally, we found a novel *Fxyd2* isoform predominantly expressed by the *Cldn16*+ TAL and *Cldn10*+ *Cldn16*+ TAL populations, while the two known *Fxyd2* isoforms^57^, encoding FXYD2-a and FXYD2-b, were abundantly expressed in the *Cldn10*+ TAL cell population (**Extended Data Figure 21b**). Based on the previously reported locations of tight-junction mosaic patterns^54^ and known protein isoforms of *Fxyd2*^57^ and *Slc12a1*^55^ along the corticomedullary axis of the TAL, our splicing atlas provides an improved model explaining how functional specialization of paracellular and transcellular ion transport along the TAL is achieved through differential localization of and spatially restricted interactions among distinct cell populations in the TAL (**Supplementary Note 5**). In summary, our full-length transcriptome atlas captured the usage of functionally distinct isoforms at the single-cell level, revealing functional heterogeneity within a poorly characterized cell population.

**Figure 5.**
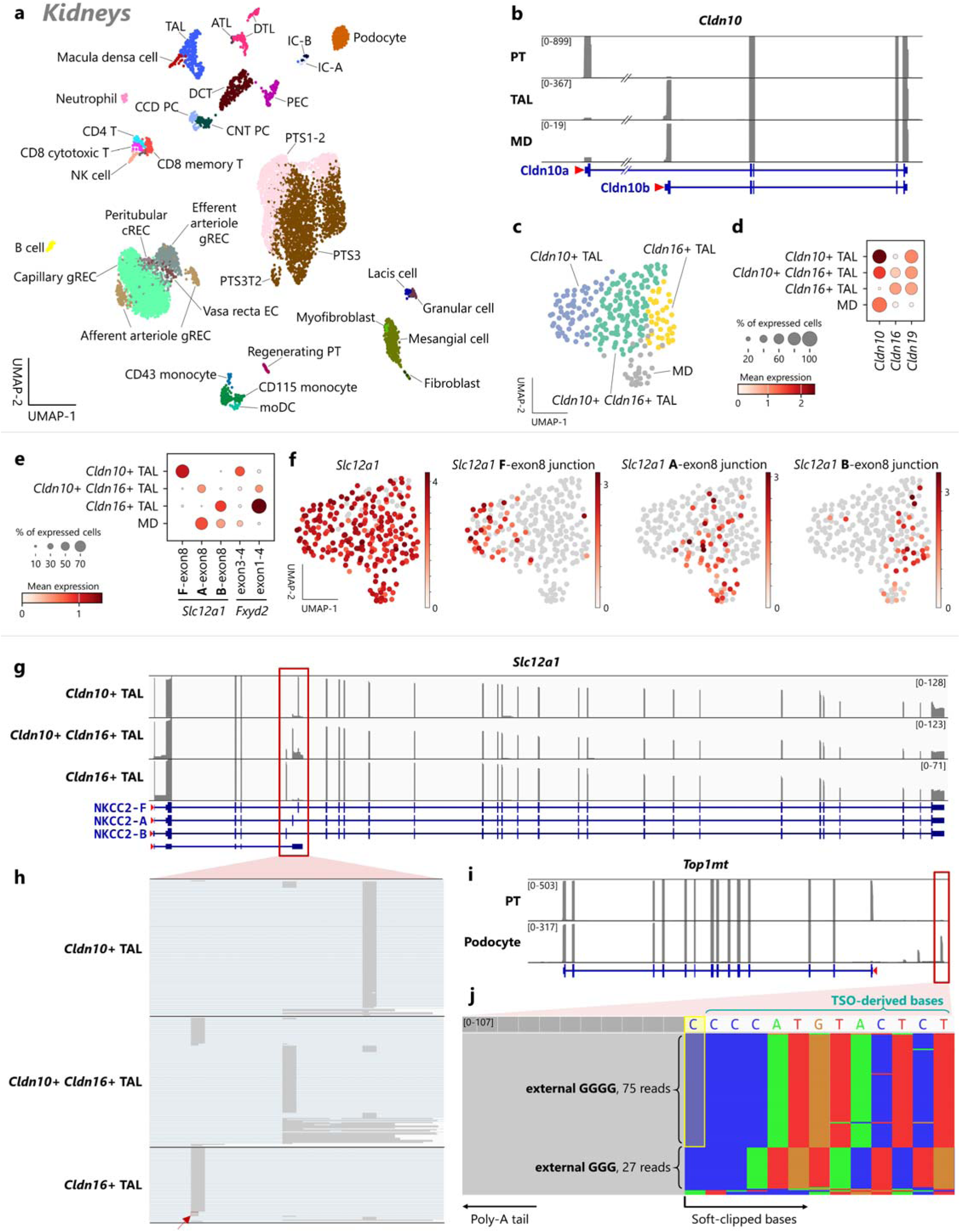
Ouro-Seq captures a high resolution of functional heterogeneity. **a**, Various cell types were identified in the kidneys. (Detailed cell type names can be found in **Extended Data Figure 10a.**) **b**, Alternative promoter usage of *Cldn10* between TAL and the proximal tubule. **c**, **d**, **e**, MD cells and three other distinct TAL cell populations were identified via subclustering of TAL cells (**c**, UMAP embedding). These TAL cell populations displayed distinct *Cldn* expression (**d**, dot plot) and differential transcript usage of *Slc12a1* and *Fxyd2* (**e**, dot plot), genes highly relevant to the biological functions of TAL. **f**, **g**, **h**, *Slc12a1* encodes three functionally distinct Na-K-Cl cotransporters (NKCC2-F, -A, and -B) through alternative exon inclusion of three cassette exons (F, A, and B), respectively. Except for macula densa cells, each TAL cell population predominantly expressed a unique NKCC2 isoform, visualized as differential splice junction usage (**f**, UMAP embeddings) and coverage patterns of *Slc12a1* transcripts (**g**, only the reads covering *Slc12a1* exon4 are shown for clarity) for the three TAL cell populations. **h**, Magnified view of the three cassette exons. The red arrow indicates a novel exon of *Slc12a1* utilizing an alternative 5’ splice site. **i**, Alternative promoter usage of *Top1mt* involving *de novo* promoters exclusively utilized by podocytes. **j**, Presence of an unencoded G (annotated with the yellow box) in cDNAs starting from one of the *de novo* promoters of *Top1mt*. MD, macula densa; PT, proximal tubule; TAL, thick ascending limb of the loop of Henle; TSO, template-switching-oligo.

### Discovery of novel promoters and genomic features

Besides the characterization of a proteome of a distinct cell population, Ouro-Seq can be utilized for more comprehensive functional characterization of known cell types. For example, we identified multiple novel isoforms of mitochondrial DNA topoisomerase 1 (*Top1mt*) exclusively expressed in podocytes or maturing sperm cells (**Figure 5i** and **Extended Data Figure 22a**), each of which utilizes a novel promoter region upstream of the known promoter. When the 5’ ends of sequencing reads utilizing the novel promoters were examined, the reverse transcription signature indicating the presence of the m7G cap at the 5’ end of an intact mRNA could be robustly identified for the majority of reads (∼70%), indicating the validity of the newly discovered promoters for *Top1mt* (**Figure 5j**). For podocytes and sperm cells, which have unique metabolic requirements to perform their biological functions, these novel *Top1mt* transcripts with altered protein-coding regions may generate specialized Top1mt isoforms that are likely essential for performing the specific functions of these cell types (**Supplementary Note 6**). Furthermore, we identified a novel antisense transcript of *Top1mt* that is specifically expressed in late spermatid cells, transcribed from a previously annotated promoter in the opposite direction (transcribed cis-regulatory element) (**Extended Data Figure 22b**). The 5’ ends of sequencing reads from the antisense transcript had the m7G cap signature at the 5’ end, as expected from previous studies^37^. The expression of such regulatory region-associated RNA may be responsible for altered promoter usage in these cells. Lastly, we identified various m7G-capped transposable element-derived transcripts in a locus-specific manner (**Extended Data Figure 23a**). For example, m7G-capped transcripts derived from an endogenous retroviral element (intracisternal A-particle^58^) were expressed in several cell types in the ovary and a few contractile cell types, including podocytes, specifically expressed in cells in the G0 phase and silenced in proliferating cells (**Extended Data Figure 23d**). Altogether, our frameworks allow sensitive identification of previously uncharacterized promoters (and novel transcribed cis-regulatory elements) utilized by rare cell types.

## DISCUSSION

Considering the molecular nature of reverse transcription and PCR amplification artifacts^18,20^, we developed an artifact depletion method without length bias via an intramolecular selective ligation reaction. Owing to its simplicity, this method can be applied to all single-cell (and spatial^19^) long-read sequencing libraries that utilize at least two different adapters at the ends of a molecule, including 3’ and 5’ barcoding-based scRNA-seq^59^, single-cell ATAC-seq^60^, single-cell multiome libraries, and spatial transcriptomics^19^, regardless of the single-cell capture platform (applicable to any scRNA-seq platform described in **Supplementary Note 1**) or long-read sequencing technology used. With the efficient depletion of barcoding artifacts and uninformative cDNA molecules, full-length transcriptomes of 10,000 single cells can be analyzed (∼5,000 reads per cell with an average read length >2 kbp) using a single PromethION flow cell (∼1,000 USD).

In Ouro-Seq, the selection processes of individual molecules are independent of each other (intramolecular interactions), whereas, in linear array assembly methods (such as HIT-scISO-Seq^24^, MAS-ISO-Seq^23^, and PB_FLIC-Seq^61^), the selection process is highly dependent on the overall library composition (intermolecular interactions). Linear array assembly methods do include selective ligation reactions that remove artifacts with invalid adapter combinations during the array assembly process^62^; however, because the assembly efficiency drops rapidly with increasing proportion of artifacts in the library, these methods require a separate artifact depletion step before the array assembly process can take place. In other words, the arrays of concatenated cDNAs cannot be assembled correctly in the presence of artifact molecules, necessitating the use of a separate artifact depletion method prior to array assembly (currently, the biotin-pulldown method with a strong bias toward shorter molecules).

We present the first reported mouse single-cell full-length transcriptomic atlas that characterized usage patterns of 217,046 unique splice junctions of 102,197 previously annotated transcripts across ∼200 cell types from 12 major organs and tissues. Overall, our isoform-resolved single-cell atlas not only provides a comprehensive view of complex cellular heterogeneity within an organism but also enables simultaneous functional annotation of both poorly characterized mRNA species and the cell populations expressing the transcripts. Additionally, our atlas could provide a valuable resource for studying the functional connections between transcription and splicing^63–65^ and the comprehensive modeling^66^ of the splicing process, including varying splicing patterns of mRNA transcripts across genes and expression patterns of splicing factors across cell types.

Currently, significant technical limitations exist for methods that do not rely on sequencing technologies for studying the expression of alternative mRNA transcripts at the resolution of individual cells. For instance, even though immunohistochemistry (IHC) can sensitively detect the protein product encoded by an mRNA transcript of interest, highly similar protein isoforms cannot be distinguished by the method if the region varying across the isoforms is inaccessible to the antibody (e.g., a transmembrane domain or an internal catalytic site). Our atlas revealed that the three distinct *Slc12a1* mRNA isoforms, encoding functionally distinct Na–K–Cl cotransporters (NKCC2-B, NKCC2-A, and NKCC2-F), are exclusively expressed by the three distinct cell populations in the thick ascending limb of the kidneys. Since the three mutually exclusive cassette exons of the *Slc12a1* isoforms encode a crucial transmembrane domain that determines the binding affinities for the substrate ions, studying expression patterns of NKCC2 isoforms across the nephron using IHC has been challenging. On the other hand, in situ hybridization (ISH) detects a 20–30 bp-long target-specific sequence in target mRNA transcripts, which can be virtually any region of mRNA species. ISH-based spatial transcriptomics technologies^67–69^ can overcome the limited number of target genes. However, if there are no unique splice junctions or exons exclusive to each alternative transcript, it is challenging to distinguish these isoforms using IHC- or ISH-based methods. Since various tropomyosin isoforms share exons, developing antibodies distinguishing specific tropomyosin isoforms has been difficult^70^, thus addressing Tpm1.8 and Tpm1.9 isoforms together. Our data revealed the exclusive usage of Tpm1.9 isoform by columnar and cuboidal epithelial cells over Tpm1.8 and vice versa for squamous epithelial cells. Also, these techniques cannot discover novel mRNA species utilizing *de novo* splice junctions, as pre-designed probes and antibodies are used for detection. Additionally, unlike long-read sequencing, identifying transcripts from pseudogenes and transposable elements is highly challenging using ISH-based methods because of the high sequence similarity between these genomic loci^71^.

Our analysis showed that the reverse transcription signatures of the m7G cap and enzymatically added poly(A) tail are highly effective for identifying full-length cDNAs from long-read scRNA-seq library laden with numerous RT and PCR artifacts. Even though we focused on the previously known reference transcriptome^27^, expression patterns of *de novo* mRNA transcripts have been captured, discovering novel promoters, splice sites, polyadenylation sites, and TE-derived transcripts. Additionally, through an integrative analysis of both mRNA species with and without m7G 5’ cap, insights into the cytoplasmic mRNA decapping, recapping, and degradation processes^72,73^ could be obtained at the single-cell level. Furthermore, it might be possible to develop a more accurate RNA velocity algorithm using the 5’ m7G cap signature; our atlas, containing differentiating spermatids, enterocytes, neurons, and other differentiating cells, could provide useful resources for developing such algorithms^40^. Lastly, as most aberrant transcripts (e.g., transcripts with retained introns and TE-derived transcripts) are eventually degraded by nuclear RNA exosome complex and cannot leave the nucleus^74^, the comparative analysis between single-nuclei and single-cells for the same cell type can reveal important insights into the dynamics and metabolisms^74^ of various aforementioned mRNA species.

Although single-cell multi-omics technologies have enabled groundbreaking research in the field of biomedical science, there have been limited studies on expression patterns of transcript isoforms, one of the critical layers of cellular information, due to technical limitations. Our long-read scRNA-seq technology and whole-organism single-cell splicing atlas will help advance the functional understanding of splicing isoforms and enhance our comprehension of various cell types.

## Supporting information

Supplementary Information for Mouse Single-Cell Long-Read Splicing Atlas by Ouro-Seq

## METHODS

### Single-cell cDNA library preparation and short-read sequencing of the library

First, single-cell suspensions were prepared for 10x Genomics 3’ v3.1 droplet-based sequencing (specific dissociation protocols for each sample are described in **Supplementary Methods**), targeting recovery of ∼8,000 cells (or nuclei); cell (or nucleus) concentration was determined using the Countess II Automated Cell Counter (Thermo Fisher Scientific). The pooled single-cell cDNA library was prepared following the manufacturer’s protocol (#CG000204 Rev B, 10x Genomics) with a few modifications. First, we increased the incubation time for reverse transcription reaction (GEM-RT reaction) from 45 minutes to 75 minutes to allow the completion of reverse transcription reactions for longer mRNA transcripts. Additionally, for the initial PCR amplification of the cDNA and all subsequent re-amplifications of the resulting cDNA library, the PCR elongation time was extended from 1 minute to 3 minutes to allow amplification of longer cDNA molecules (up to 6 kbp). We observed that increasing the extension time longer than 3 minutes occasionally results in a noticeable increase in the amount of 10x adapter dimers in the amplified library (and, accordingly, the reduced amount of the library). Additionally, we repeatedly observed that Ouro-SizeSelect robustly enriched cDNA molecules up to 6 kbp from the single-cell cDNA library that was initially amplified with the original extension time of 1 minute, implying that the increase of the extension time of the initial cDNA amplification step is not strictly necessary for characterizing (a subset of) longer cDNA molecules (up to 6 kbp). Next, the short-read scRNA-seq library was prepared using up to half of the amplified cDNA library, strictly following the manufacturer’s instructions (#CG000204 Rev B, 10x Genomics). The short-read scRNA-seq library was sequenced on a HiSeq platform (Illumina) (28 andℒ91 bases for read1 and read2, respectively), targeting 200M reads per sample (> 20,000 reads per cell). Subsequently, short-read scRNA-seq count matrices were generated from the base call files (BCLs) using Cell Ranger (v6.1.2, 10x Genomics) with the pre-built mouse reference (refdata-gex-mm10-2020-A, 10x Genomics).

### Nanopore long-read sequencing of the single-cell cDNA library using PromethION

The Nanopore sequencing library was prepared from a single-cell cDNA library using the Native Barcoding Kit 24 V14 kit (#SQK-NBD114.24, Oxford Nanopore Technologies) following the manufacturer’s protocol. The sequencing library was then loaded to a PromethION flow cell (R10.4.1) (# FLO-PRO114M, Oxford Nanopore Technologies) and sequenced on the PromethION 24 (Oxford Nanopore Technologies) with a sampling rate of 5 kHz. The raw sequencing output (FAST5 or POD5) was basecalled with Guppy (version 6.5.7) using the “SUP” basecalling model (the most accurate and slowest) “dna_r10.4.1_e8.2_400bps_5khz_sup.cfg”. The low-quality reads were filtered using the minimum quality score of 9.

For the construction of the Mouse Single-cell Long-Read Splicing Atlas, the single-cell cDNA libraries from the 22 samples were processed with Ouro-Deplete (for the depletion of cDNAs from mtDNA and rRNA) and subsequently, Ouro-SizeSelect (up to 6 kbp). The Ouro-Seq libraries were analyzed using 32 PromethION flow cells (∼1.5 flow cells per sample). In total, the sequencing runs generated ∼43 terabytes of the raw sequencing outputs (FAST5 and POD5 formats) and 3,388,782,353,700 base pairs of long-read sequencing data after filtering (**Supplementary Table 1**, the summarized result is shown in **Extended Data Table 1**).

### Ouro-Seq

#### Selective self-cyclization reaction

For implementing the selective self-cyclization of the cDNA molecules with valid 3’ and 5’ ends, a homology-based DNA assembly method (Gibson assembly) was utilized. First, 5–10 ng of a single-cell cDNA library is amplified for 3–5 cycles with a pair of primers that introduce 20 bp-long direct repeats at the ends of the cDNA molecules with valid 3’ and 5’ ends: “part-R1-Gibson” and “part-TSO-Gibson” for the 10x Genomics transcriptomics libraries and “BC0062-Gibson” and “BC0108-Gibson” for the SPLiT-Seq libraries (**Extended Data Table 3**). Next, 10-50 ng (up to 200 ng) of the resulting cDNA library is circularized using NEBuilder HiFi DNA Assembly Master Mix (#E2621S, New England Biolabs) with a total reaction volume of 10 µl following the manufacturer’s protocol. Using a higher concentration (> 200 ng/10 µl) of input cDNA molecules in the self-cyclization reaction mixture can decrease the self-cyclization efficiency, because the reaction rate of inter-molecular ligation reaction, which competes with intra-molecular ligation reaction that is solely responsible for self-cyclization reaction, albeit much smaller than that of the intra-molecular ligation reaction when the concentration is low, increases rapidly as the concentration increases, proportional to the square of the concentration. The resulting self-cyclization reaction mixture that contains (self-ligated) circular cDNA molecules is then subjected to different downstream protocols according to specific applications.

#### Reversal reaction of self-cyclization

After the selective self-cyclization reaction, circular cDNA molecules become resistant to the action of exonucleases, allowing depletion of cell barcode-free and cell barcode-doublet artifacts with exonuclease treatment. However, circular DNA molecules are unsuitable for PCR amplification (due to the difficulties of complete denaturation) or long-read sequencing experiments (due to no ends available for sequencing adapter ligation). For downstream applications, the circular cDNA molecules can be efficiently linearized with CRISPR-Cas9 nuclease, reversing the self-cyclization process (hence, the reversal reaction of self-cyclization). First, Cas9-sgRNA ribonucleoprotein complexes (RNPs) are assembled by mixing the following reagents and incubating the mixture at room temperature for 10 minutes: 7 µl of nuclease-free water, 9 µl of 10 ng/µl sgRNA targeting the intra-molecular ligation junction (synthesized using the “sgRNA.Gibson-Junc” oligo in **Extended Data Table 3** by following the protocol described in **Supplementary Methods**), 2 µl of 10x NEBuffer r3.1 (#B6003S, New England Biolabs), and 2 µl of 1 µM SpCas9 nuclease (#M0386S, New England Biolabs). Subsequently, the assembled Cas9-sgRNA RNPs (20 µl) and 1 µl of 10x NEBuffer r3.1 are added to 9 µl of purified circular DNA library dissolved in nuclease-free water. The linearization reaction mixture (30 µl) is then incubated at 37°C for 25 minutes to digest the self-ligation junctions of the circular cDNA molecules. For cleanup, 1 µl of Proteinase K (#P8107S, New England Biolabs) is added to the linearization reaction mixture, which is incubated at room temperature for 10 minutes. Finally, the linearization reaction mixture is purified with 1.2x SPRIselect beads (#B23318, Beckman Coulter) to yield linear cDNA molecules with valid 3’ and 5’ ends.

#### Comprehensive artifacts depletion with Ouro-Seq

For standard depletion of cell barcode-free and cell barcode-doublet artifacts without any additional targeted depletion or enrichment operations of target cDNA molecules (for the descriptions of Ouro-Deplete and Ouro-Enrich methods, see **Supplementary Methods**), the self-cyclization reaction mixture is directly subjected to exonuclease treatment and subsequently, the reversal reaction of self-cyclization. First, a mixture of exonucleases, 10 units of Exonuclease III (#M0206S, New England Biolabs), 0.5 units of Lambda Exonuclease (#M0262S, New England Biolabs), and 20 units of Thermolabile Exonuclease I (#M0568S, New England Biolabs), is added to the self-cyclization reaction mixture on ice. Next, the reaction mixture is incubated at 37 °C for 20 minutes. The exonucleases are heat-inactivated by incubating the reaction mixture at 70 °C for 10 minutes. The self-cyclization reaction mixture is purified with 1.2x SPRIselect beads and resuspended in 9 µl of nuclease-free water, yielding a circular cDNA library, depleted of cell barcode-free and cell barcode-doublet artifacts. After the self-cyclization reversal reaction, a linear cDNA library can be obtained, which can be directly analyzed with long-read sequencing or re-amplified with PCR for downstream applications.

#### Robust size-selection with Ouro-SizeSelect

The first step of Ouro-SizeSelect is the depletion of cDNA molecules smaller than 0.9 kbp using 0.5x SPRIselect beads. Subsequently, sequential size-selection steps using 0.9x, 0.8x, 0.75x, 0.725x, 0.7x, 0.69x, and 0.68x size-selection solutions with size cutoffs at 1.25, 1.75, 2.25, 2.75, 3.25, 3.75, and 4.5 kbp are performed, respectively (for the full description of the method, see **Supplementary Methods**). For each size-selection step, the resulting library can be either directly subjected to the next size-selection step or optionally re-amplified using PCR (**Supplementary Methods**). After all the sequential size-selection steps of Ouro-SizeSelect are completed, the size-selection results are evaluated using a gel-electrophoresis-based platform, such as the Agilent Bioanalyzer 2100 system (Agilent). Finally, for a long-read sequencing experiment, the size-selected libraries are pooled with the following weight ratio (the numbers are in parentheses, followed by the expected size range): the original artifact-depleted library (10 ng, 0.25-1.5 kbp), the size-selected libraries after 0.5x SPRIselect (40 ng, 0.75-2 kbp), 0.9x (40 ng, 1.25-2.5 kbp), 0.8x (40 ng, 1.75-3 kbp), 0.75x (20 ng, 2.25-3.5 kbp), 0.725x (20 ng, 2.75-4 kbp), 0.7x (20 ng, 3.25-4.5 kbp), 0.69x (20 ng, 3.75-5.25 kbp), and 0.68x (20 ng, 4.5-6 kbp) size-select solutions (**Extended Data Figure 1c**).

### Computational analysis

#### Comprehensive analysis of full-length transcriptome at single-cell level with Ouro-Tools

For the construction of the mouse single-cell long-read splicing atlas (v202405) and the benchmarking of previously published long-read scRNA-seq datasets, the Ouro-Tools (v0.1.1) pipeline has been utilized (for the full description of the pipeline, see **Supplementary Methods**). The Ouro-Tools pipeline comprises five main modules (**Supplementary Figure 1**), as briefly described below. The raw long-read pre-processing module (**Supplementary Figure 2**) has a dual function for (1) providing comprehensive quality control metrics of a long-read scRNA-seq experiment (e.g., **Extended Data Figure 2-3**) and (2) pre-processing of raw long-read sequencing data for the downstream analysis. The raw long-read pre-processing module efficiently identifies and excludes the following molecules from the subsequent analysis: cell barcode-free and cell barcode-doublet artifacts, chimeric cDNA molecules, molecules generated by mispriming events, and uninformative cDNA molecules (e.g., mtDNA-derived and rRNA-derived cDNA molecules); each valid cDNA molecule remained after the filtering is re-oriented so that the poly(A) tail is placed at the 3’ end, generating a strand-specific long-read scRNA-seq data. Furthermore, for a long-read scRNA-seq experiment that utilized Ouro-Enrich for targeted enrichment of the transcripts of interest, the module searches for the sign of a topological modification introduced by Ouro-Enrich (after an Ouro-Enrich experiment, an inverted transposition occurs around the target site, see **Figure 2a**), reconstructing the original cDNA from the read. The resulting strand-specific cDNA reads are aligned to the reference genome, guided by the known splice junctions from gene annotations. Next, the barcode extraction module (**Supplementary Figure 3**) extracts and corrects cell barcode (CB) and unique molecular identifier (UMI) sequences from the soft-clipped sequence of each read, exporting the results as a “barcoded” BAM file (**Supplementary Figure 3**). Specifically, for the correction of UMI sequences, the raw UMIs are grouped for each barcode attachment site (i.e., polyadenylation site) on the reference genome for each corrected CB and clustered using a linear-time UMI clustering algorithm (**Supplementary Methods**). Subsequently, the biological full-length identification module classifies cDNAs based on the presence of the reverse transcription signatures of the m7G cap (valid 5’ end) and the enzymatically added poly(A) tail (valid 3’ end), identifying the cDNA molecules representing full-length mRNAs. Consequently, the cDNA molecules representing fragmented mRNA transcripts are excluded from the downstream analysis. Next, the size distribution normalization module removes the effects of highly abundant mRNA transcripts specific to certain cell types and experimental conditions to estimate each sample’s underlying mRNA size distributions. Subsequently, the module adjusts the mRNA size distribution of each sample to the reference mRNA size distribution to reduce the technical variability originating from the differences in the read length distributions between samples. Lastly, the single-cell long-read count module quantifies isoforms by utilizing the reverse transcription signatures of the 3’ and 5’ ends of full-length mRNA molecules (i.e., the TSS and TES matching algorithm, see **Supplementary Methods**), while simultaneously reducing unwanted technical variations in read length distributions for accurate integration of multiple long-read scRNA-seq datasets (**Figure 3a**). Specifically, the module normalizes the mRNA size distribution of every cell in a sample using global scale factors calculated for the sample (i.e., the optimal correction ratios that were calculated by the previous module), before applying the size factor normalization method for individual cells in the sample. The resulting size-distribution-normalized single-cell long-read count matrix is exported as a feature-barcode matrix (MEX) format, compatible with major single-cell analysis platforms.

#### Construction of the Mouse Single-Cell Long-Read Splicing Atlas

Following the previous studies^19–21,26,76^, the short-read and long-read scRNA-seq datasets were integrated by clustering and annotating the cells using the short-read scRNA-seq datasets and subsequently overlaying each cell with the long-read scRNA-seq data. Particularly, this strategy was selected since background noise removal of the raw single-cell count matrix, including the removal of ambient RNA, can only be performed using the algorithms developed for short-read scRNA-seq data. Additionally, due to the higher sparsity of long-read scRNA-seq, we primarily utilized short-read scRNA-seq for more accurate clustering and annotation of the cells. The following steps were utilized to process the short-read scRNA-seq datasets. First, for each sample, background noise-removal (for remedying ambient RNA) was performed using cellbender^77^ using the parameters expected-cells=5000 and total-droplets-included=60000. The denoised single-cell count matrices were further analyzed using Scanpy^78^. The cells were initially filtered using the parameters min_gene=300 and min_count=550. The counts were normalized using the size factor normalization with the total count of 10,000 as the target, and the resulting normalized counts were log-transformed. Initial doublet detection and removal were performed using Scrublet^79^ with the expected doublet rate of 0.15 and the sample names as the batch key; cells with doublet scores larger than 0.5 were considered doublets and discarded in the downstream analysis. For each organ or tissue, highly variable genes were selected using the parameters min_mean=0.0125, max_mean=3, min_disp= −0.25, and flavor=“seurat”; dimensionality reduction was performed using Principal Component Analysis (PCA), and the resulting top 100 PCs were stored in the AnnData object. For the global analysis of the atlas, all detected genes were selected for dimensionality reduction with PCA; the resulting top 200 PCs were stored in the AnnData object. Next, batch effects in the stored PCs were removed using the Harmony^80^ algorithm with the maximum number of iterations of 20 and the sample names as the batch key. For each organ and the entire atlas, the batch-effect-corrected top 100 and 200 PCs were subsequently utilized for constructing the neighborhood graphs of the single cells, with 15 and 125 as the numbers of neighbors, respectively. Lastly, uniform manifold approximation and projection (UMAP) embeddings of the single cells were constructed using the neighborhood graph.

The single cells were analyzed at the organ level for initial clustering and annotations^12,81^. For each organ or tissue, using the batch-effect-corrected PCs, single cells were initially clustered using the leiden^82^ algorithm, setting the resolution parameter to 1. Next, the clustering results were visualized on the UMAP embeddings, and the groups of the clusters with inaccurate cluster boundaries were combined and subclustered to refine the clustering results. During the subclustering process, a group of biologically similar clusters without exclusive marker gene(s) were combined. Simultaneously, the clusters of low-quality cells and doublets without unique marker gene(s) are identified, respectively, and excluded from the downstream analysis. Once the subclustering process was completed, the resulting clusters were annotated using the structured vocabulary of the Cell Ontology^83,84^ (CL) to facilitate^43^ the exploration of our single-cell full-length transcriptome atlas, totaling 215 cell types across 12 organs and tissues. As previously observed^12,75^, after clustering and visualizing all cells of the atlas with UMAP (**Figure 4b**), we observed moderate to high organ-specificity for the shared cell types that are resident for various tissues and organs, such as fibroblasts, macrophages, smooth muscle cells, and endothelial cells. In contrast, we found that most cells of the circulating immune cell types, such as B, T, and NK cells, are shared across the organs.

Subsequently, we utilized the standard Ouro-Tools pipeline (v0.1.1) to process the 3.4 Tbp of Nanopore long-read sequencing data generated from the 22 Ouro-Seq libraries (see **Supplementary Methods** for a detailed description). For the identification of full-length cDNAs, valid TSSs are identified using all 22 samples collectively. The normalization of mRNA size distributions was performed by constructing the reference mRNA size distribution using all 22 samples. The transcripts derived from the transposable elements and the regulatory elements were quantified at the resolution of individual cells based on the annotations from the RepeatMasker resource^85,86^ (in the format available in the UCSC Table Browser^87^) and the Ensembl Regulatory Build^88^, respectively. Lastly, the barcoded BAM files of the atlas were split into BAM files of individual cells for the downstream analysis.

#### Benchmarking of single-cell long-read methods

We compare the performance of Ouro-Seq with other single-cell long-read RNA-Seq methods including FLOUR-seq^40^ (utilizing a microwell-based scRNA-seq method), ScISOr-Seq^18,19,76^, SnISOr-Seq^21^, FlsnRNA-seq^89^, FLT-seq^46,90^, scCOLOR-Seq^91^, and Ouro-Seq (current study) using the standard Ouro-Tools pipeline (v0.1.1) (see **Supplementary Methods** for a detailed description). First, the raw FASTQ files of the 133 sequencing runs from the previous 13 studies^18–21,26,40,46,76,89–93^ were downloaded from the Sequence Read Archive (SRA). Of note, the sequencing runs that contained the unprocessed reads of the concatenated cDNA arrays (e.g., unprocessed sequencing reads from MAS-ISO-seq^23^ and HIT-scISOseq^24^ experiments) were discarded since these reads need alternative preprocessing pipelines, and the same types of summary metrics of individual cDNA molecules could not be obtained using the current version (v0.1.1) of the raw preprocessing module of Ouro-Tools, which is currently optimized for the detection of the chimeric cDNA molecules commonly observed in Nanopore sequencing experiments. Next, each group of sequencing runs of the same sample is aggregated into a single FASTQ file, resulting in 40 datasets, totaling 2,292,319,826,251 base pairs (2.3 Tbp) of long-read sequencing data (the total number of reads is 1,907,986,081, with an average read length of 1,201 bp). Subsequently, the raw long-read pre-processing module was utilized to retrieve the QC metrics of each long-read scRNA-seq method. During the process, we detected the datasets that follow the output format of the SiCeLoRe pipeline^20^; all reads are strand-specific reads that contain poly(A) tails at their 3’ ends. Consequently, a study^20^ “Lebrigand2020” containing these samples was dropped from the comparison since the summary statistics of the raw sequencing reads of cDNA molecules cannot be obtained from the output from the SiCeLoRe pipeline. However, the complete benchmarking result, including the samples from the “Lebrigand2020” study, can be found in **Supplementary Table 2**; the summarized result is shown in **Extended Data Table 2**.

## DATA AVAILABILITY

All processed data, including raw short-read and long-read sequence data of our atlas, have been deposited in the Korea Sequence Read Archive^94^ (KRA, publicly accessible at https://kbds.re.kr/KRA) under accession KAP240772.

## CODE AVAILABILITY

Ouro-Tools is available at https://github.com/ahs2202/ouro-tools/releases/tag/v0.1.1, along with tutorials of the Ouro-Tools pipeline, complete with example Ouro-Seq datasets. All original code has been deposited at Zenodo (https://doi.org/10.5281/zenodo.13334958) and will be publicly available as of the date of publication.

## ACKNOWLEDGEMENTS

J.P. acknowledges that this research was supported by the Samsung Science and Technology Foundation (SSTF-BA2001-
11 to J.P.) and the National Research Foundation (NRF) of Korea funded by the Ministry of Science and ICT (MSIT)
(RS-2024-00335026). We thank the Genomic Technologies Group at the Garvan Institute of Medical Research for
providing excellent long-read sequencing services that generated a part of the long-read scRNA-5 seq data of our atlas. We thank our colleagues Seo-Gyeong Bae and Gyeong-Dae Kim for providing technical support during the analysis of previously published short-read scRNA-seq data.

## AUTHOR CONTRIBUTIONS

J.P. and S.S. conceptualized the standardized high-throughput long-read scRNA-seq method. H.A., J.P., and S.C. developed the core ideas of Ouro-Seq. S.C., H.A., and J.P. optimized the Ouro-Seq protocols. S.C., H.A., S.S., J.P., and J.Y. optimized the tissue dissociation protocols. S.C. and S.S. prepared the scRNA-seq libraries. S.C., S.O., and H.A. performed long-read sequencing. J.Y. and H.A. analyzed scRNA-seq data and compiled the cell type annotations of the atlas. H.A., C.L., J.P., and R.L. developed the key concepts of Ouro-Tools. H.A. implemented the Ouro-Tools. H.A. analyzed long-read scRNA-seq data of the atlas. Lastly, H.A., J.P., and S.C. compiled the results and wrote the manuscript.

## DECLARATION OF INTERESTS

J.P., H.A., and S.C. are inventors on a patent application related to this work filed by Gwangju Institute of Science and Technology.

## SUPPLEMENTARY INFORMATION

Supplementary Note 1: Limitations of single-cell full-length transcriptome analysis using short-read sequencing alone.

Supplementary Note 2: Unintentional amplification of cell barcode-doublet artifacts by the asymmetric PCR-based artifact depletion methods.

Supplementary Note 3: Diverse transcript usage patterns of *Tpm1* across cell types.

Supplementary Note 4: Previous observations indicating cellular heterogeneity in the thick ascending limb of the loop of Henle.

Supplementary Note 5: An improved model for explaining the location-specific formation of mosaic tight junctions in the thick ascending limb of the loop of Henle.

Supplementary Note 6: *De novo* promoters of *Top1mt*.

Supplementary Methods

Supplementary Table 1: Comprehensive summary of the 22 Ouro-Seq-applied long-read scRNA-seq datasets for the Mouse Single-Cell Long-Read Splicing Atlas.

Supplementary Table 2: Comprehensive summary of benchmarking results of various long-read scRNA-seq methods.

Supplementary Table 3: List of DNA oligo sequences utilized for sgRNA synthesis for Ouro-Deplete and Ouro-Enrich (Ouro-Cell-Enrich) experiments.

Supplementary Table 4: Proportions of long-reads representing full-length cDNAs across genes for the Ouro-Seq libraries prepared from *Mus musculus*.

Supplementary Table 5: Numbers of detected genes, isoforms, splice junctions, exons, transcribed regulatory elements, and transcribed transposable elements for each cell type.

Supplementary Table 6. The list of SAM tags utilized by Ouro-Tools.

Supplementary Table 7. The list of bitwise flags utilized by the single-cell long-read count module of Ouro-Tools for indicating the classification results of individual reads.

Supplementary Figure 1. An overview of the Ouro-Tools pipeline.

Supplementary Figure 2. A flowchart representing the “LongFilterNSplit” workflow of the Ouro-Tools pipeline.

Supplementary Figure 3. A flowchart representing the “LongExtractBarcodeFromBAM” workflow of the Ouro-Tools pipeline.

Supplementary Figure 4. A flowchart representing the biological full-length identification module of the Ouro-Tools pipeline.

Supplementary Figure 5. A flowchart representing the “LongCreateReferenceSizeDistribution” workflow of the Ouro-Tools pipeline.

Supplementary Figure 6. A flowchart representing the “LongExportNormalizedCountMatrix” workflow of the Ouro-Tools pipeline.

## EXTENDED DATA TABLES AND FIGURES

**Extended Data Table 1.** Summary of the 22 Ouro-Seq-applied long-read scRNA-seq datasets for the Mouse Single-Cell Long-Read Splicing Atlas.

| Sample name | total number of base pairs (bp) | total number of base pairs for reads with poly(A) (bp) | total number of base pairs for reads without mispriming events (bp) | total number of base pairs for reads without cell barcode (bp) | total number of base pairs aligned to unwanted sequences (bp) |
| --- | --- | --- | --- | --- | --- |
| mBrain | 193,362,031,853 | 150,138,632,264 (78%) | 74,732,568,447 (39%) | 13,936,693,505 (7%) | 461,929,678 (0%) |
| mBrain Nuclei | 144,137,779,859 | 110,293,773,668 (77%) | 59,072,199,072 (41%) | 13,453,368,821 (9%) | 991,689,153 (1%) |
| mSkin | 165,157,058,835 | 135,452,970,515 (82%) | 79,194,082,224 (48%) | 11,961,469,139 (7%) | 2,122,723,153 (1%) |
| mSkin2 | 149,981,379,714 | 126,824,130,533 (85%) | 70,609,469,431 (47%) | 7,177,761,403 (5%) | 2,308,658,875 (2%) |
| mHeart | 144,242,278,439 | 126,593,478,679 (88%) | 68,931,210,027 (48%) | 10,851,825,096 (8%) | 2,054,495,307 (1%) |
| mHeart Nuclei | 172,702,337,387 | 131,262,472,932 (76%) | 46,627,621,468 (27%) | 17,328,574,372 (10%) | 5,869,544,466 (3%) |
| mIntestine | 145,078,058,094 | 114,064,182,450 (79%) | 58,614,572,103 (40%) | 11,478,805,086 (8%) | 761,656,610 (1%) |
| mIntestine2 | 146,279,677,415 | 121,087,072,171 (83%) | 72,523,024,552 (50%) | 9,518,290,159 (7%) | 3,781,752,974 (3%) |
| mKidney | 122,137,627,629 | 106,440,883,741 (87%) | 63,079,904,703 (52%) | 9,343,189,261 (8%) | 2,204,860,985 (2%) |
| mKidney Glomerulus | 151,192,452,761 | 122,556,980,882 (81%) | 65,770,637,008 (44%) | 11,792,278,685 (8%) | 7,078,041,510 (5%) |
| mLiver | 173,485,621,195 | 142,883,445,092 (82%) | 64,147,358,723 (37%) | 18,884,099,082 (11%) | 1,744,121,395 (1%) |
| mLiver2 | 129,713,577,985 | 105,528,422,300 (81%) | 56,667,733,815 (44%) | 6,307,845,727 (5%) | 6,640,156,485 (5%) |
| mLiver Nuclei | 79,038,580,122 | 57,515,645,244 (73%) | 29,055,160,936 (37%) | 7,459,871,040 (9%) | 7,533,511,853 (10%) |
| mLung | 129,698,533,289 | 104,640,985,629 (81%) | 49,033,733,653 (38%) | 10,416,623,578 (8%) | 770,468,259 (1%) |
| mLung2 | 218,821,390,150 | 181,022,629,439 (83%) | 87,636,847,485 (40%) | 17,932,634,796 (8%) | 1,167,630,778 (1%) |
| mLung3 | 147,641,528,068 | 127,231,202,651 (86%) | 66,390,940,687 (45%) | 8,688,830,411 (6%) | 2,051,871,604 (1%) |
| mOvary | 163,871,026,494 | 133,286,552,081 (81%) | 59,692,397,865 (36%) | 12,073,872,685 (7%) | 1,415,648,296 (1%) |
| mPBMC | 155,916,992,131 | 122,676,456,834 (79%) | 54,343,546,229 (35%) | 13,978,742,033 (9%) | 1,181,394,519 (1%) |
| mPancreas | 151,741,488,732 | 132,193,277,318 (87%) | 72,752,312,336 (48%) | 7,750,349,742 (5%) | 2,718,553,601 (2%) |
| mPancreas Nuclei | 193,641,982,025 | 142,271,356,425 (73%) | 59,112,082,031 (31%) | 16,668,934,216 (9%) | 2,787,240,126 (1%) |
| mTestis | 161,082,419,471 | 140,351,458,004 (87%) | 99,104,790,230 (62%) | 11,054,383,573 (7%) | 180,057,907 (0%) |
| mThymus | 149,858,532,052 | 123,422,076,424 (82%) | 56,564,039,161 (38%) | 13,807,309,890 (9%) | 243,922,020 (0%) |
| *Total* | **3,388,782,353,700** |  |  |  |  |

**Extended Data Table 2.**
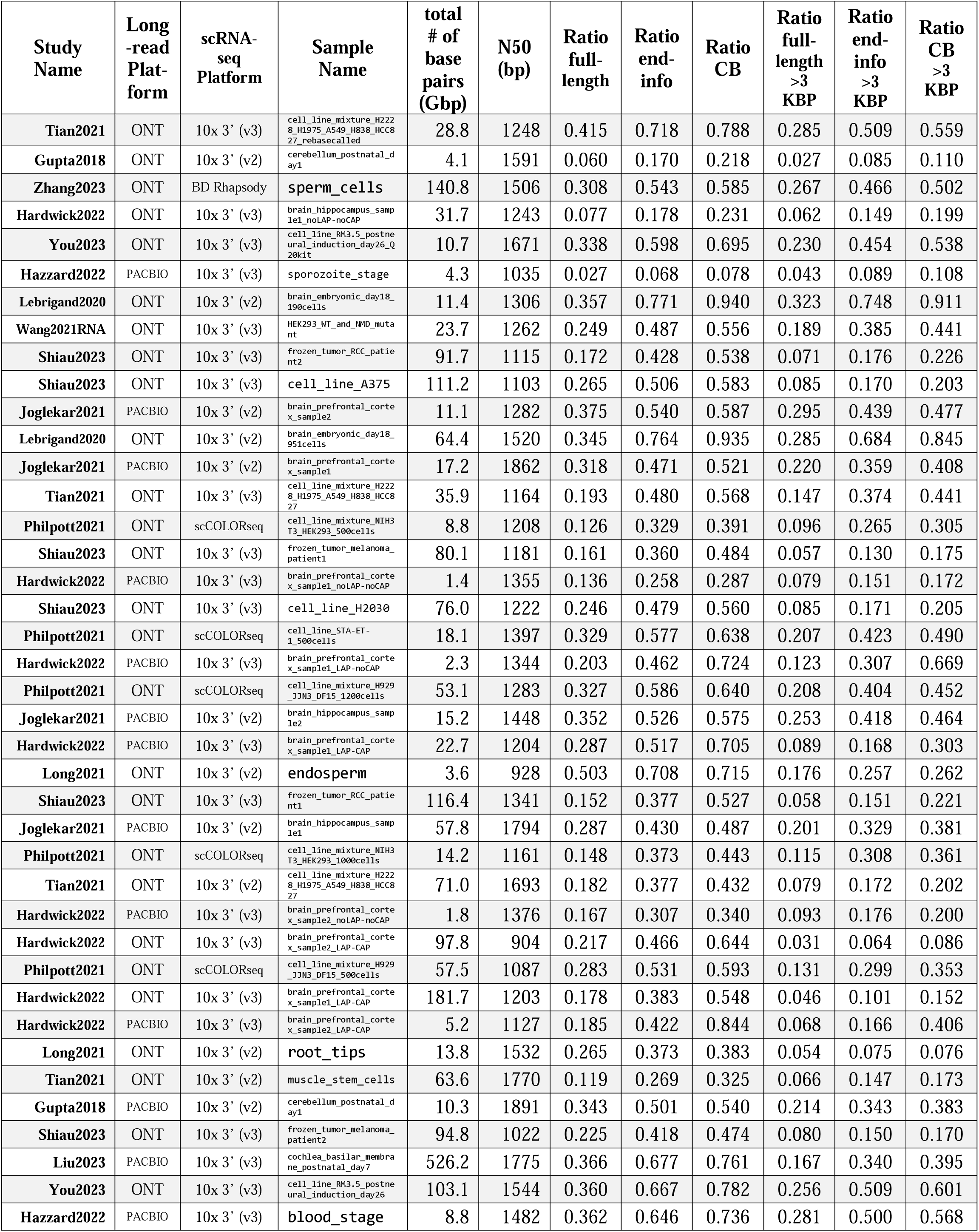
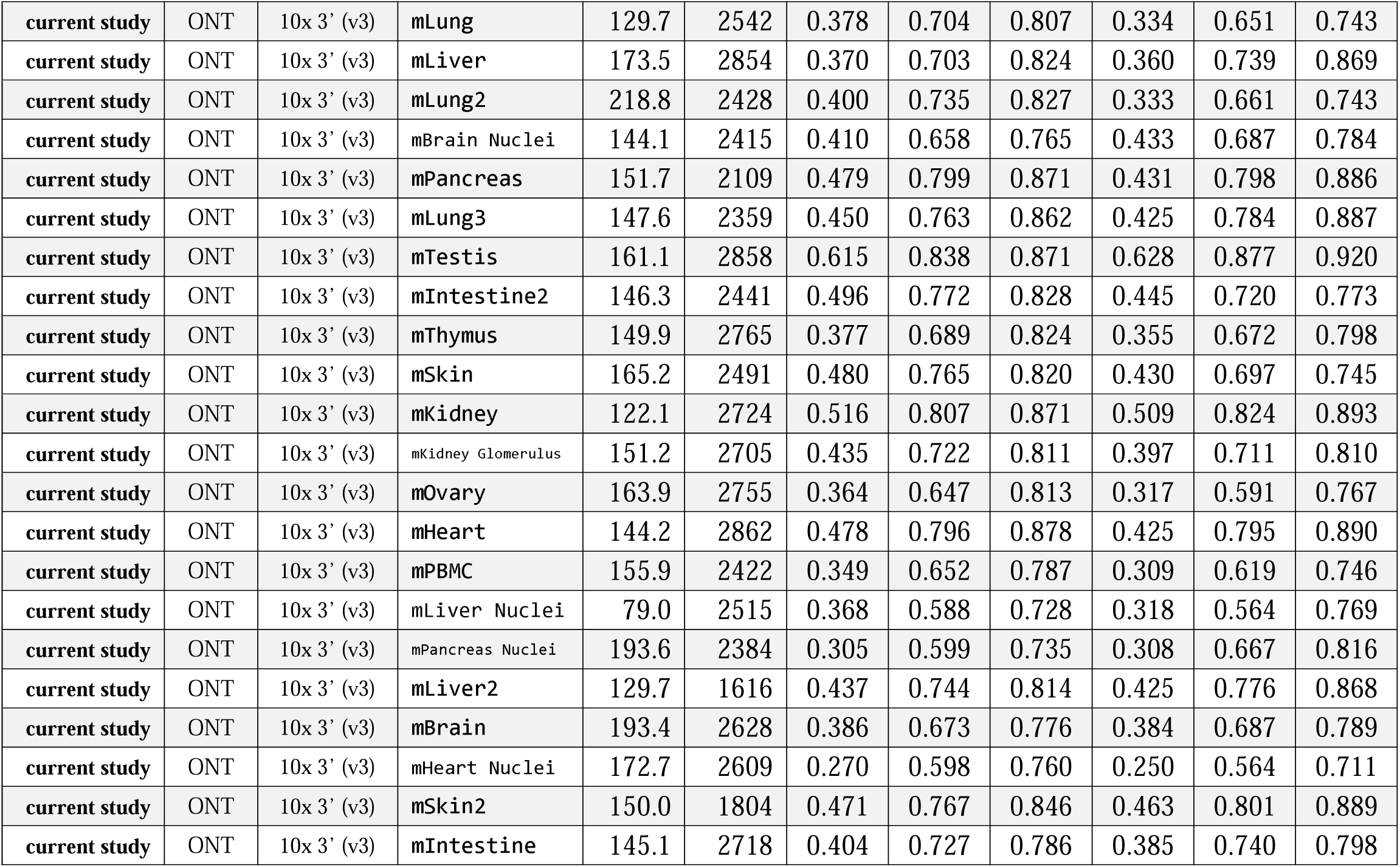
Summary of benchmarking results of various long-read scRNA-seq methods.

**Extended Data Table 3.**
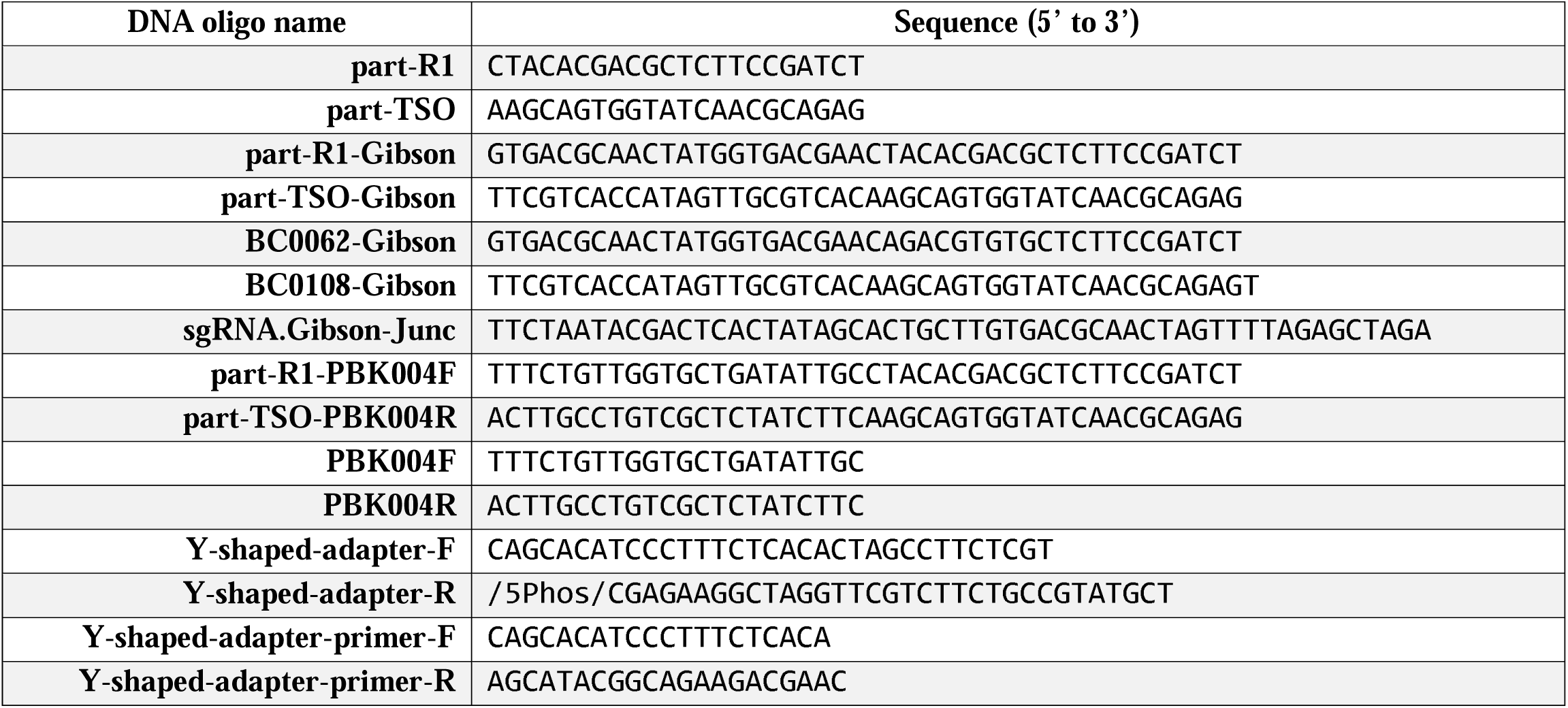
List of DNA oligo sequences utilized for Ouro-Seq experiments.

**Extended Data Figure 1.**
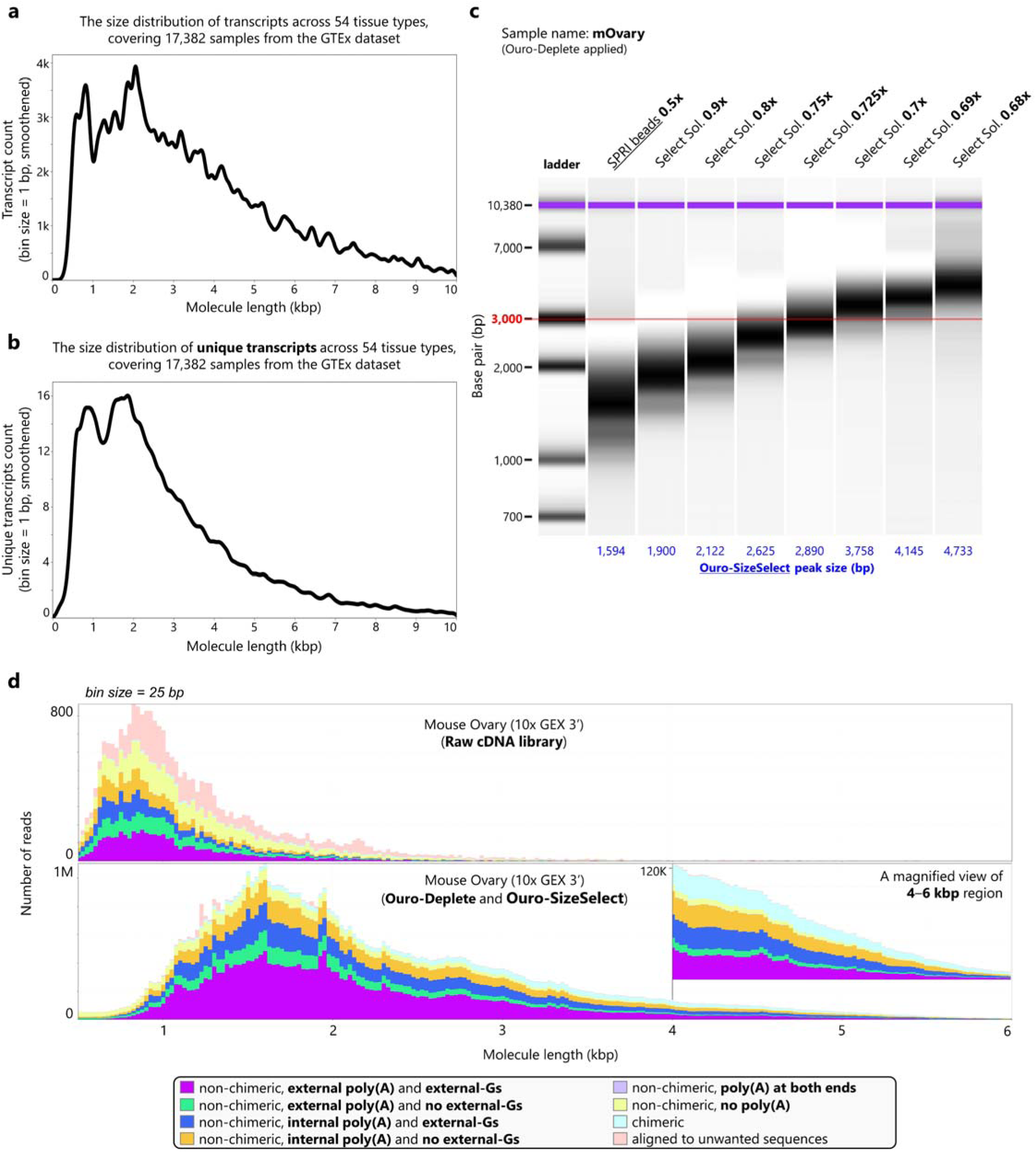
Robust enrichment of longer cDNAs from long-read scRNA-seq library using Ouro-SizeSelect. **a**, The mRNA size distribution of all the 17,382 short-read RNA-seq samples from the GTEx was reconstructed using the transcript counts of RNA-seq experiments. **b**, The size distribution of unique mRNA transcripts detected in all the 17,382 samples from the GTEx. **c**, A series of sequential size selection steps are employed in Ouro-SizeSelect for robust enrichment of longer cDNAs from a pooled long-read scRNA-seq library. **d**, A raw single-cell cDNA library from the mouse ovary (above) was processed with Ouro-Deplete and Ouro-SizeSelect (below), resulting in efficient characterization of longer full-length cDNAs (>2 kbp). Colors represent different classes of cDNA molecules. cDNAs without signatures of the mispriming events (“non-chimeric, external poly(A) and external-Gs”) are colored as violet.

**Extended Data Figure 2.**
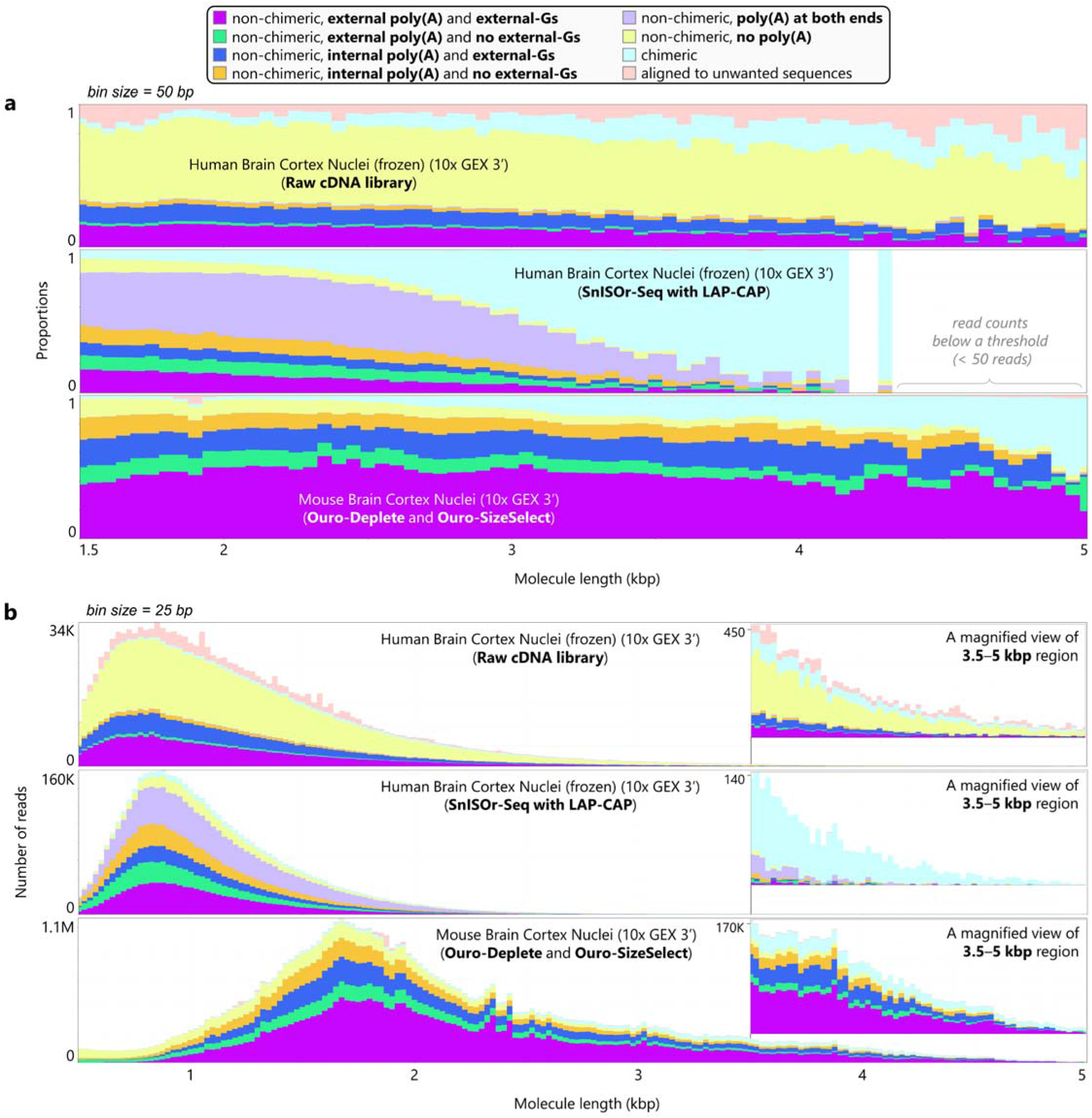
Comparison between the asymmetric PCR-based artifact removal method and Ouro-Seq. **a**, **b**, Ouro-Seq efficiently depletes both types of barcoding artifacts, cell barcode-free artifact (“non-chimeric, no poly(A)”) and cell barcode-doublet artifact (“non-chimeric, poly(A) at both ends”), without significant length bias. Colors represent different categories of cDNAs. The proportion of *in vitro* full-length cDNA molecules (cDNAs without mispriming signatures, “non-chimeric, external poly(A) and external-Gs”) is largely invariant in the raw single-cell cDNA library (top) and long-read scRNA-seq library processed by Ouro-Seq (bottom) across a wide range of cDNA sizes. However, in the long-read scRNA-seq library processed by the asymmetric PCR-based artifact removal method, the proportion of *in vitro* full-length cDNAs decreased as the cDNA size increased, indicating a strong length bias of the artifact removal method. **a**, Proportions of each cDNA category before (top, sample# “brain_prefrontal_cortex_sample2_noLAP-noCAP,” **Extended Data Table 2**) or after (middle, sample# “brain_prefrontal_cortex_sample2_LAP-CAP”) applying the asymmetric PCR-based artifact removal method (SnISOr-Seq); proportions of each cDNA category after applying Ouro-Seq (Ouro-Deplete with Ouro-SizeSelect) (bottom, sample# “mBrain Nuclei”). **b**, Read counts of each cDNA category for the same samples.

**Extended Data Figure 3.**
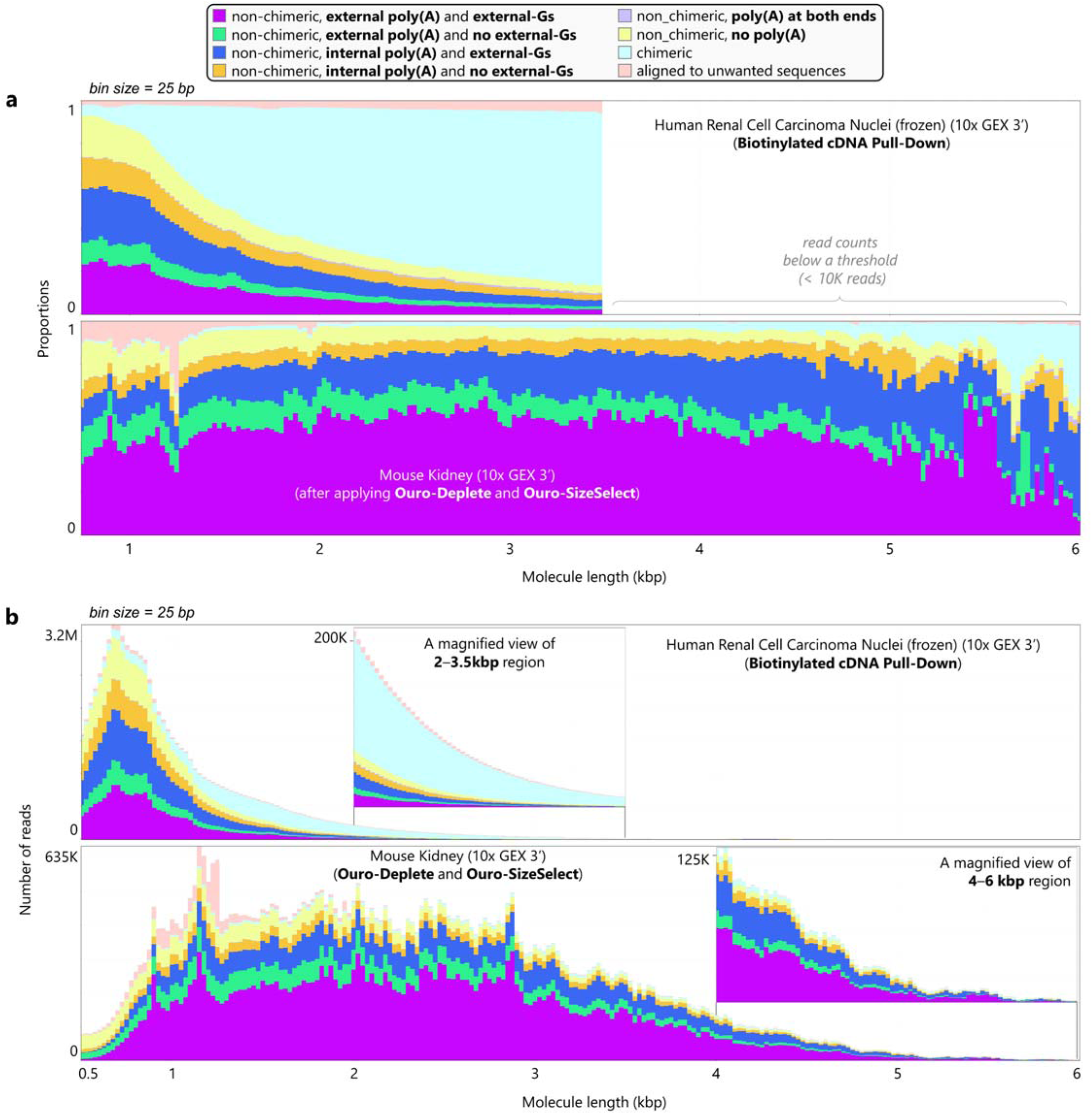
Comparison between the biotin selection-based artifact removal method and Ouro-Seq. **a**, **b**, Ouro-Seq enables the characterization of a wide range of full-length mRNA transcripts at the resolution of individual cells. Colors represent different categories of cDNAs. The proportion of *in vitro* full-length cDNA molecules (cDNAs without mispriming signatures, “non-chimeric, external poly(A) and external-Gs,” colored as violet) is largely invariant in the long-read scRNA-seq library processed by Ouro-Seq (below) across the entire size range covered by Ouro-Seq. However, in the long-read scRNA-seq library processed by the biotin selection-based artifact removal method, the proportion of *in vitro* full-length cDNAs decreased as the cDNA size increased. Additionally, the biotin selection-based artifact removal method was not effective at removing the cell barcode-doublet artifact (“non-chimeric, poly(A) at both ends,” colored as a light shade of purple), accounting for 3%–5% of barcoded cDNA larger than 2 kbp. **a**, Proportions of each cDNA category after applying the biotin selection-based artifact removal method (above, sample# “frozen_tumor_RCC_patient2,” **Extended Data Table 2**) or after applying Ouro-Seq (Ouro-Deplete with Ouro-SizeSelect) (below, sample# “mKidney”). **b**, Read counts of each cDNA category for the same samples. The read length distribution of the long-read scRNA-seq library processed by the biotin selection method indicates that the artifact removal process is strongly biased toward shorter cDNAs (<1 kbp).

**Extended Data Figure 4.**
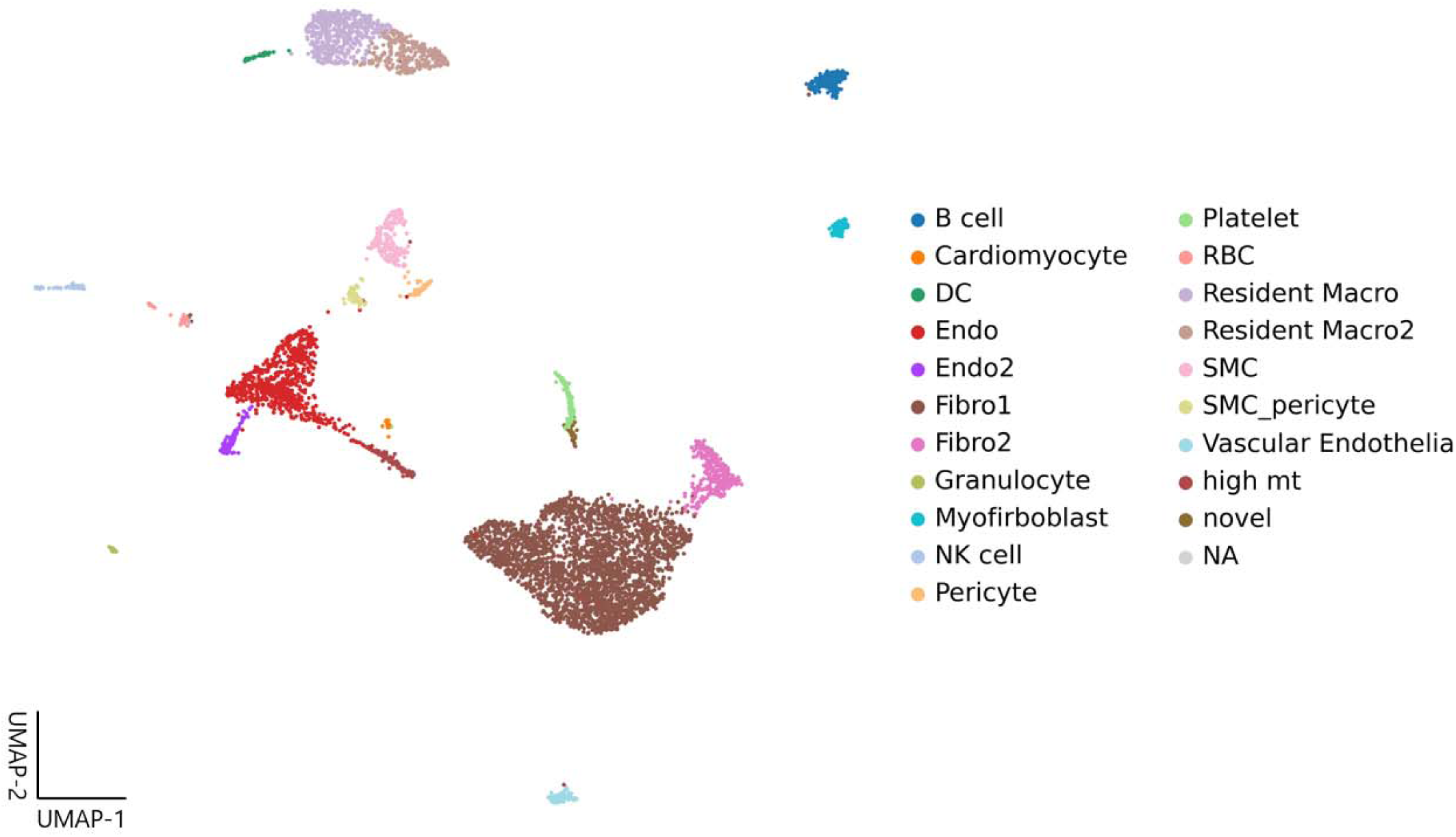
Targeted long-read sequencing of cells of interest with Ouro-Cell-Enrich. High-throughput scRNA-seq technologies, such as the 10x Genomics systems, generate a pooled single-cell cDNA library containing barcoded cDNAs from hundreds to thousands of single cells. However, due to the high cost of long-read sequencing, it is often beneficial to prioritize a subset of the cells captured in scRNA-seq experiments during the characterization of full-length transcriptomes of individual cells with long-read scRNA-seq. Therefore, we applied Ouro-Cell-Enrich to a pooled single-cell cDNA library containing barcoded cDNAs from 6,964 cells isolated from the ouseheart, enriching barcoded cDNAs from 99 cells of interest (myofibroblasts). A UMAP plot of all cells captured in the pooled single-cell cDNA library is shown, colored by annotated cell types.

**Extended Data Figure 5.**
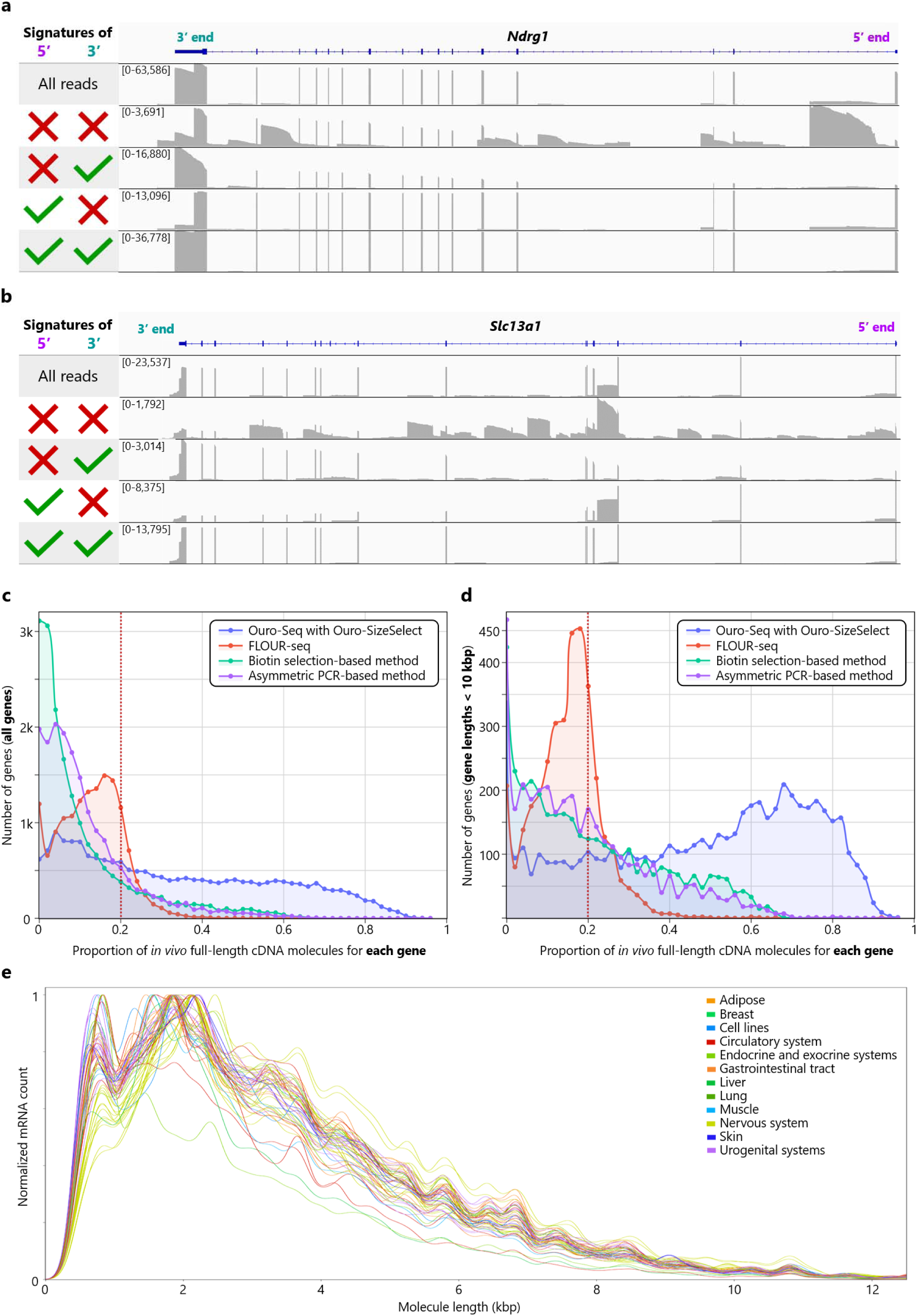
Removing technical variation specific to long-read scRNA-seq technologies with Ouro-Tools. **a**, **b**, After removing barcoding artifacts with Ouro-Seq, truncated cDNAs from mispriming events and degraded mRNAs are identified using the biological full-length identification module of Ouro-Tools. cDNAs containing an unencoded-G (5’ cap signature) and an enzymatically added poly(A) tail (3’ end signature) are classified as biological full-length cDNAs. As shown in the classification results of cDNAs aligned to *Ndrg1* (**a**), *Slc13a1* (**b**), and *Errfi1* (Figure 3d), the truncated cDNAs lacking 5’ and 3’ end signatures primarily represent degraded mRNAs and partial cDNAs generated through mispriming at A-rich tracts in introns and occasionally exons of pre-mRNAs and mRNAs. **c**, **d**, Histograms of proportions of *in vivo* full-length cDNA molecules for all protein-coding genes (**c**) and for protein-coding genes that are smaller than 10 kbp (**d**) across different barcoding artifact removal methods, indicating that the majority of protein-coding genes can be effectively analyzed by Ouro-Seq in full length (**Supplementary Table 4**); for the biotin selection-based and asymmetric PCR-based methods, the **Shiau2023** and **Hardwick2022** datasets were utilized, respectively. **e,** Utilizing ∼17,000 short-read RNA sequencing datasets covering 54 organs and tissues from the GTEx Consortium, we observed that the underlying mRNA size distributions of various organs and tissues are nearly identical after removing the effects of highly abundant, tissue-specific mRNAs on mRNA size distributions. Based on the assumption that individual single cells’ underlying mRNA size distributions are primarily identical, we developed the size distribution normalization module of Ouro-Tools.

**Extended Data Figure 6.**
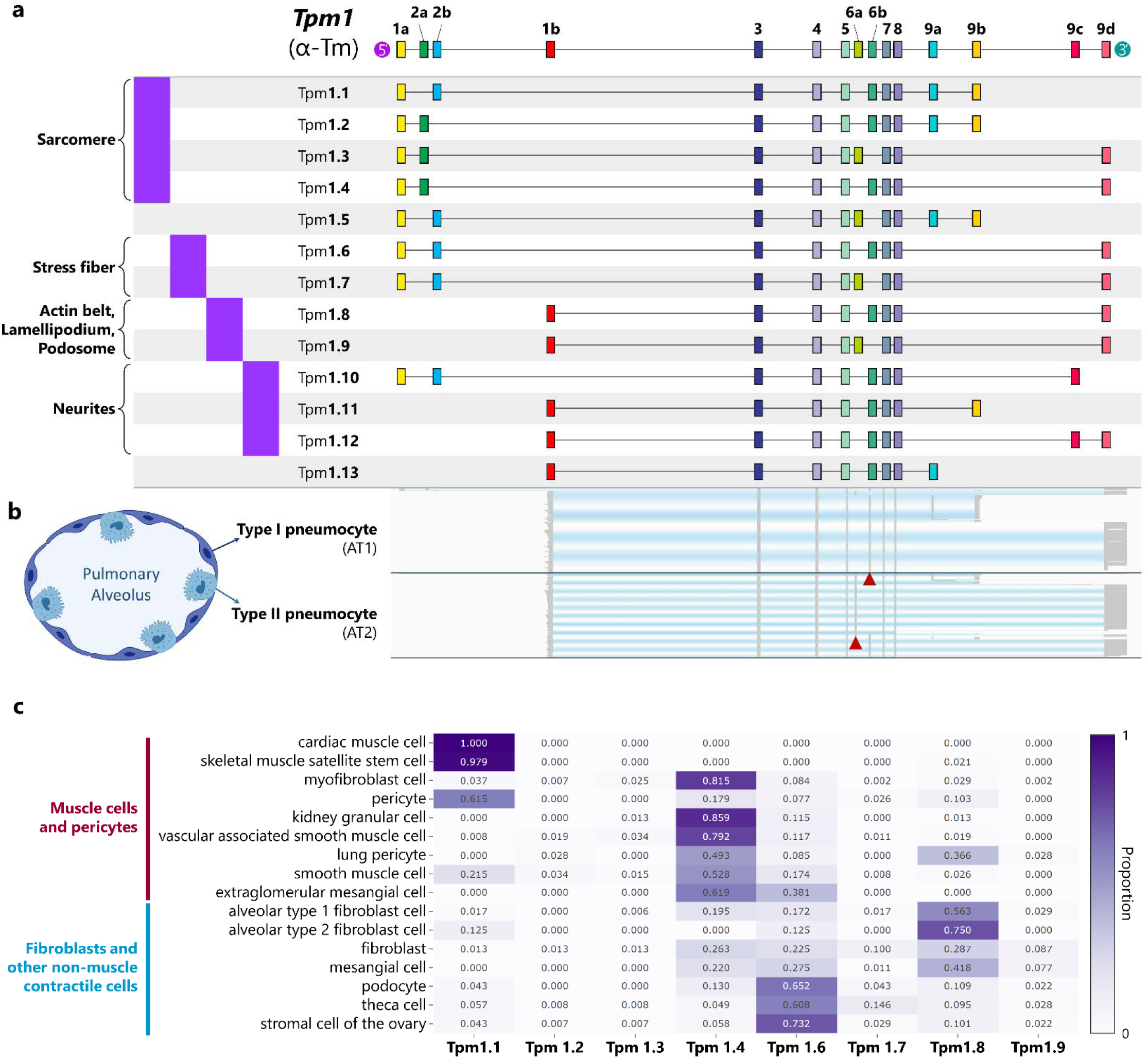
A compendium of *Tpm1* isoform expression patterns across cell types. **a**, Tropomyosins are a large family of highly conserved proteins constituting integral parts of most actin filaments in vertebrates, regulating interactions between actin-binding proteins and the actin filaments. Among the four tropomyosin genes encoding tropomyosins (*Tpm1*–*4*), the *Tpm1* gene displays the most complex variation in the spliced mRNA transcripts (**a**), generated by alternative promoter usage, alternative polyadenylation, and alternative splicing. Tpm1.1–1.4 are categorized as muscle tropomyosins, regulating muscle contractions in various muscle cells. Other tropomyosins encoded by *Tpm1* are categorized as non-muscle tropomyosins, regulating diverse cellular functions involving actin filaments in non-muscle cells, such as cell migration, neuron development, and maintenance of apical-basal polarity. **b**, Individual full-length *Tpm1* transcripts for type 1 and type II pneumocytes, respectively (an illustration of the alveolus was created with BioRender.com, accessed on 22 June 2024). Using antibodies, distinguishing between certain tropomyosin isoforms (e.g., Tpm1.8 and Tpm1.9) is often challenging due to extensive exon sharing between tropomyosin isoforms. We observed largely mutually exclusive expression patterns of Tpm1.8 and Tpm1.9 across diverse epithelial cell types; the simple cuboidal or columnar epithelial cells predominantly express Tpm1.9, while the simple squamous epithelial cells predominantly express Tpm1.8. **c**, Relative abundance of Tpm1.1–1.4 and Tpm1.6–1.9 across diverse contractile cell types, including muscle cells, pericytes, and fibroblasts.

**Extended Data Figure 7.**
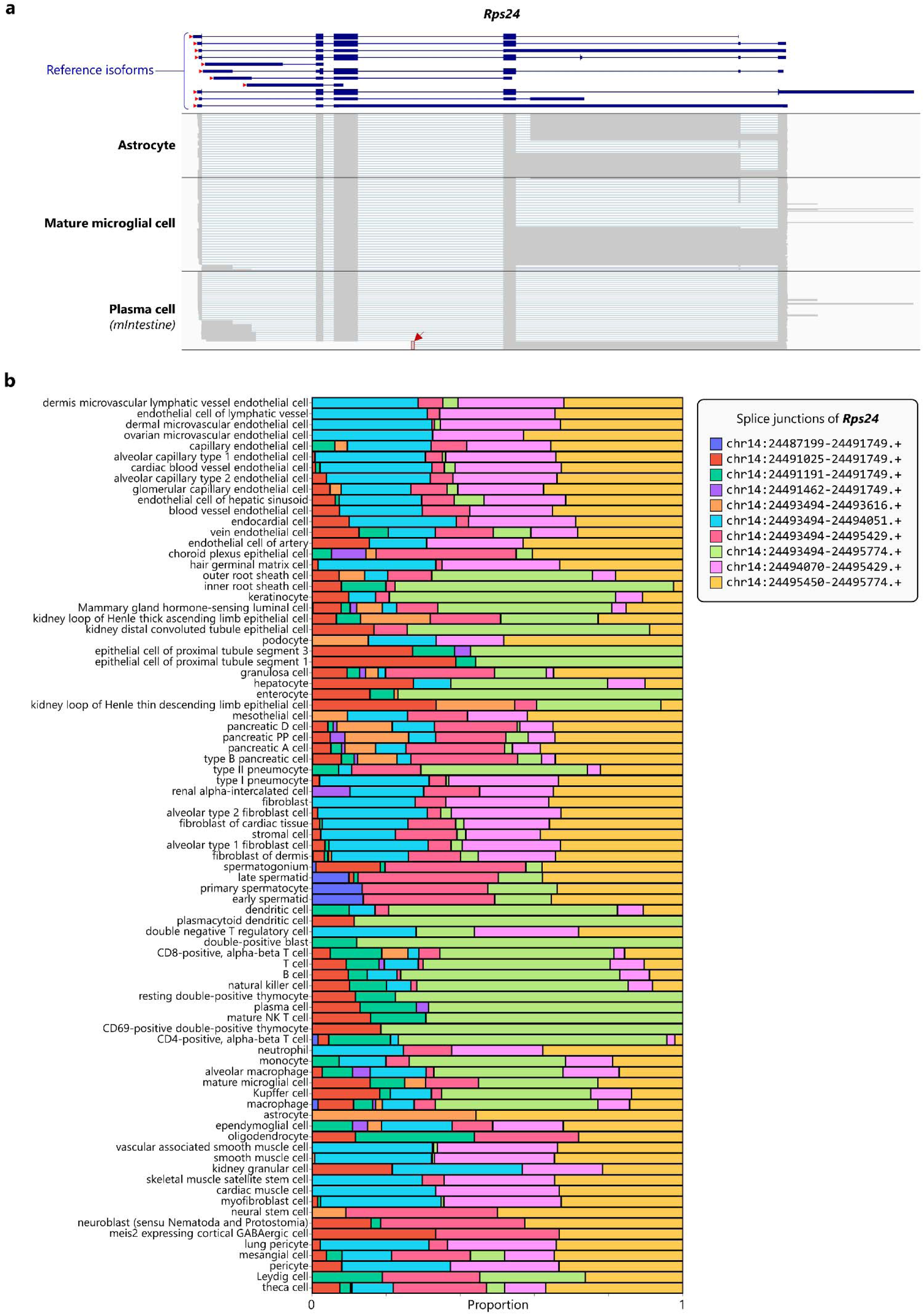
Comprehensive characterization of known and novel transcripts across a wide range of mRNA lengths. **a**, Expression patterns of a variety of known and novel *Rps24* mRNA isoforms, ranging widely in size from 468 bp to 4,059 bp, were characterized across cell types. Individual full-length *Rps24* transcripts for selected cell types are shown, including a 2.5 kbp-long *de novo Rps24* isoform specifically expressed by astrocytes. Additionally, a novel promoter of *Rps24* (indicated by a red arrowhead) was identified from novel *Rps24* transcripts expressed in plasma cells in the intestine; the reverse transcription signature of m7G-cap was detected from transcripts utilizing the novel promoter. **b**, Diverse *Rps24* transcript expression patterns were captured in our splicing atlas. Relative frequencies (size-distribution-normalized) of known *Rps24* splice junctions utilized by each cell type are shown.

**Extended Data Figure 8.**
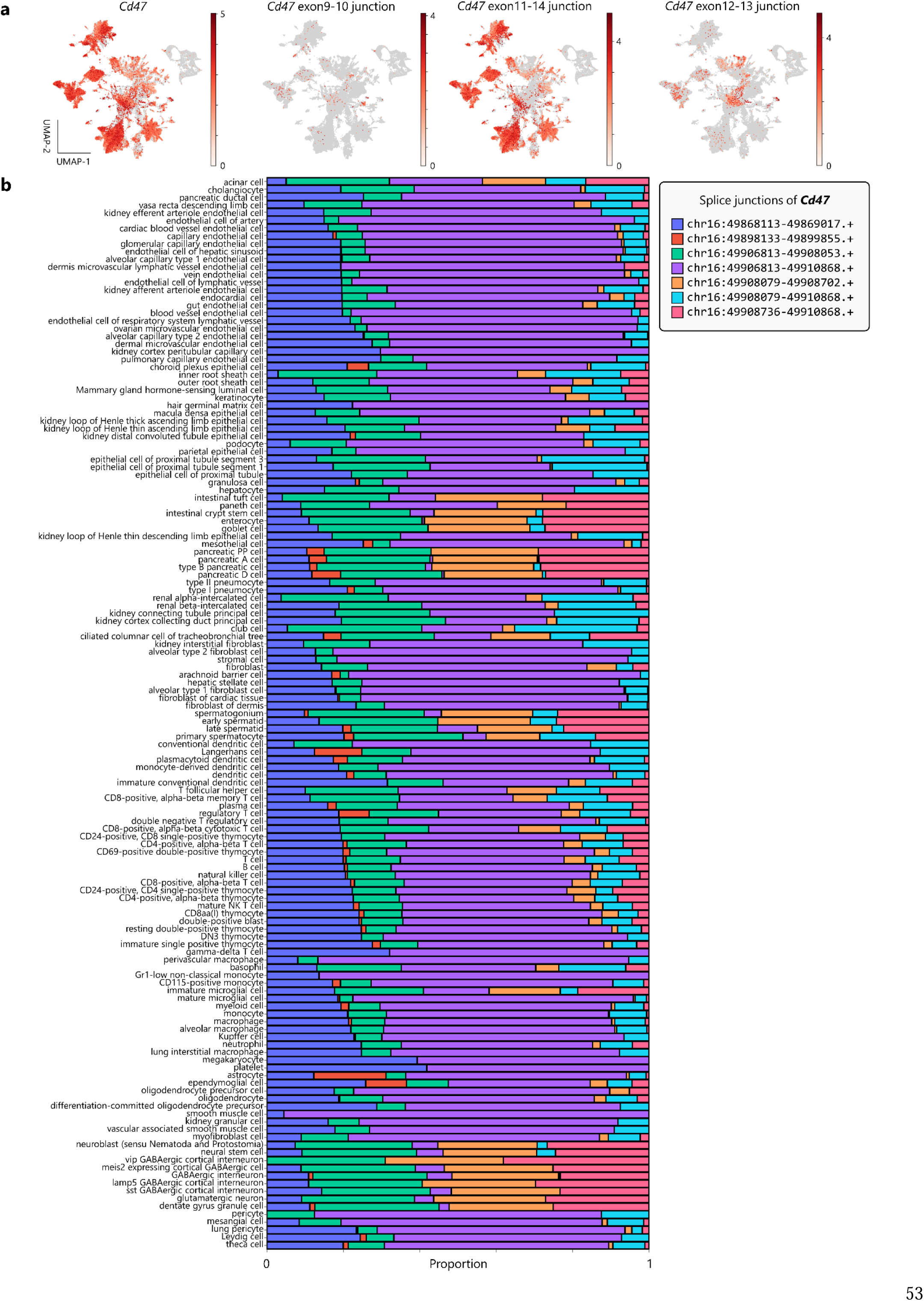
Alternative *Cd47* transcript usage across cell types. **a**, **b**, Diverse *Cd47* transcript expression patterns were captured in our splicing atlas (**a**, UMAP embeddings and **b**, stacked bar plots). Relative frequencies (size-distribution-normalized) of known *Cd47* splice junctions utilized by each cell type are shown.

**Extended Data Figure 9.**
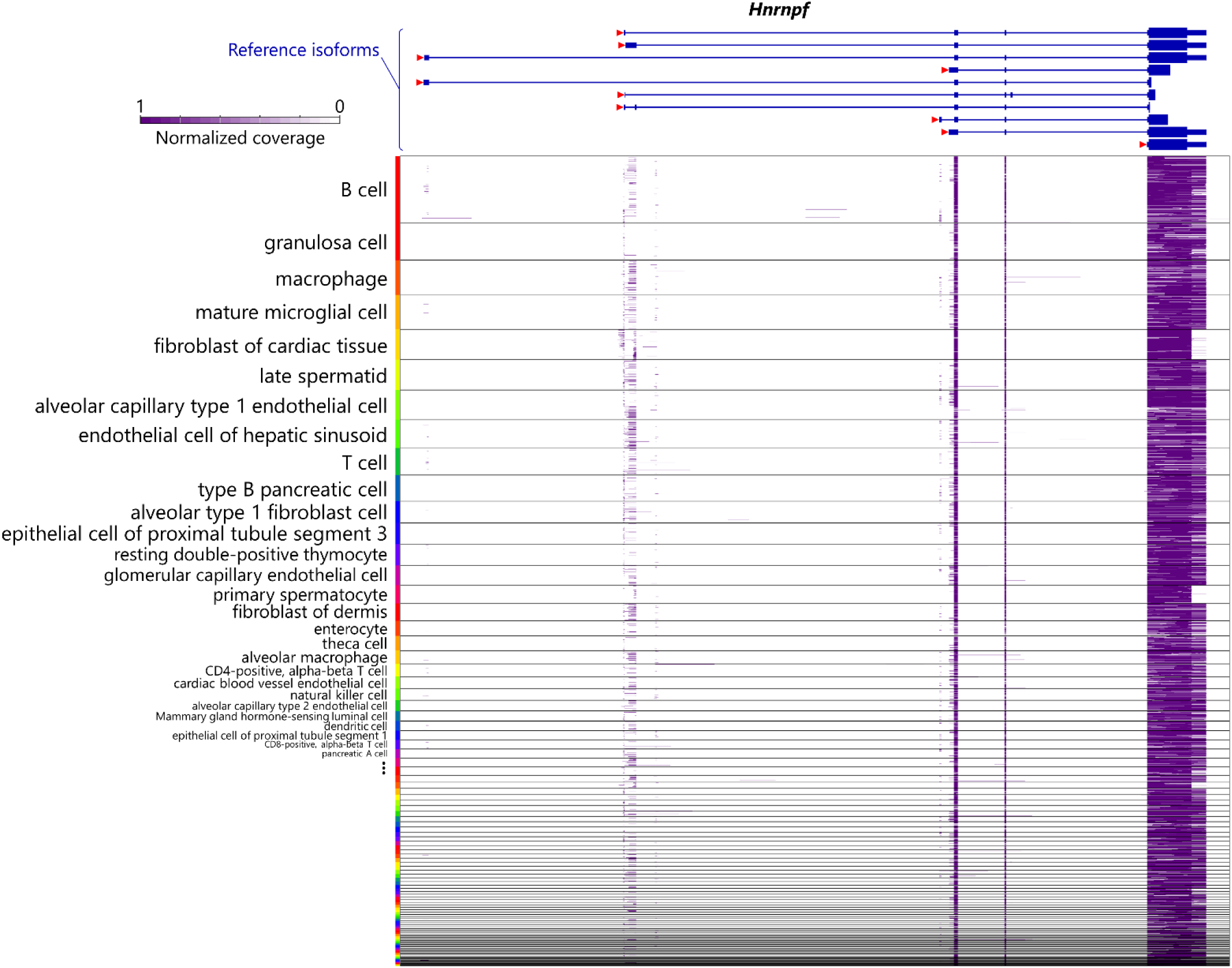
Characterization of alternative promoter usage and alternative polyadenylation usage across cell types. Coverage patterns of full-length cDNAs aligned to *Hnrnpf* at the resolution of individual cells. We observed that each single cell utilizes multiple promoters of *Hnrnpf* simultaneously to generate various *Hnrnpf* transcripts with different 5’ untranslated regions.

**Extended Data Figure 10.**
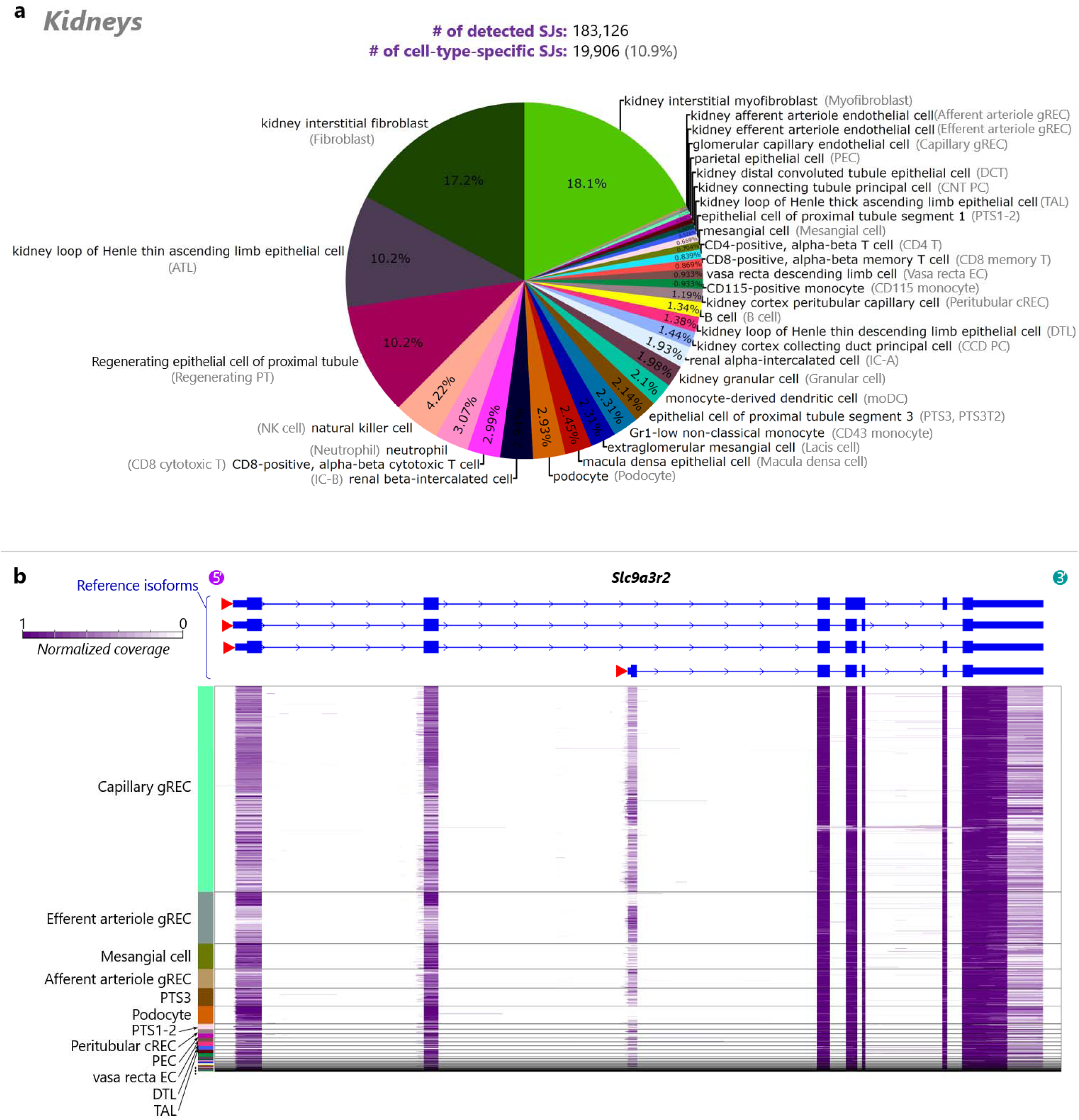
Cell-type-specific alternative splicing events detected in the mouse kidney. **a**, A pie chart of the numbers of unique marker splice junctions for each cell type in the mouse kidney. **b**, Single-cell-level coverage patterns of full-length cDNAs aligned to *Slc9a3r2*. Using the psuedobulk-based differential transcript usage detection module of Ouro-Tools, we identified various cell type-specific alternative splicing events of the genes related to the key functions of the organ, such as *Slc9a3r2*, which is known to regulate the function of sodium–hydrogen antiporter NHE3 in the proximal tubule. Detailed cell type names are shown in **a**.

**Extended Data Figure 11.**
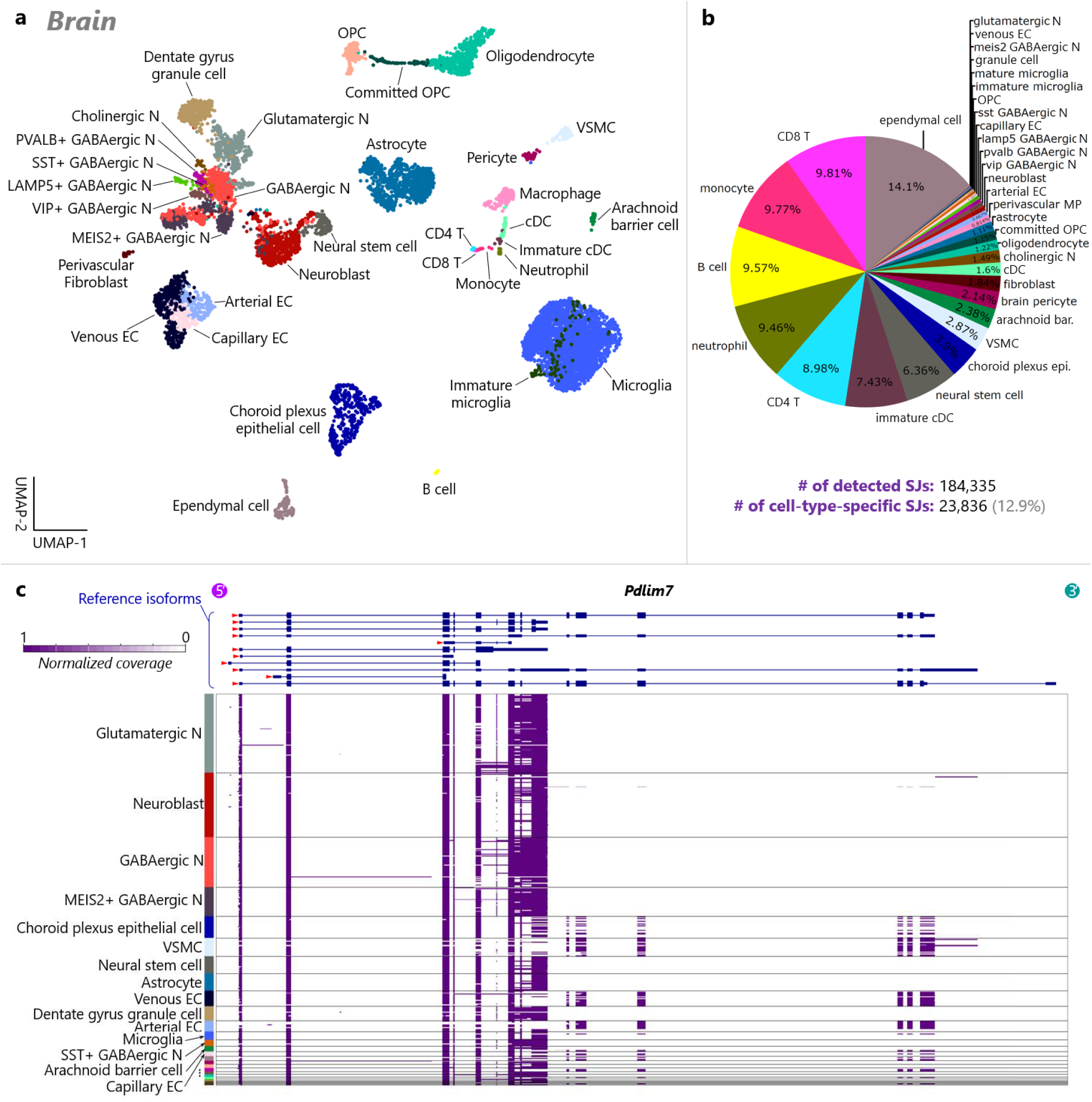
Cell-type-specific alternative splicing events detected in the mouse brain. **a**, A UMAP plot of cells and nuclei isolated from the mouse brain, colored by annotated cell types. **b**, A pie chart of the numbers of unique marker splice junctions for each cell type in the mouse brain. **c**, Single-cell-level coverage patterns of full-length cDNAs aligned to *Pdlim7*. N, neuron; OPC, oligodendrocyte precursor cell; cDC, conventional dendritic cell; EC, endothelial cell; VSMC, vascular smooth muscle cell.

**Extended Data Figure 12.**
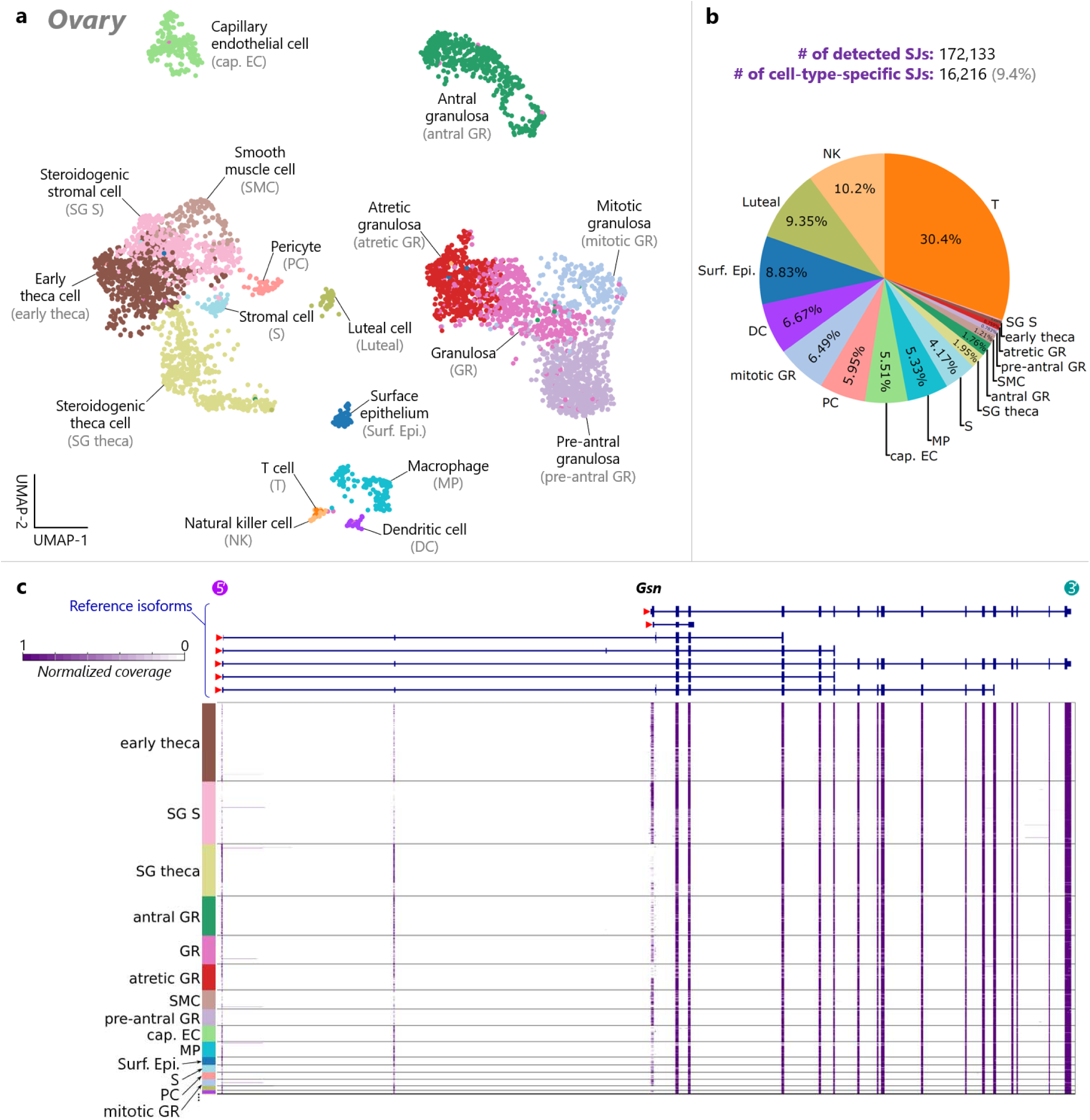
Cell-type-specific alternative splicing events detected in the mouse ovary. **a**, A UMAP plot of cells isolated from the mouse ovary, colored by annotated cell types. **b**, A pie chart of the numbers of unique marker splice junctions for each cell type in the mouse ovary. **c**, Single-cell-level coverage patterns of full-length cDNAs aligned to *Gsn*. Detailed cell type names are shown in **a**.

**Extended Data Figure 13.**
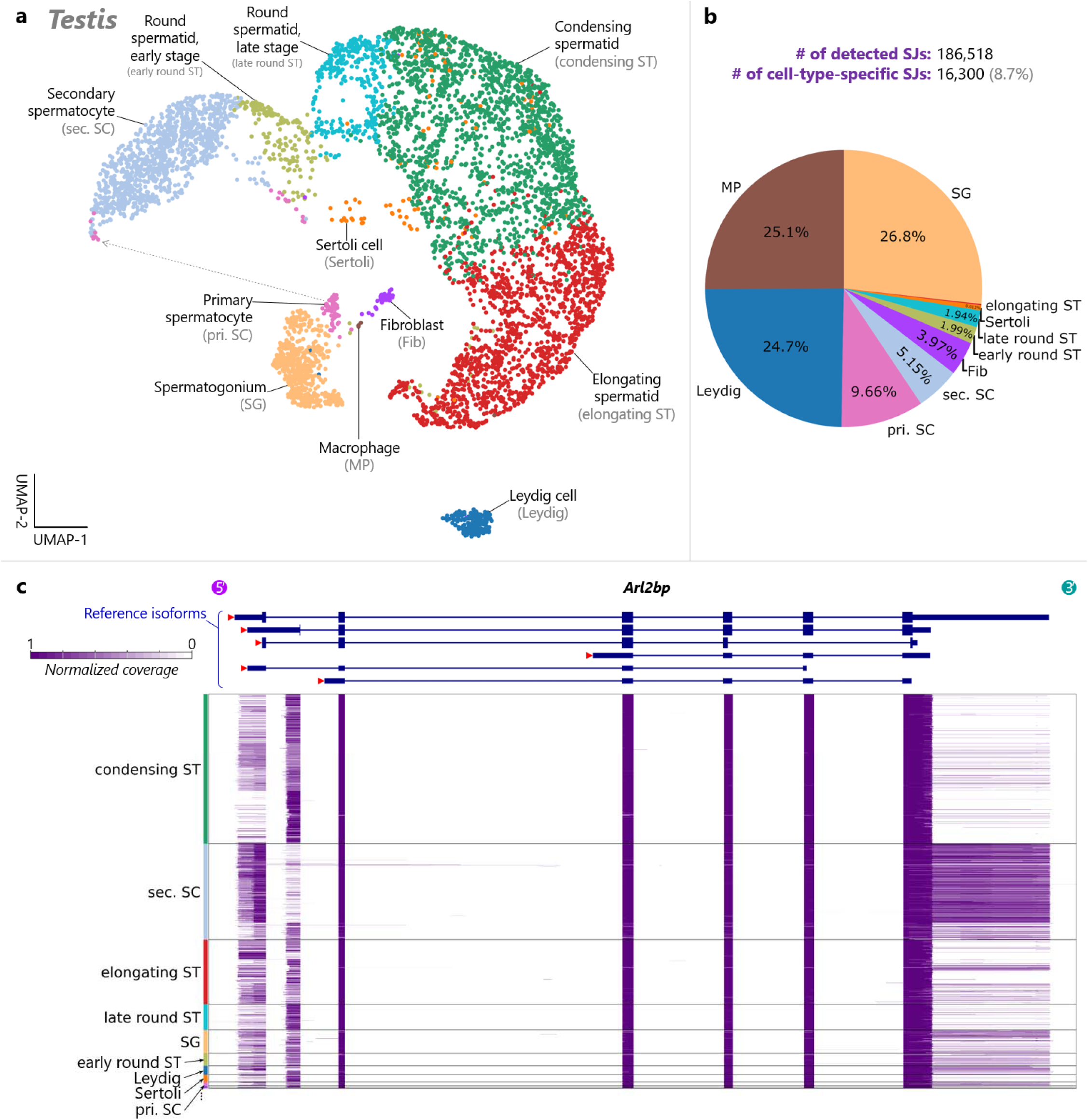
Cell-type-specific alternative splicing events detected in the mouse testis. **a**, A UMAP plot of cells isolated from the mouse testis, colored by annotated cell types. **b**, A pie chart of the numbers of unique marker splice junctions for each cell type in the mouse testis. **c**, Single-cell-level coverage patterns of full-length cDNAs aligned to *Arl2bp*. Detailed cell type names are shown in **a**.

**Extended Data Figure 14.**
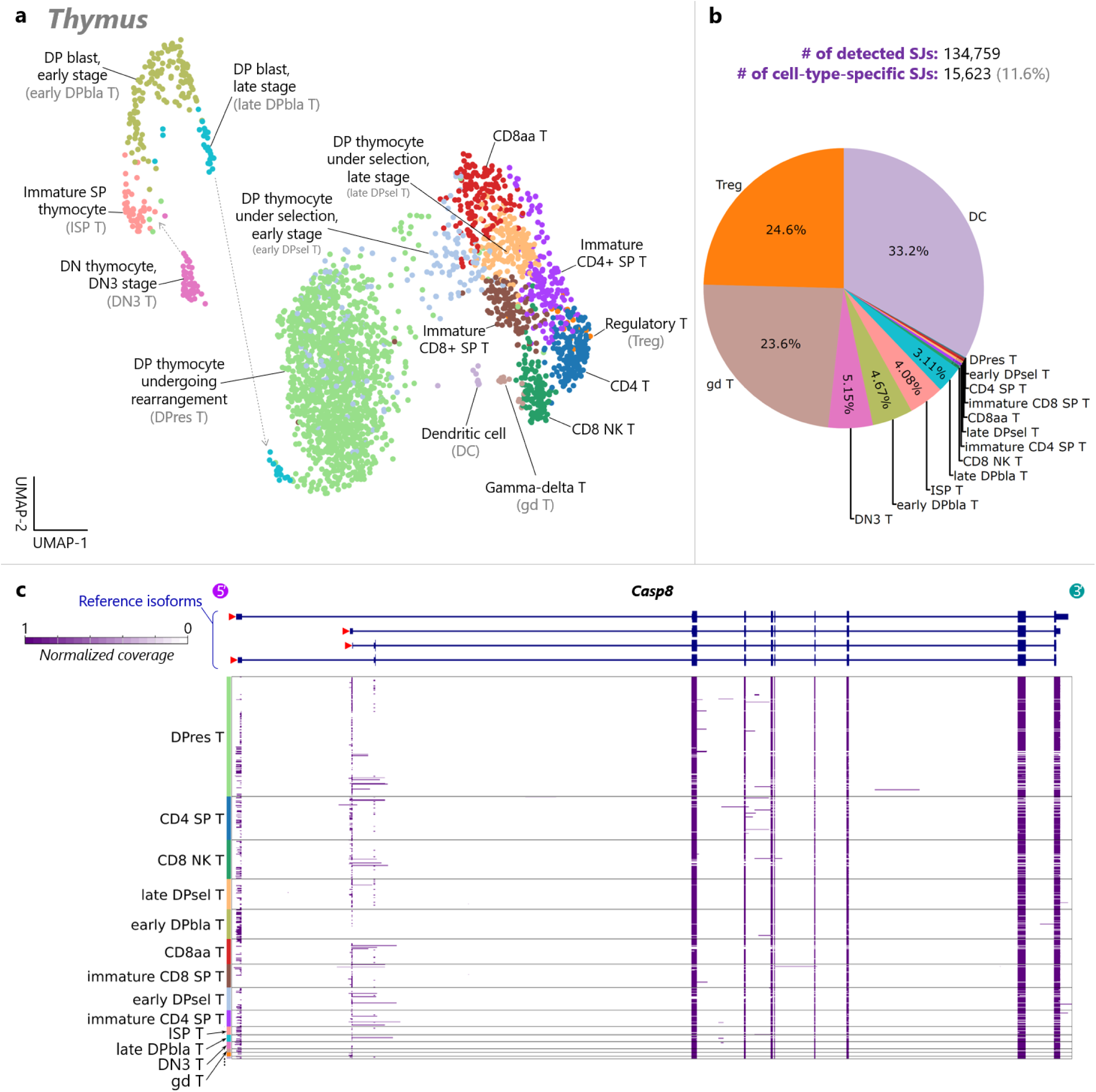
Cell-type-specific alternative splicing events detected in the mouse thymus. **a**, A UMAP plot of cells isolated from the mouse thymus, colored by annotated cell types. **b**, A pie chart of the numbers of unique marker splice junctions for each cell type in the mouse thymus. **c**, Single-cell-level coverage patterns of full-length cDNAs aligned to *Casp8*. Detailed cell type names are shown in **a**.

**Extended Data Figure 15.**
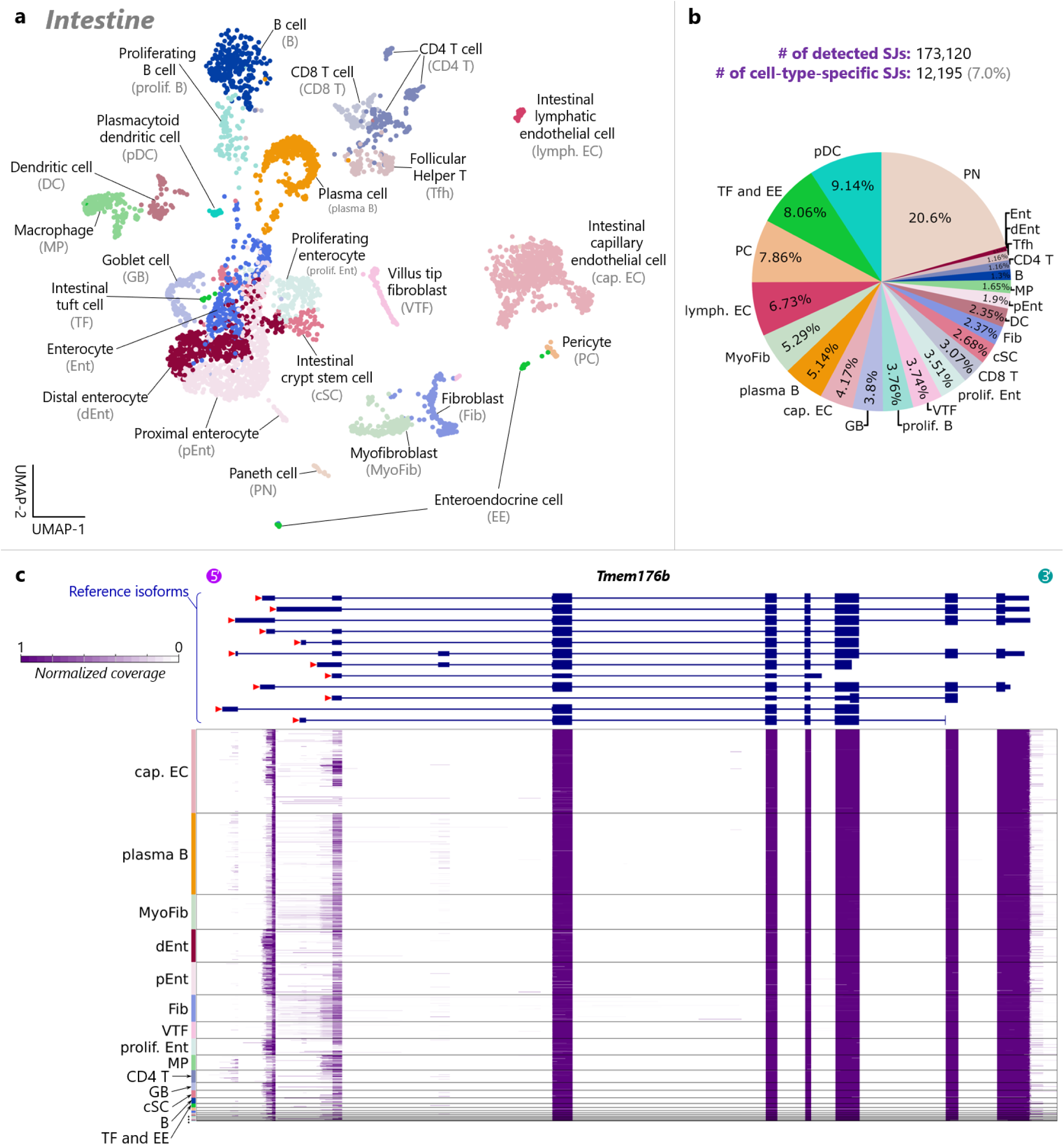
Cell-type-specific alternative splicing events detected in the mouse intestine. **a**, A UMAP plot of cells isolated from the mouse intestine, colored by annotated cell types. **b**, A pie chart of the numbers of unique marker splice junctions for each cell type in the mouse intestine. **c**, Single-cell-level coverage patterns of full-length cDNAs aligned to *Tmem176b*. Detailed cell type names are shown in **a**.

**Extended Data Figure 16.**
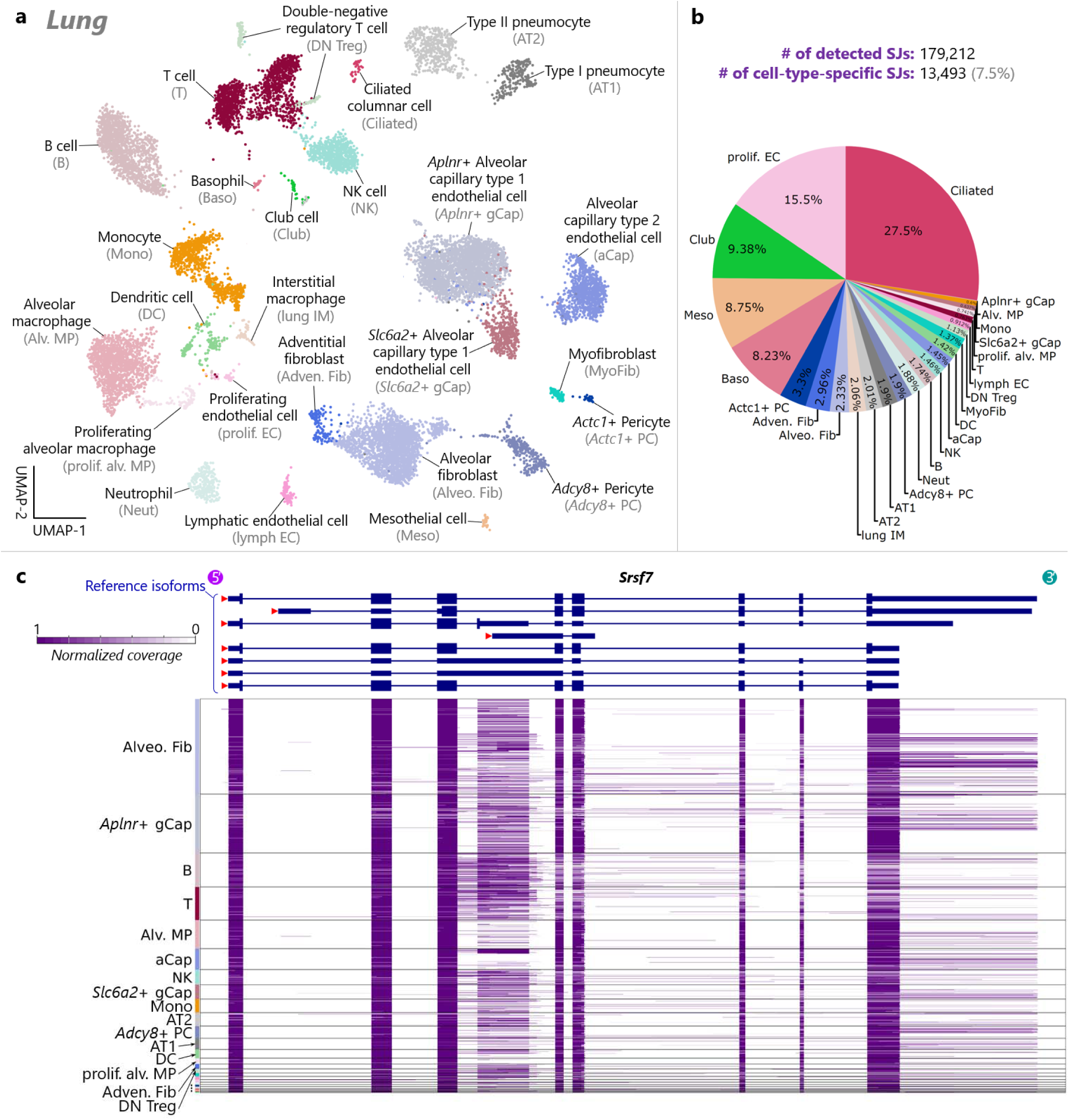
Cell-type-specific alternative splicing events detected in the mouse lung. **a**, A UMAP plot of cells isolated from the mouse lung, colored by annotated cell types. **b**, A pie chart of the numbers of unique marker splice junctions for each cell type in the mouse lung. **c**, Single-cell-level coverage patterns of full-length cDNAs aligned to *Srsf7*. Detailed cell type names are shown in **a**.

**Extended Data Figure 17.**
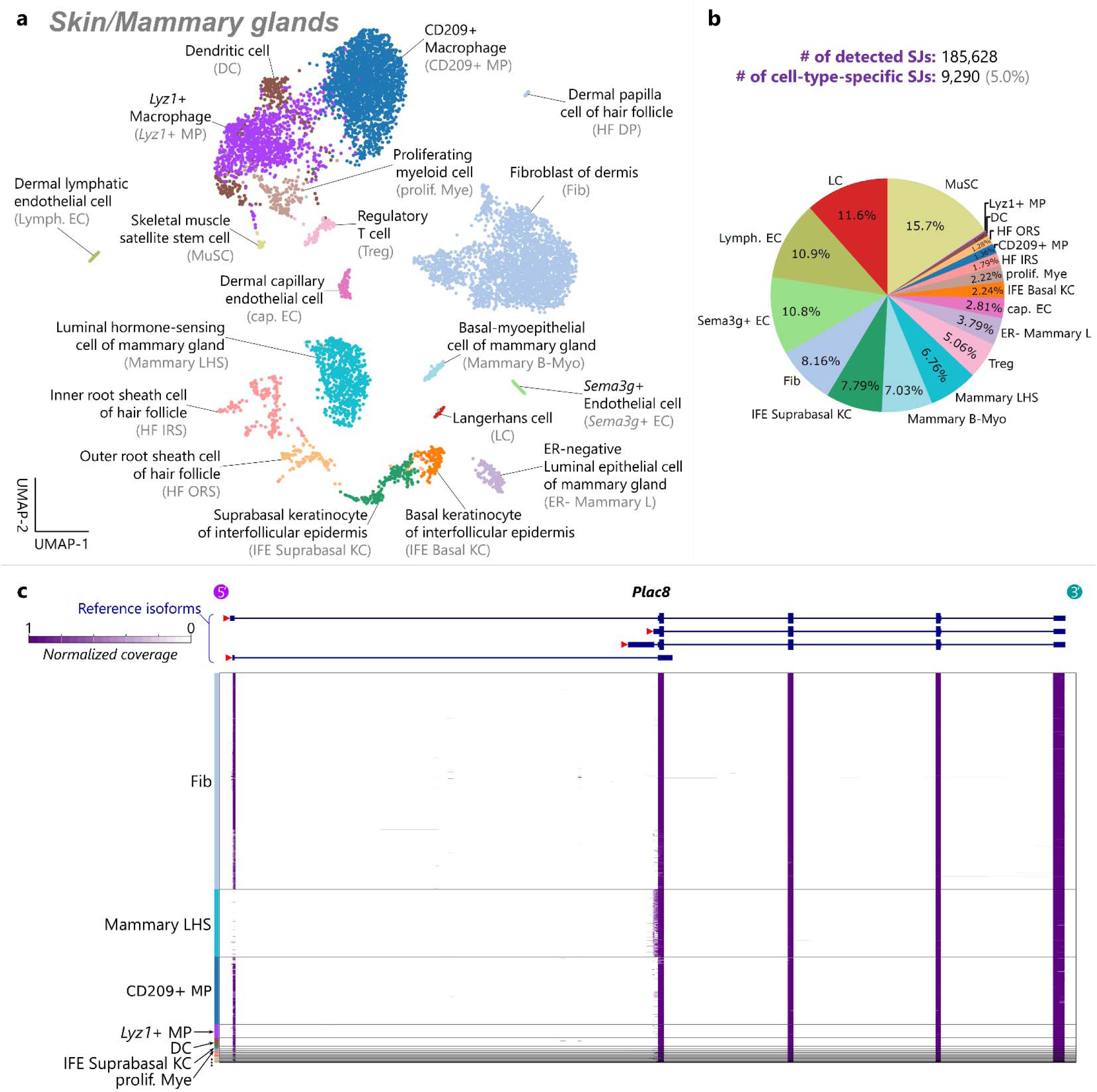
Cell-type-specific alternative splicing events detected in the mouse skin and mammary glands. **a**, A UMAP plot of cells isolated from the mouse skin and mammary glands, colored by annotated cell types. **b**, A pie chart of the numbers of unique marker splice junctions for each cell type in the mouse skin and mammary glands. **c**, Single-cell-level coverage patterns of full-length cDNAs aligned to *Plac8*. Detailed cell type names are shown in **a**.

**Extended Data Figure 18.**
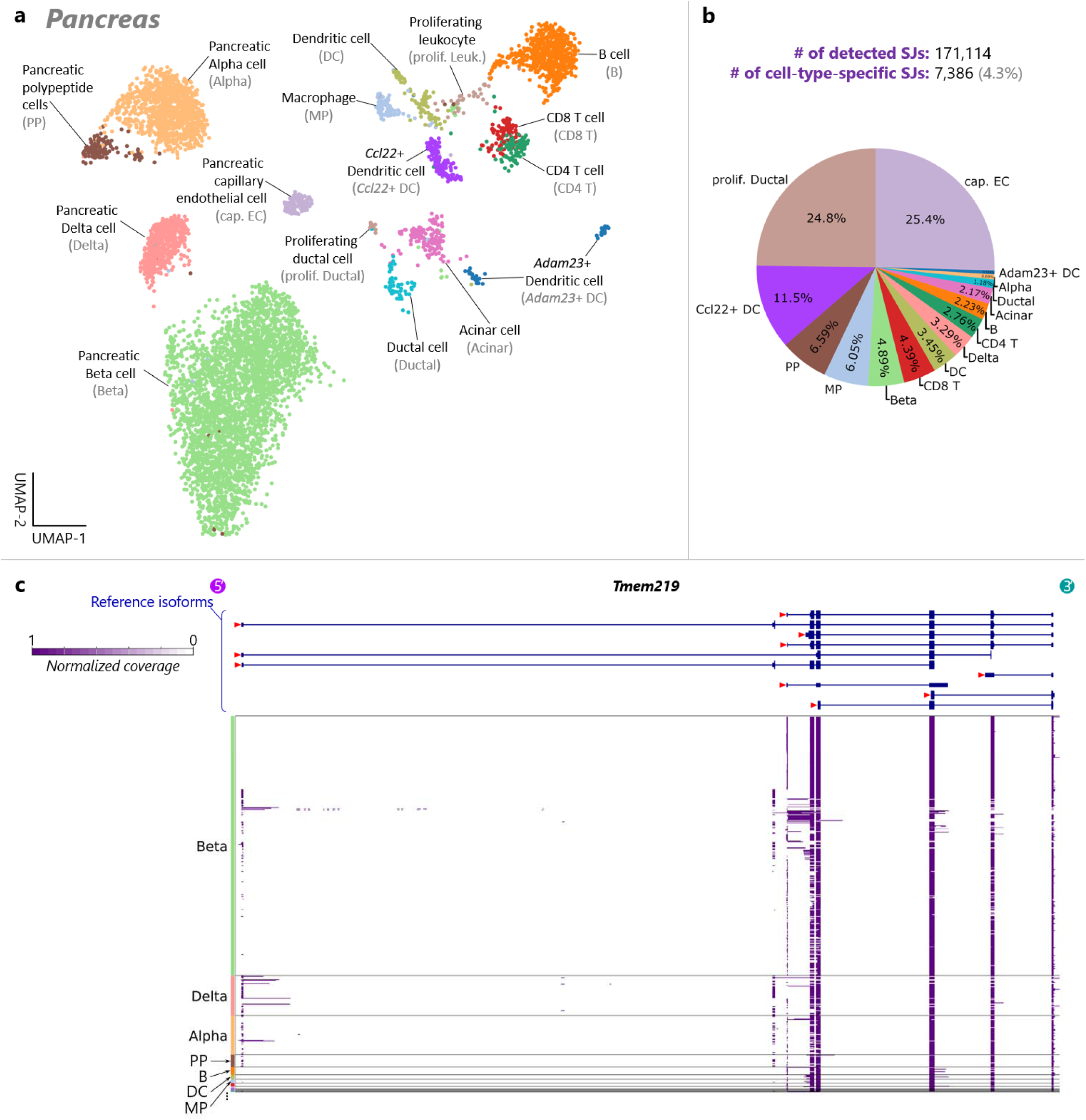
Cell-type-specific alternative splicing events detected in the mouse pancreas. **a**, A UMAP plot of cells and nuclei isolated from the mouse pancreas, colored by annotated cell types. **b**, A pie chart of the numbers of unique marker splice junctions for each cell type in the mouse pancreas. **c**, Single-cell-level coverage patterns of full-length cDNAs aligned to *Tmem219*. Detailed cell type names are shown in **a**.

**Extended Data Figure 19.**
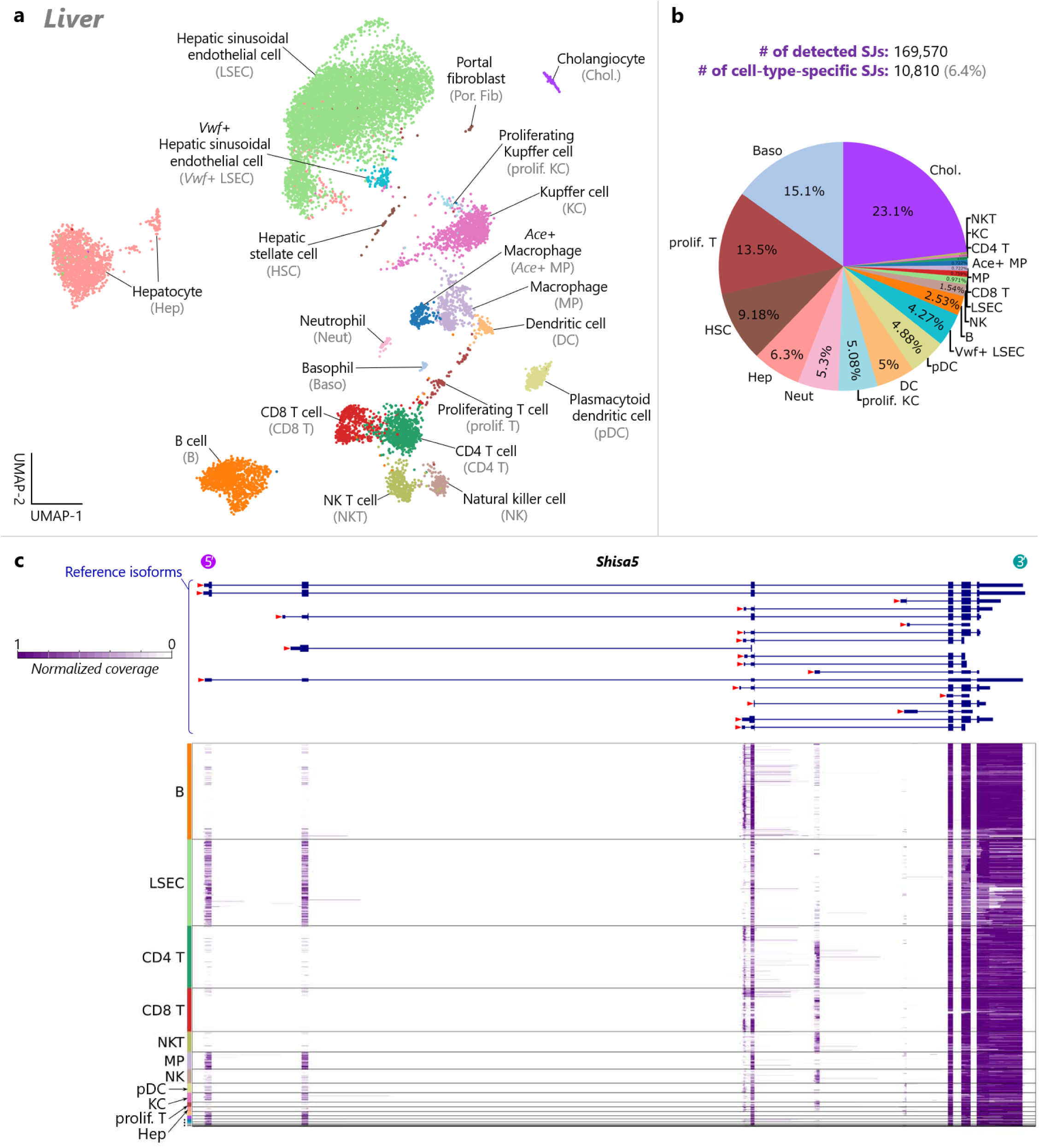
Cell-type-specific alternative splicing events detected in the mouse liver. **a**, A UMAP plot of cells and nuclei isolated from the mouse liver, colored by annotated cell types. **b**, A pie chart of the numbers of unique marker splice junctions for each cell type in the mouse liver. **c**, Single-cell-level coverage patterns of full-length cDNAs aligned to *Shisa5*. Detailed cell type names are shown in **a**.

**Extended Data Figure 20.**
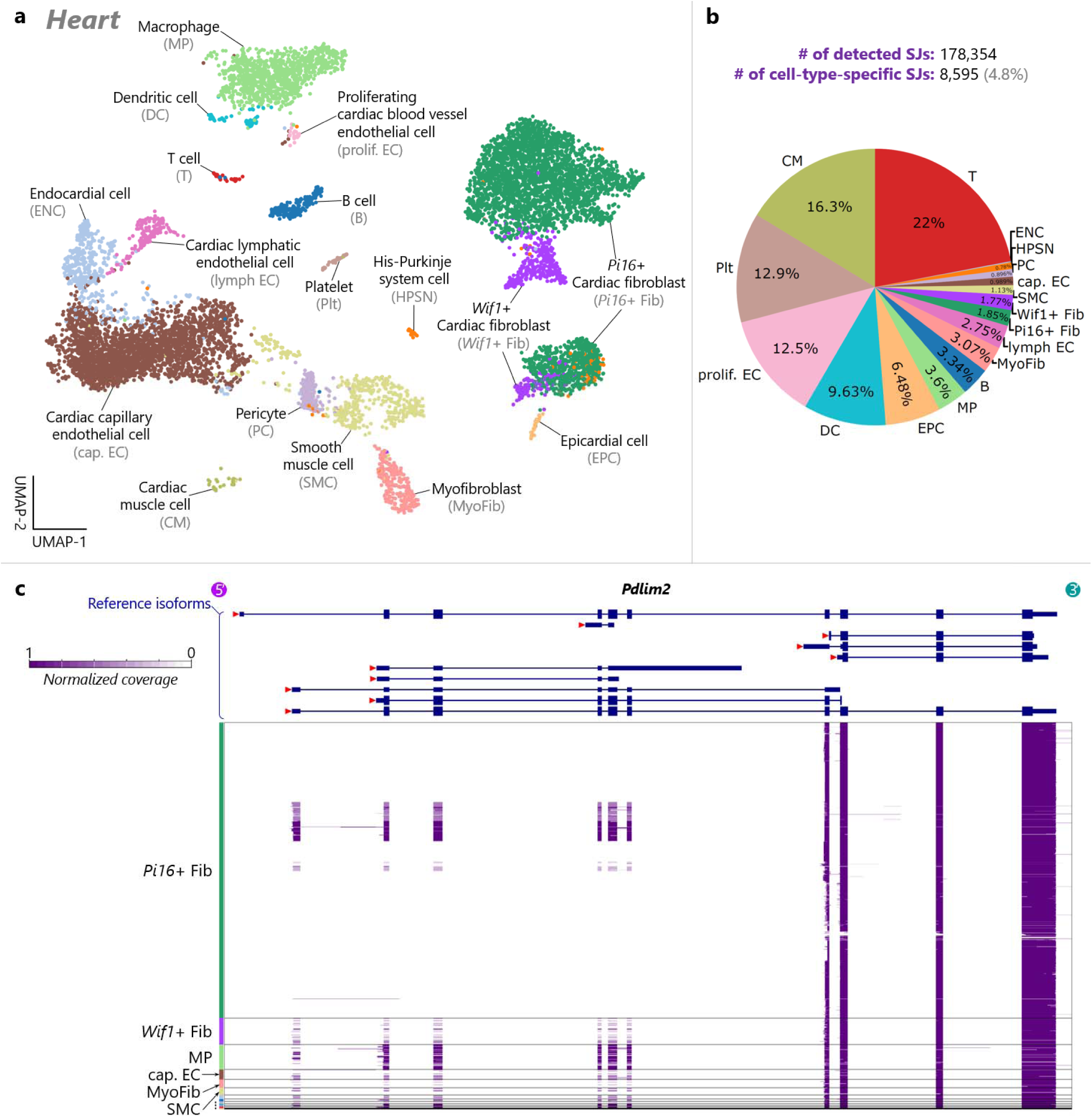
Cell-type-specific alternative splicing events detected in the mouse heart. **a**, A UMAP plot of cells and nuclei isolated from the mouse heart, colored by annotated cell types. **b**, A pie chart of the numbers of unique marker splice junctions for each cell type in the mouse heart. **c**, Single-cell-level coverage patterns of full-length cDNAs aligned to *Pdlim2*. Detailed cell type names are shown in **a**.

**Extended Data Figure 21.**
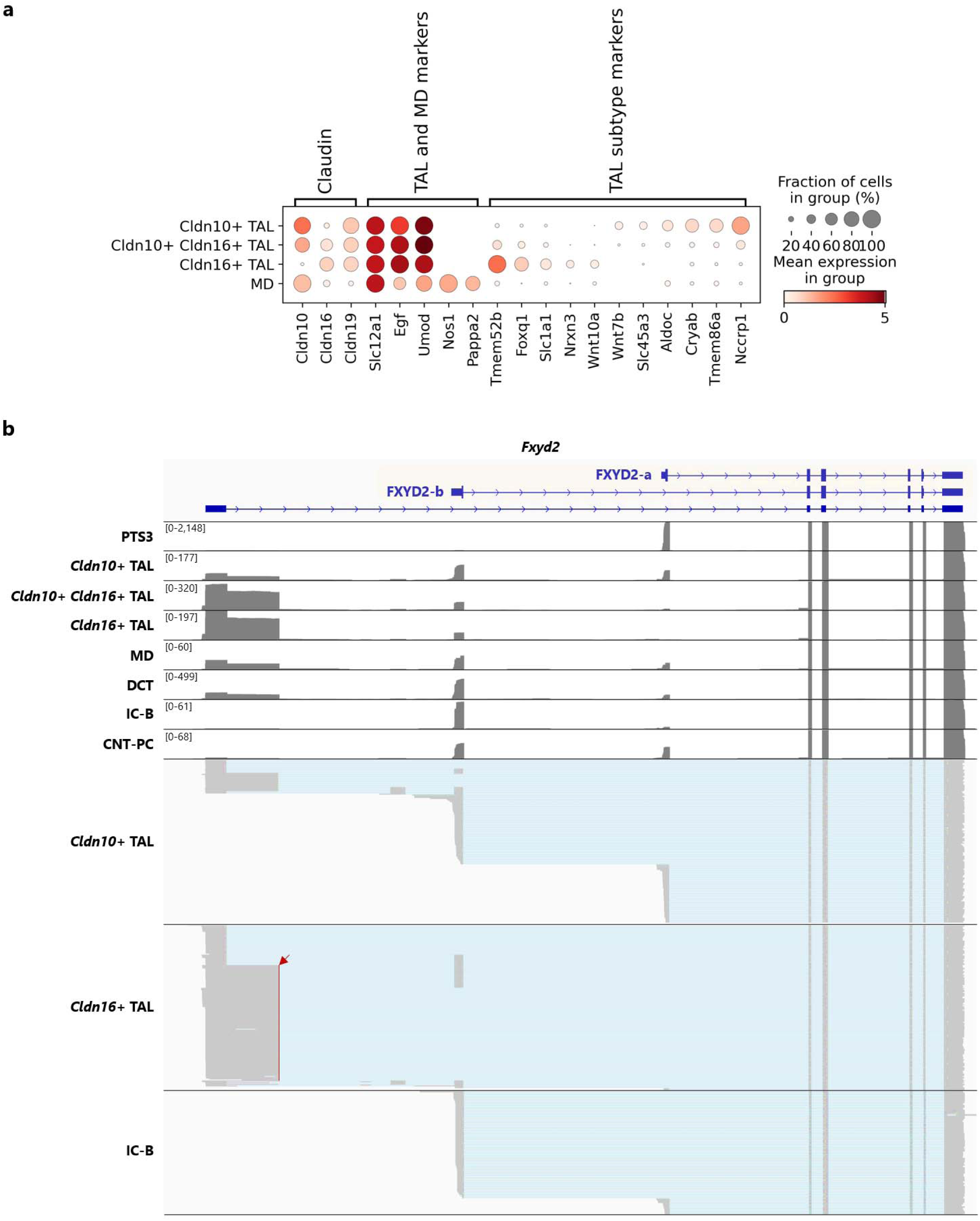
Unique isoform expression patterns of distinct cell populations in the thick ascending limb of the loop of Henle. **a**, Short-read-based gene-level analysis identified various unique marker genes for *Cldn10*+ TAL and *Cldn16*+ TAL populations, including previously identified marker genes for claudin-based subpopulations of TAL (**Supplementary Note 6**). However, no unique marker genes could be confidently identified for the *Cldn10*+ *Cldn16*+ TAL population. **b**, Individual full-length cDNAs aligned to *Fxyd2* for *Cldn10*+ TAL and *Cldn16*+ TAL subpopulations, including novel *Fxyd2* transcripts utilizing a novel 5’ exon with an alternative 5’ splice site (indicated by a red arrowhead).

**Extended Data Figure 22.**
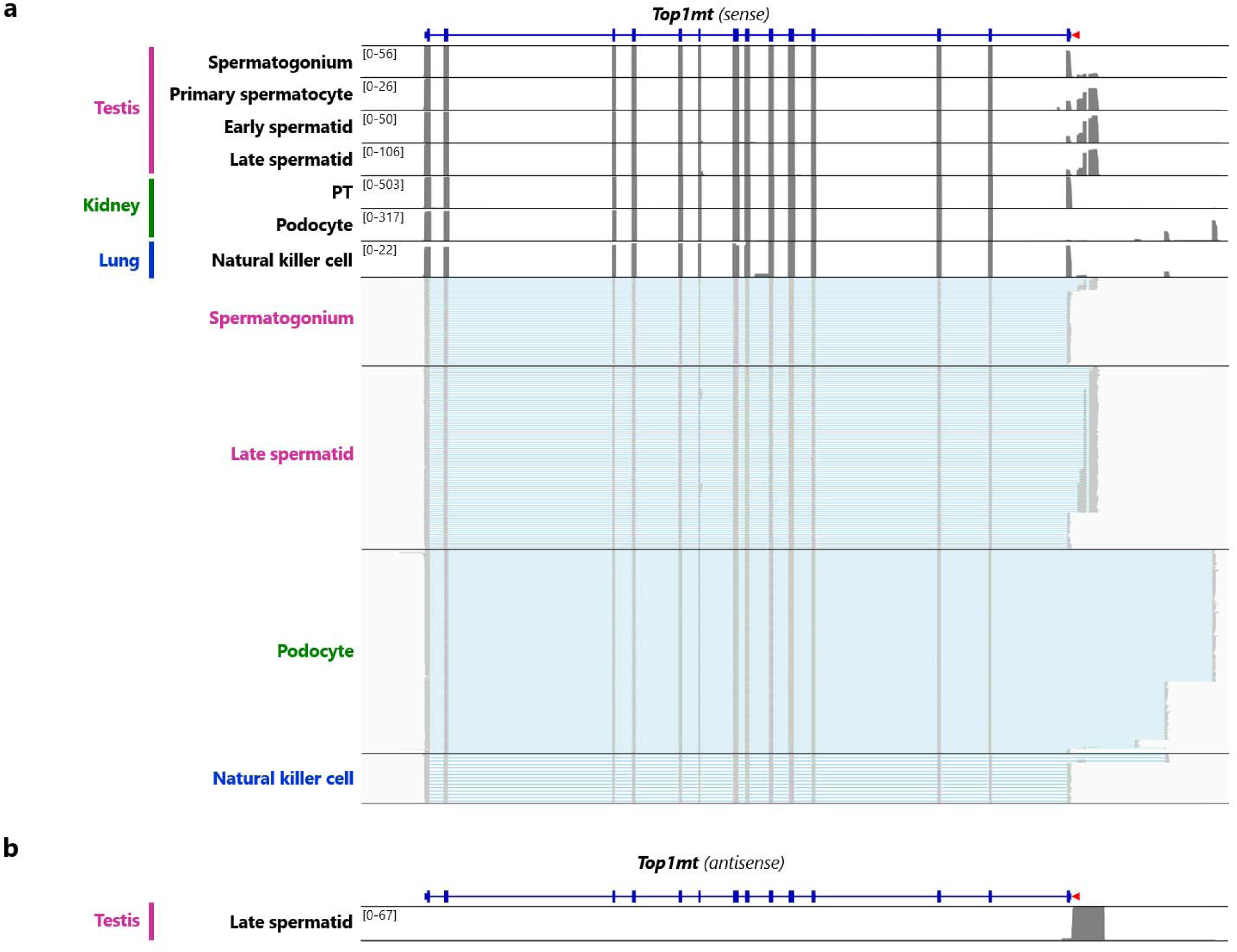
Discovery of novel promoters utilized by rare cell types. **a**, Individual full-length cDNAs aligned to *Top1mt* for several cell types utilizing various novel *Top1mt* promoters, replacing the original 5’ exon containing the start of the coding region with novel 5’ exons, likely altering the Top1mt protein structure and function. Interestingly, the podocytes exclusively utilized the novel *Top1mt* promoters, whereas the other cell types utilized the original promoter to a varying degree. Additionally, we identified a novel 9-th exon of Top1mt exclusively utilized by late spermatid cells. **b**, Late spermatids in the mouse testis expressed a transcribed regulatory element spanning the genomic region covering the novel *Top1mt* promoters specifically utilized by spermatids, with antisense orientation to the *Top1mt* transcripts.

**Extended Data Figure 23.**
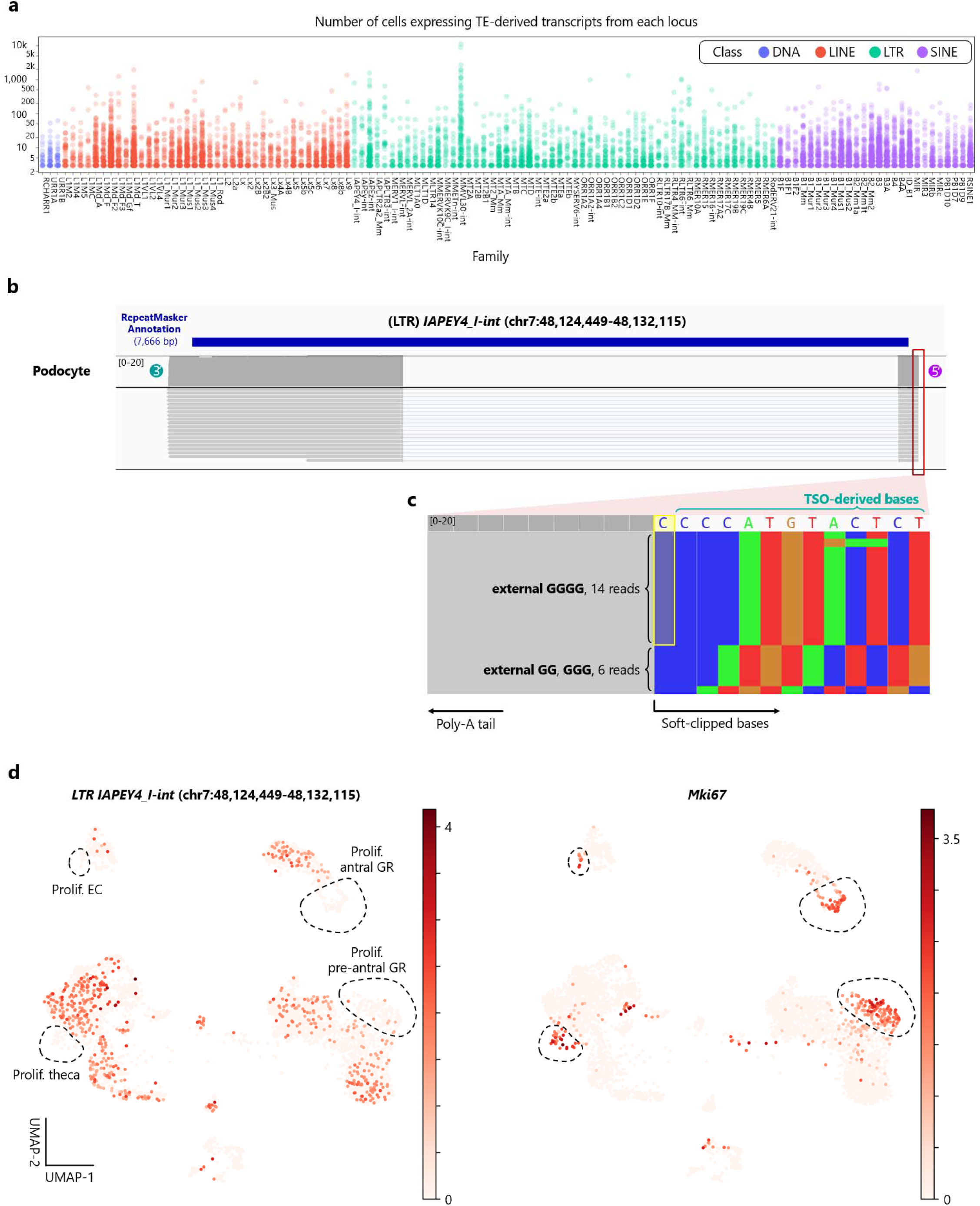
Locus-specific quantification of transposable element-derived transcripts across single cells. **a**, Using our atlas, we detected m7G-capped, polyadenylated RNA species from a variety of transposable elements across the genome. The number of cells expressing full-length RNA transcripts for each genomic locus is shown, colored by the class of the transposable elements. **b**, Individual full-length cDNAs aligned to an endogenous retroviral element, IAPEY4_I at chr7:48,124,449-48,132,115, expressed by podocytes. **c**, Presence of an unencoded G (annotated with the yellow box) in cDNAs aligned to the endogenous retroviral element. **d**, The endogenous retroviral element was highly expressed by several cell types in the ovary, including granulosa, theca, and luteal cells (the detailed cell type annotations can be found in **Extended Data Figure 12**). Interestingly, The endogenous retroviral element was exclusively expressed in cells in the G0 phase and silenced in proliferating cells, displaying a highly negative correlation with *Mki67* expression, a marker of proliferation. Prolif., proliferating; EC, endothelial cell; GR, granulosa cell.

## Notes

https://kbds.re.kr/KAP240772

### Competing Interest Statement

J.P., H.A., and S.C. are inventors on patent applications related to this work filed by Gwangju Institute of Science and Technology.

### Summary of Updates

Funding information was updated (J.P. acknowledges that this research was supported by the Samsung Science and Technology Foundation (SSTF-BA2001-11 to J.P.) and the National Research Foundation (NRF) of Korea funded by the Ministry of Science and ICT (MSIT) (RS-2024-00335026)).

