## Supplementary Information for Mouse Single-Cell Long-Read Splicing Atlas by Ouro-Seq for "Mouse Single-Cell Long-Read Splicing Atlas by Ouro-Seq"

### SUPPLEMENTARY NOTES

#### Supplementary Note 1: Limitations of single-cell full-length transcriptome analysis using short-read sequencing alone.

Massively parallel cell barcoding-based single-cell RNA sequencing (scRNA-seq) technologies, such as microdroplet-based (e.g., Drop-Seq^1^ and the 10x Genomics Chromium platform^2^), combinatorial indexing-based (e.g., SPLiT-seq^3^, sci-RNA-seq3^4^, and scifi-RNA-seq^5^), microwell-based (e.g., Microwell-seq 2.0^6,7^ and the BD Rhapsody platform^8^), and templated emulsification-based (e.g., PIP-seq^9^) technologies, can measure transcriptomes of thousands to even millions of individual cells. These scRNA-seq methods generate a pooled full-length cDNA library (i.e., a library containing single-cell-barcoded cDNAs from multiple cells) as an intermediate product after the initial reverse transcription (RT) reaction^10^. Each scRNA-seq method utilizes the 3’ or 5’ cell barcoding strategy^10^; cell barcode (CB) and unique molecular identifier (UMI) sequences are attached to either the 3’ or 5’ end of a full-length cDNA depending on the barcoding strategy. However, the barcoded full-length cDNAs are fragmented into short cDNA fragments in the size range between 200–600 bp to meet the requirements of a typical short-read sequencing experiment during the construction of a short-read scRNA-seq library. Consequently, the resulting short-read scRNA-seq library consists of the 3’ or 5’ end fragments of cDNAs (depending on where CB and UMI sequences were attached), generating sequencing data with extreme 3’ or 5’ end bias. As a result, a large portion of the rich information contained in a full-length cDNA is lost, including locations of the polyadenylation site and promoter utilized for transcription of an mRNA, a complete list of splice junctions utilized by the mRNA, all single nucleotide polymorphisms (SNPs) and single nucleotide variants (SNVs) covered by the mRNA, all RNA editing events occurred in the original mRNA, and genomic structural variants captured by the mRNA (i.e., the detection of a chimeric mRNA transcript).

Certain lower-throughput single-cell transcriptome profiling methods (e.g., Smart-seq2^11^, Smart-seq3^12^, and the Fluidigm C1 platform^13^) and massively parallel cell barcoding-based scRNA-seq methods utilizing random priming strategies (e.g., scFAST-Seq^14^ and SPLiT-Seq with random priming round1 barcoding step) are considered as short-read “full-length” scRNA-seq methods because transcriptome information of individual cells can be gathered throughout the gene body. However, since only the fragments of cDNAs (ranging in size from 200 to 600 bp) are analyzed by these methods, the full-length transcript structures have to be inferred^15^ from these short-reads with the help of transcript assembly tools, which has been known to assemble only 20%–40% of the human transcriptome^16,17^. Moreover, a recent comprehensive analysis^18^ of the Smart-seq2 datasets of *Tabula Muris Senis*^19^ and *BICCN Cortex^20^* revealed pervasive coverage artifacts in datasets generated by these short-read “full-length” scRNA-seq methods, which greatly hinder accurate transcript quantification in individual cells and introduce significant batch effects across datasets; consequently, the researchers focused on quantification of splice junctions sharing the same 3’ (or 5’) splice sites across cells, which is less affected by the presence of coverage artifacts.

Previous long-read scRNA-seq studies revealed that reverse transcription and PCR artifacts account for a significant portion of the full-length cDNA library^21,22^, significantly reducing the efficiency of long-read scRNA-seq. With long-read sequencing, these artifact molecules can be easily distinguished from genuine full-length cDNA molecules. In contrast, short-read “full-length” scRNA-seq methods have no^11^ or limited^12^ capabilities to identify and exclude cDNA fragments originating from these artifact molecules, manifesting as pervasive coverage artifacts in short-read “full-length” scRNA-seq datasets.

#### Supplementary Note 2: Unintentional amplification of cell barcode-doublet artifacts by the asymmetric PCR-based artifact depletion methods

Asymmetric PCR-based cell barcode-free artifact depletion methods increase the number of cell-barcode-containing molecules in the library through selective linear amplification of the molecules. However, asymmetric PCR inadvertently amplifies cell barcode-doublet artifacts exponentially, containing the same type of PCR handle at both ends; a detailed description of cell barcode-doublet artifacts, including the possible mechanisms by which these artifacts can be generated during scRNA-seq experiment, can be found in the official 10x Genomics Document #CG000376.

The asymmetric PCR-based method was first utilized in SnISOr-Seq^23^ (LAP-CAP), the latest single-cell long-read RNA-sequencing method that enabled an efficient characterization of cell type-specific alternative splicing patterns using single-nuclei cDNA libraries, which typically contains a higher proportion of cell barcode-free artifacts than single-cell cDNA libraries. We observed that after applying the asymmetric PCR-based artifact depletion method, the proportion of cell barcode-doublet artifacts to all the cell-barcode-containing molecules in the library significantly increased from ~1% (the baseline level) to ~50% (**Figure 1f** and **Extended Data Figure 2a**). Since the PCR amplification of the cell barcode-doublet artifacts smaller than ~2 kbp is suppressed due to the intramolecular complementary base pairing of its two ends (i.e., PCR-suppression^24^), the fraction of cell barcode-doublet artifacts was somewhat smaller for the cDNAs smaller than 2 kbp (20%–45%) but still significantly larger than that of the raw cDNA library (~1%) before applying the method (**Extended Data Figure 2b**).

#### Supplementary Note 3: Diverse transcript usage patterns of *Tpm1* across cell types

The *Tpm1* gene is one of the four tropomyosin genes that are highly conserved across vertebrate species^25^. Through combination of alternative promoter usage, alternative polyadenylation, and alternative splicing, the four tropomyosin genes of vertebrates collectively generate more than 40 known tropomyosins (Tpm), a diverse family of proteins that co-polymerize with filamentous actin (F-actin) and regulate various properties and functions of actin filaments *in vitro*^26^, which appear to be responsible for the structural and functional diversification of the actin filament systems in fungi and metazoa^27^. The comprehensive variety of tropomyosin isoforms of vertebrates supports various cellular functions involving actin filaments, such as epithelial–mesenchymal transition (EMT), cell migration, cell division, mechano-transduction, and barrier functions^28^. The two highly similar tropomyosin isoforms encoded by *Tpm1*, Tpm1.8 and Tpm1.9 are essential components of the actin belt near the apical surface of epithelial cells^29^, the actin structures of lamellipodia^30,31^, and the actin structures involved in focal adhesion assembly and mechanosensing^32^ (**Extended Data Figure 6a**). Interestingly, we observed largely mutually exclusive expression of either Tpm1.8 or Tpm1.9 in most epithelial cell types. For example, type I pneumocytes, which are simple squamous epithelial cells, predominantly expressed Tpm1.8, while type II pneumocytes, which are simple cuboidal epithelial cells, predominantly expressed Tpm1.9 (**Extended Data Figure 6b**). Notably, pancreatic islet beta and alpha cells primarily expressed Tpm1.9 like many simple cuboidal or columnar epithelial cells (**Extended Data Figure 6c**). Even though the pancreatic islet cells do not display typical tubular epithelium structures, the cells are clustered in rosette-like structures centering veins with artery-to-vein polarity^33,34^, which might explain this unexpected observation.

Consistent with this observation, since proximal tubule cells are simple cuboidal epithelial cells, Tpm1.9 was the most and second most highly expressed *Tpm1* isoform in the PTS1/2 and PTS3 epithelial cells, respectively. Surprisingly, we observed that the muscle tropomyosin Tpm1.1 is abundantly expressed in proximal tubule cell (**Figure 4d**), especially in epithelial cells of proximal tubule segment 1 and 2 (PTS1/2 or proximal convoluted tubule), where Tpm1.1 is the most highly expressed *Tpm1* isoform. Considering proximal tubular endocytosis is more prominent in PTS1/2, the higher expression of Tpm1.1 may be related to the highly developed endolysosomal system of PTS1/2.

Moreover, we discovered a variety of *de novo Tpm1* transcripts with novel splice junctions. For instance, previously uncharacterized *Tpm1* transcripts^35^ were expressed by oligodendrocyte precursor cells (OPCs), differentiation-committed OPCs, and mature oligodendrocytes (**Figure 4d**). Interestingly, once committed to differentiation, the committed OPCs lost the expression of the novel *Tpm1* mRNA isoforms containing the “1a” exon of *Tpm1*, assuming the transcript usage pattern of mature oligodendrocytes. Also, GABAergic neurons and *Meis2*+ GABAergic neurons expressed previously uncharacterized tropomyosin transcripts^35^ and presynaptic compartment-specific tropomyosin isoforms Tpm1.10 and Tpm1.12^36^ (**Figure 4d**). Notably, Tpm1.12 was expressed in GABAergic neurons but not in *Meis2*+ GABAergic neurons. In summary, our atlas accurately captured a compendium of expression patterns of both known and previously unknown mRNA transcripts across diverse cell types, as exemplified by the transcripts encoded in *Tpm1*.

#### Supplementary Note 4: Previous observations indicating cellular heterogeneity in the thick ascending limb of the loop of Henle

As an organ responsible for maintaining homeostasis by regulating extracellular fluid volume and composition, the kidney requires a high level of coordination between various cell types in the nephron. Tubuloglomerular feedback (TGF) system is one such example, where sodium and chloride ion concentrations in the tubular fluid are actively monitored by macula densa (MD) cells at the end of the thick ascending limb of the loop of Henle (TAL)^37^, which is communicated to the myocytes of afferent arteriole of the glomerulus, which adjust the glomerular filtration rate (GFR) accordingly. Therefore, we analyzed the full-length transcriptomes of cells isolated from the mouse kidney for a more comprehensive characterization of the cell types in the nephrons.

Analogous to transcellular channels, tight junctions (TJs) constructed by distinct sets of claudin proteins function as highly selective paracellular ion channels throughout the renal tubule. For example, Cldn10a is involved in paracellular Cl^–^ transport in the proximal tubule^38^, while Cldn10b is involved in paracellular Na^+^ transport in the thick ascending limb^39^ (**Figure 5b**). The epithelial cells of the thick ascending limb also express other members of claudin genes, *Cldn3*, *Cldn16*, and *Cldn19*^39^. Interestingly, there have been independent observations that the tight junctions (TJs) of the thick ascending limb in the cortex and the outer stripe of the outer medulla (OSOM) display mosaic patterns, where there are stretches of tight junctions only containing Cldn10b while the remaining tight junctions are entirely devoid of Cldn10b^39,40^; these Cldn10b-negative tight junctions are composed of Cldn3, Cldn16, and Cldn19 (under a high-Ca2+ diet, additionally Cldn14^39^) and are involved in paracellular transport of divalent cations (such as Ca^2+^ and Mg^2+^)^39,41^.

Even though the exact mechanism of the mosaic TJs in TAL could not be completely elucidated, S. Milatz and her colleagues concluded that at least two distinct cell types in the region are responsible for generating such patterns^39^. Based on cytosolic Cldn10b protein expression, two distinct cell populations, Cldn10b+ (77%) and Cldn10b– (23%) cells, could be identified in the cortical region of thick ascending limb^39^. However, since intracellular expression of the Cldn16 protein is mostly absent in mouse cells, the authors could not classify cell populations based on protein expression of *Cldn16*^39^. Previously, it has been reported that there are at least two morphologically distinct cell populations^42^ (i.e., rough-surfaced and smooth-surfaced cells) in the region of the thick ascending limb where the mosaic pattern of tight junctions can be observed^39^. Furthermore, recent short-read single-cell RNA-seq studies identified two distinct cell types that express either *Cldn16* alone or together with *Cldn10* in mouse^43^ and human^44^, adding more strength to the hypothesis that interactions between distinct TAL cell types generate the mosaic pattern of tight junctions in the cortical region of TAL.

#### Supplementary Note 5: An improved model for explaining the location-specific formation of mosaic tight junctions in the thick ascending limb of the loop of Henle

NKCC2-F has the lowest binding affinities for Na^+^, K^–^, and Cl^–^ ions^45,46^ and is abundantly expressed in the portion of TAL in the inner stripe of the outer medulla (ISOM)^47^ where Cl^–^ ion concentration is the highest and virtually all tight junctions are composed of Cldn10b^39^ with no permeability for divalent cations^39,41^. On the other hand, NKCC-B, which has the highest binding affinities for Na^+^, K^–^, and Cl^–^ ions^45,46^, is most abundantly expressed in the cortical portion of TAL^47-49^ where Cl^–^ ion concentration is the lowest and the most extensive mosaic patterns of tight junctions composed of either Cldn10b alone or Cldn3/Cldn16/Cldn19 are observed^39^ with high permeabilities for both monovalent cations and divalent cations^39,41^. Lastly, NKCC-A, with intermediate binding affinities for Na^+^, K^–^, and Cl^–^ ions, is expressed in both cortical and medullary parts of TAL^47-49^ but most abundantly expressed at the portion TAL in the outer stripe of the outer medulla (OSOM)^47,48^; based on the mRNA expression, mRNAs encoding NKCC-A appears to be the most predominant *Slc12a1* mRNAs across the entire length of TAL^48^. Consistent with the previous observations^50^, we observed MD cells expressing both NKCC2-B and NKCC2-A isoforms (**Figure 5e**). However, the expression of NKCC isoforms at the resolution of individual TAL cells has not been previously characterized. Our splicing atlas accurately captured the heterogeneity of full-length transcriptomes of TAL cells, clearly showing that expression of each NKCC isoform is primarily restricted to each of the three TAL cell populations in a mutually exclusive manner; NKCC-F/A/B isoforms exclusively expressed in the *Cldn10*+/*Cldn10*+ *Cldn16*+/*Cldn16*+ TAL cell populations, respectively. Consequently, given the previously known spatial locations of each NKCC isoform along the TAL, the locations of the three TAL cell populations were determined. The *Cldn10*+ TAL cell population represents a subset of TAL cells in the ISOM, while the *Cldn16*+ TAL cell population represents a subset of cortical TAL cells; the *Cldn10*+ *Cldn16*+ TAL cell population represents TAL cells in all regions of TAL, most abundant in the OSOM region.

Based on the existing characterizations of TAL cells using short-read single-cell RNA-Seq^43,44,51^ and the cellular and functional heterogeneity of the TAL cells revealed by Ouro-Seq in the current study, we suggest an improved model simultaneously explaining the location-specific formation of mosaic tight junctions and functional specializations of epithelial cells along the medulla-cortex axis in the thick ascending limb. First, the *Cldn10*+ TAL cells, lacking expression of *Cldn16*, primarily represent the TAL cells in the inner stripe of the outer medulla, which can be largely identified as the “M-TAL” (the medullary thick ascending limb) cluster from a recent large-scale single-cell atlas of human kidneys^51^. These cells express NKCC2-F, the most appropriate isoform for the higher concentrations of Na^+^, K^–^, and Cl^–^ ions in the tubular fluid in the region. Consequently, the *Cldn16*+ TAL cells, lacking expression of *Cldn10*, represent the Cldn10b-negative TAL cells described in the cortical region (~25% of the cortical TAL cells) and, to a lesser extent, in the OSOM region of TAL. These cells express NKCC2-B, the most appropriate isoform for the lower concentrations of Na^+^, K^–^, and Cl^–^ ions in the tubular fluid in the region. Most importantly, the *Cldn10*+ *Cldn16*+ TAL cell population, expressing NKCC2-A with intermediate binding affinities for Na^+^, K^–^, and Cl^–^ ions, functions as a keystone cell population that creates mosaic patterns of tight junctions in the cortex and OSOM regions. In the cortex and OSOM regions, when a *Cldn10*+ *Cldn16*+ TAL cell contacts a *Cldn16*+ TAL cell (or when two *Cldn16*+ TAL cells make contact with each other), a tight junction devoid of Cldn10b is formed^39^, while when two *Cldn10*+ *Cldn16*+ TAL cells make contact, a tight junction consisting only of Cldn10b is formed^39^, creating the mosaic patterns of two distinct tight junctions, capable of paracellular transports of both monovalent and divalent cation ions. Conversely, in the ISOM region, when a *Cldn10*+ TAL cell makes contact with another *Cldn10*+ TAL cell or less frequently, a *Cldn10*+ *Cldn16*+ TAL cell (or when two *Cldn10*+ *Cldn16*+ TAL cells make contact with each other), a tight junction solely composed of Cldn10b is formed, resulting in exclusive usage of Cldn10b tight junctions^39^, impermeable to divalent cations^39,41^. The spatial confinement of the *Cldn16*+ TAL cells mainly in the cortical region and the inability of Cldn10b to form a heterophilic interaction with any other claudin proteins expressed in TAL cells^39^ (such as Cldn3, Cldn16, and Cldn19) also constitute other key mechanisms by which mosaic tight junctions are restricted to the cortical and OSOM regions.

Our single-cell full-length transcriptome analysis using Ouro-Seq also identified various *de novo* isoforms expressed in the TAL cells. For instance, we detected the usage of an alternative 5’ splice site, resulting in the inclusion of 45 bp-shorter cassette exon B (the red highlighted box, **Figure 5h**) in a novel *Slc12a1* transcript, encoding a variant of NKCC2-B isoform that is precisely 15 amino acids shorter. Additionally, we identified the novel *Fxyd2* isoform most abundantly expressed by the *Cldn16*+ TAL cell population (**Extended Data Figure 21b**). Mutations in FXYD2 (the human homolog of Fxyd2) have been linked to dominant hypomagnesaemia^52^, and it is a known downstream target of Hnf1b, a transcription factor responsible for renal tubule segmentation during kidney development. Also, Hnf1b has been described as a key transcription factor determining the identity of the *Cldn16*+ TAL cell population in a recent study^44^, possibly explaining the differential transcript expression of *Fxyd2* among distinct TAL cell populations.

In summary, using Ouro-Seq, we characterized full-length transcriptomes of distinct cell populations in the thick ascending limb, linking the unique NKCC2 isoform expression patterns to each TAL cell population, identifying an essential role of the *Cldn10*+ *Cldn16*+ TAL cell population for regulating paracellular transports in TAL.

#### Supplementary Note 6: *De novo* promoters of *Top1mt*

*Top1mt* encodes a mitochondria-specific topoisomerase, which recently emerged during vertebrate evolution^53^, resulting from the duplication and subsequent diversification of an ancestral gene that has become the *Top1* gene, encoding one of the five nuclear topoisomerases in vertebrates. Even though *Top1mt* is considered “non-essential” for mouse development due to the presence of other topoisomerases of type II and type III in mitochondria^54^, it appears essential for mitochondrial homeostasis^55^, especially during regeneration^56^ and adaptation to biological and chemical stresses^57^.

Currently, there is only a single annotated transcript for *Top1mt*, the first 5’ exon of which contains the start of a protein-coding sequence (CDS); any alternative promoter usage events will result in the inclusion of a novel 5’ exon, encoding a distinct protein isoform of *Top1mt* or generating an mRNA transcript that does not encode any functional protein. Podocytes and late spermatids mostly lack expression of the previously annotated *Top1mt* transcript, indicating that the known Top1mt protein encoded by the sole annotated transcript will not be expressed in these cell types. For the remaining cell types (~200 cell types) characterized in our splicing atlas, only the annotated *Top1mt* transcript was expressed, starting at the known transcript start site. One exception was natural killer cells, which expressed a low but detectable fraction of the novel *Top1mt* transcript that starts from one of the newly discovered promoters utilized by podocytes (**Extended Data Figure 22a**).

Since the basement membrane of glomerular microvasculature only partially envelops the glomerular endothelial cells, the high pressure of the afferent arteriole is directly transmitted to intraglomerular mesangial cells and eventually podocytes^58^, which collectively envelopes the glomerular basement membrane enveloping the rest of the mesangium. Podocytes monitor and adaptively respond to the constantly changing pressure of the arteriole, utilizing various cytoskeletal structures and contractile systems^58^ (**Extended Data Figure 6d**), which require high energy consumption and, thus, efficient mitochondrial respiration. Consequently, mature mouse podocytes are rich in mitochondria and rely heavily on mitochondrial respiration to meet the energy requirements (accounting for 77% of the total cellular respiration)^59^. Therefore, the exclusive usage of novel *Top1mt* transcripts in mouse podocytes implies that these novel *Top1mt* transcripts have certain biological functions related to mitochondrial homeostasis in podocytes.

### SUPPLEMENTARY METHODS

*Ouro-Seq*

**Synthesis of sgRNA for Ouro-Seq**

For targeted linearization of circular DNAs using CRISPR-Cas9 nuclease, single guide RNAs (sgRNAs) were synthesized using EnGen® sgRNA Synthesis Kit, *S. pyogenes* (#E3322, New England Biolabs) following the recommendations of the manufacturer’s protocol with a few modifications. To increase the sgRNAs yield, reaction time for *in vitro* transcription was increased from 30 minutes to 2 hours. Importantly, DNase I treatment was omitted after the in vitro transcription reaction to prevent the carry-over of DNA endonuclease. The synthesized sgRNA was purified by using Monarch RNA Cleanup Kit (#T2040, New England Biolabs), quantified with NanoDrop Microvolume Spectrophotometers (Thermo Fisher Scientific), diluted with nuclease-free water (50 ng/µl for targeted depletion and 10 ng/µl for targeted enrichment and digestion of intramolecular-ligation-junctions), aliquoted into separate tubes for individual reactions, and stored at -20 °C until the Ouro-Seq experiments.

**Preparation of a Y-shaped adapter**

The resulting linear single-cell cDNA library from Ouro-Enrich can be directly used for long-read sequencing experiments by ligating sequencing adapters of the long-read sequencing platforms. Alternatively, a Y-shaped adapter can be ligated to the linear cDNA molecules, introducing PCR handles at both ends of the molecules, which enables subsequent re-amplification using PCR. The following steps can be used to prepare Y-shaped adapters. First, the oligo annealing buffer is prepared by adding 0.1 mL of 5 M NaCl to 9.9 mL of TE buffer (10 mM Tris, 1 mM EDTA, pH 8.0). Next, the oligo annealing reaction mixture is assembled by mixing the following components: 98 µl of the oligo annealing buffer, 1 µl of 100 µM “Y-shaped-adapter-F” oligo, and 1 µl of 100 µM “Y-shaped-adapter-R” oligo (**Extended Data Table 3**). The oligo annealing reaction mixture is then heated to 95 °C for 3 minutes using a thermocycler and slowly cooled to room temperature, using the ramp rate of -0.1 °C/second. The resulting reaction mixture contains 1 µM of the hybridized Y-shaped adapter, which is diluted to 0.5 µM with nuclease-free water and stored at -20 °C until the Ouro-Enrich experiments.

**Targeted depletion of uninformative cDNAs with Ouro-Deplete**

For additional depletion of uninformative cDNA molecules, the self-cyclization reaction mixture is subjected to a CRISPR-Cas9-based digestion reaction that linearizes the circular target molecules in the reaction mixture. Subsequently, exonucleases remove the linearized target molecules, cell barcode-free artifacts, and cell barcode-doublet artifacts from the library during the exonuclease treatment. First, Cas9-sgRNA ribonucleoprotein complexes (RNPs) are assembled by mixing the following reagents and incubating the reaction mixture at room temperature for 10 minutes: 7 µl of nuclease-free water, 9.5 µl of 50 ng/µl sgRNA targeting unwanted cDNA molecules, 2 µl of 10x NEBuffer r3.1 (#B6003S, New England Biolabs), and 0.5 µl of 20 µM SpCas9 nuclease (#M0386T, New England Biolabs). Next, the assembled Cas9-sgRNA RNPs (20 µl) are added to the self-cyclization reaction mixture (10 µl), which is then incubated at 37 °C for 30 minutes for targeted linearization of uninformative cDNA molecules. For cleanup, 1 µl of thermolabile Proteinase K (#P8111S, New England Biolabs) is added to the reaction mixture, which is incubated at 37 °C for 15 minutes. The thermolabile proteinases are heat-inactivated by incubating the reaction mixture at 55 °C for 10 minutes. From this point, the remainder of the protocol is identical to that of Ouro-Seq. After the exonuclease treatment and self-cyclization reversal reaction, the resulting single-cell cDNA library will be depleted of uninformative cDNAs and artifact molecules.

The sequencing efficiency of single-cell long-read RNA-seq is often significantly reduced by the presence of uninformative yet highly abundant cDNA molecules in the library, which are derived from mitochondrial DNA (mtDNA) transcripts, ribosomal RNA (rRNA) (rRNAs lack poly(A) tail, but has internal priming sites), and other highly abundant transcripts (often constitute ambient RNAs) such as hemoglobin mRNA (released when red blood cell ruptures). These cDNA molecules are often very short (~600 bp) and more efficiently amplified by PCR than other cDNA molecules (1–6 kbp). As a result, these uninformative molecules often constitute 30%–70% of a single-cell full-length cDNA library, decreasing sequencing output for other informative cDNA molecules.

To effectively deplete cDNAs from mitochondrial DNA and rRNAs from the single-cell RNA sequencing cDNA library with Ouro-Deplete, we designed sgRNAs targeting 15 and 26 sites of mtDNA and the ribosomal RNA gene (rDNA) of human and mouse (synthesized using the “sgRNA.mouse_mtDNA_rRNA_26” and “sgRNA.human_mtDNA_rRNA_15” oligos in **Supplementary Table 3**), respectively. Additionally, the depletion efficiency can be further increased by including sgRNAs targeting unwanted cDNAs that evade the depletion process. In the current study, after applying Ouro-Deplete using the sgRNAs, the proportion of molecules derived from mtDNA and rRNA dropped below 10% for most samples.

Existing methods for depleting unwanted target molecules from the single-cell sequencing library, such as scDASH^60^, require PCR amplification because these methods merely prevent the amplification of target molecules by separating the 3’ PCR handle from the 5’ PCR handle of the target molecule. However, exonucleases completely degrade the linearized target molecules in Ouro-Deplete, making the subsequent PCR amplification optional.

**Rapid high-throughput targeted enrichment with Ouro-Enrich**

In Ouro-Seq and Ouro-Deplete, the self-cyclization reversal reaction linearizes every circular DNA molecule in the purified circular cDNA library. In Ouro-Enrich, a purified circular cDNA library from Ouro-Seq or Ouro-Deplete is subjected to targeted linearization with CRISPR-Cas9 nuclease to enrich target molecules. First, Cas9-sgRNA ribonucleoprotein complexes (RNPs) are assembled by mixing the following reagents and incubating the reaction mixture at room temperature for 10 minutes: 7 µl of nuclease-free water, 9 µl of 10 ng/µl sgRNA targeting cDNA molecules of interest, 2 µl of 10x NEBuffer r3.1 (#B6003S, New England Biolabs), and 2 µl of 1 µM SpCas9 nuclease (#M0386S, New England Biolabs). Subsequently, the assembled Cas9-sgRNA RNPs (20 µl) and 1 µl of 10x NEBuffer r3.1 are added to 9 µl of the purified circular DNA library (from Ouro-Seq or Ouro-Deplete) dissolved in nuclease-free water. The targeted linearization reaction mixture (30 µl) is then incubated at 37 °C for 25 minutes to digest target molecules. For cleanup, 1 µl of Proteinase K (#P8107S, New England Biolabs) is added to the targeted linearization reaction mixture, incubated at room temperature for 10 minutes, purified with 1.2x SPRIselect beads, and resuspended in 10 µl of nuclease-free water to yield a selectively linearized cDNA library, which can be directly used as inputs for targeted long-read sequencing experiments; the remaining non-target circular cDNA molecules in the library are not sequenced when ligation-based library preparation methods are used since sequencing adapters cannot be ligated to circular DNA molecules.

For re-amplification of the linearized target molecules, the library is end-prepped and ligated to Y-shaped adapters. First, the dA-tailing reaction mixture is assembled by adding the following components: 10 µl of the selectively linearized cDNA library, 2.5 µl of 10x NEBuffer 2 (#B7002S, New England Biolabs), 0.5 µl of 5mM dATP (diluted in nuclease-free water, #N0440S, New England Biolabs), 2 µl of Klenow fragment (3’→5’ exo-) (#M0212S, New England Biolabs), and 10 µl of nuclease-free water. The dA-tailing reaction mixture (25 µl) is incubated at 37 °C for 30 minutes, purified with 1.2x SPRIselect beads, and resuspended in 10 µl of nuclease-free water. Next, 1 µl of the hybridized Y-shaped adapter (0.5 µM; prepared using the protocol described above) and 11 µl Blunt/TA Ligase Master Mix (#M0367S, New England Biolabs) are added to the 10 µl of dA-tailed selectively linearized cDNA library. The adapter ligation reaction mixture is then incubated at room temperature for 30 minutes, purified with 1x SPRIselect beads, and resuspended in nuclease-free water. Lastly, the adapter-ligated linearized cDNA library is amplified for 5–12 cycles with the primers “Y-shaped-adapter-primer-F” and “Y-shaped-adapter-primer-R” (**Extended Data Table 3**) and purified with 1.2x SPRIselect beads.

For the targeted enrichment of cDNA molecules transcribed from the 11 mouse mtDNA-encoded genes (*mt-Atp6*, *mt-Atp8*, *mt-Cytb*, *mt-Co1*, *mt-Co2*, *mt-Co3*, *mt-Nd1*, *mt-Nd2*, *mt-Nd3*, *mt-Nd4*, and *mt-Nd5*) with Ouro-Enrich, we synthesized sgRNAs using the subset (“sgRNA.mouse_mtDNA_gene_11” oligos in **Supplementary Table 3**) of the oligos utilized for the targeted depletion of the cDNA molecules from mtDNA and rDNA with Ouro-Deplete. For the targeted enrichment of cDNA molecules covering 13 single nucleotide variants (SNVs) of interest detected from the single-cell cDNA library^61^ of a patient with relapsed AML (sample# AML02-Rel), sgRNAs were synthesized using the “sgRNA.human_BM-MNC_AML02-CR_genomic-loci-with-SNV_22” oligos (**Supplementary Table 3**).

Ouro-Enrich enables high-throughput enrichment of full-length cDNA molecules containing specific target sequences from a single-cell cDNA library while simultaneously depleting cell barcode-free and -doublet artifacts from the library. However, by including the CRISPR-based targeted depletion step of Ouro-Deplete, Ouro-Enrich can be utilized as a complex search operator by only retrieving the molecules matched with specific exon or splice junction usage patterns from a single-cell cDNA library, such as cDNA molecules that including a specific exon “A” while not including specific exon “B” or “C,” in a highly parallel way. It is also possible to utilize the logical AND operators to enrich specific cDNA molecules from a single-cell cDNA library by consecutively applying two or more Ouro-Enrich experiments to the library. Also, Ouro-Enrich, similar to Ouro-Seq, has a minimal bias for different molecule sizes, allowing a consecutive run of multiple Ouro-Enrich experiments without applying additional size selection to compensate for the length-bias of the process. Consequently, Ouro-Enrich enables a high-throughput selection of cDNA molecules matched with specific splicing patterns. Overall, Ouro-Enrich provides a highly flexible and versatile target enrichment framework and complements existing target capture techniques.

**Cost-effective full-length transcriptome analysis for the cells of interest with Ouro-Cell-Enrich**

Previously, enrichment of molecules containing specific cell barcodes from a pooled single-cell library was only possible through PCR with a set of primers capturing the barcodes, which is low throughput and has a significant risk of “re-barcoding,” possibly capturing a wrong cell barcode that contains a few mismatches through a mispriming event and eventually masking the mismatches in the output library. Recently, a hybridization capture-based method was developed to enrich molecules of a few specific cell barcodes (~5 cells) without the risk of re-barcoding, utilizing locked nucleic acid-containing (LNA) hybridization probes that are ~20 bp in length^62^. However, the hybridization-based method requires an overnight reaction and has low throughput due to a high cost per target cell barcode.

Ouro-Enrich can be utilized for the enrichment of molecules containing specific cell barcodes from various pooled single-cell (spatial) transcriptomics and genomics libraries^10^, including 10x Genomics single-cell and spatial transcriptomics products, by selecting an appropriate protospacer adjacent motif (PAM) sequence (or introducing one by performing PCR with a primer with a desired substitution) around the cell barcode sequence. With this specific implementation of Ouro-Enrich, which we named Ouro-Cell-Enrich, targeted enrichment of 50–100 cells can be achieved for 120–200 USD (Oligo Pools, Integrated DNA Technologies). Currently (as of May 2024), at least 1,000 USD is required for characterizing full-length transcriptomes of one single-cell cDNA library containing thousands of cells. However, researchers are often interested in a specific subset of cells or want to subsample the cells during the exploratory phase of the research project, where Ouro-Cell-Enrich can be utilized.

Using Ouro-Cell-Enrich, full-length cDNA molecules belonging to a total of 778 cells were enriched from 5 pooled single-cell cDNA libraries: 19 and 8 HSC cells from the AML02-CR and AML02-Rel samples^61^ using the “sgRNA.CB.human_BM-MNC_AML02-CR_HSC” and “sgRNA.CB.human_BM-MNC_AML02-Rel_HSC” oligos, respectively; 301 and 115 LMPP cells from the AML02-CR and AML02-Rel samples using the “sgRNA.CB.human_BM-MNC_AML02-CR_LMPP” and “sgRNA.CB.human_BM-MNC_AML02-Rel_LMPP” oligos, respectively; 20 LSC cells from the AML02-CR sample using the “sgRNA.CB.human_BM-MNC_AML02-CR_LSC” oligos; 42 granulocytes and 23 cardiomyocytes from the mHeart sample using the “sgRNA.CB.mouse_heart_Cardiomyocyte+Granulocyte” oligos; 99 myofibroblasts from the mHeart sample using the “sgRNA.CB.mouse_heart_Myofirboblast” oligos; 53 stellate cells from the mLiver sample using the “sgRNA.CB.mouse_liver_Stellate_cell” oligos; 46 hepatocytes and 7 cholangiocytes from the mLiver sample using the “sgRNA.CB.mouse_liver_Hepatocyte+Cholangiocyte” oligos; 45 ALOH cells from the mKidney sample using the “sgRNA.CB.mouse_kidney_ALOH” oligos (all oligos were purchased as Oligo Pools from Integrated DNA Technologies) (**Supplementary Table 3**). As a result, we obtained the robust enrichment of the target cells across the samples for widely varying numbers of target cells, with an average enrichment accuracy of ~90% (**Figure 2g**). When less than a couple dozen cells (~10 cells) were targeted, we observed generally low enrichment accuracy (~75%), which can be attributed to Cas9 off-target cleavage of cDNA molecules containing similar cell barcodes with a few mismatches, which may be improved by the use of the high-fidelity CRISPR-Cas9 variants in the Ouro-Cell-Enrich experiments.

**Robust size selection with Ouro-SizeSelect**

Largely, there are two types of methods for the separation of DNA molecules according to their sizes and secondary structures: electrophoresis-based (e.g., agarose gel electrophoresis) and precipitation-based (e.g., silica spin-column or carboxylated magnetic beads). Initially, we employed an electrophoresis-based automated size-selection system (BluePippin, Sage Science) to enrich longer cDNA molecules from a single-cell full-length cDNA library. However, we observed that even a minute amount of PCR-bubble^63^ artifacts in a PCR product can significantly interfere with electrophoresis-based DNA size-selection methods. Moreover, the exact amount of PCR-bubbles can be highly variable across individual samples and PCR conditions. Therefore, we developed Ouro-SizeSelect, a precipitation-based DNA size-selection method that can tolerate higher levels of PCR-bubbles by employing a series of sequential precipitation-based DNA size-selection steps with increasing size cutoff (currently, total 8 steps with 0.9, 1.25, 1.75, 2.25, 2.75, 3.25, 3.75, and 4.5 kbp cutoffs, respectively). Because Ouro-SizeSelect is based on solid-phase reversible immobilization (SPRI) beads, it is faster, more automation-friendly, and more scalable than electrophoresis-based methods.

First, a stock solution (10ml) for precipitation-based DNA size-selection (“size-select solution”) is prepared by thoroughly mixing the reagents in the following order: 3.75 ml of nuclease-free water (#AM9932, Invitrogen), 3.2 ml of 5 M NaCl RNase-free) (#AM9759, Invitrogen), 100 µl of 1 M Tris-HCl (RNase-free), pH 8 (#AM9855G, Invitrogen), 20 µl of 0.5 M EDTA (RNase-free), pH 8 (#AM9260G, Invitrogen), 200 µl^64^ of 10 % (v/v) solution of Tween-20 (#28320, Thermo Scientific), and lastly, 2.75 ml of 40% (w/w) solution of PEG 8000 (#P1458-50ML, Sigma-Aldrich). The size-select solution can be aliquoted into separate nuclease-free conical tubes and stored at 4-10 °C until the Ouro-SizeSelect experiments. The first step of Ouro-SizeSelect is the depletion of cDNA molecules smaller than 0.9 kbp using 0.5x SPRIselect beads (**Extended Data Figure 1c**) using the manufacturer’s protocol. Subsequently, sequential size-selection steps using 0.9x, 0.8x, 0.75x, 0.725x, 0.7x, 0.69x, and 0.68x size-selection solutions with size cutoffs at 1.25, 1.75, 2.25, 2.75, 3.25, 3.75, and 4.5 kbp are performed, respectively (**Extended Data Figure 1c**). For each size-selection step, the resulting library can be either directly subjected to the next size-selection step or optionally re-amplified using PCR.

A typical size-selection step utilizing the size-selection solution is described as follows (“0.9x” size-select solution as an example). First, a “0.9x” size-select solution is prepared by adding 90 µl (hence “0.9x”) of the size-select solution to 100 µl of nuclease-free water, mixed thoroughly with vortexing. Next, in a 0.2ml nuclease-free PCR tube (#1402-4700, USA Scientific), 10–40ng of an input single-cell cDNA library is diluted to 50 µl using nuclease-free water. Subsequently, 50 µl of SPRIselect solution (containing carboxylated magnetic beads) is added to the library, mixed, and (after a brief spin-down) incubated at room temperature for 3 minutes. The magnetic beads are then pelleted using a magnetic stand. After discarding the supernatant, the tube is removed from the magnetic stand. Next, the pelleted beads are resuspended in 50 µl of the 0.9x size-select solution and incubated at room temperature for 5 minutes with vigorous vortexing for 10 seconds every 2 minutes. After a brief spin-down, the beads are pelleted by placing the tube on a magnetic stand. The wash process using the 0.9x size-select solution is repeated twice (a total of three washes). After discarding the supernatant, the pelleted beads are washed with 80% ethanol and dried for up to 60 seconds. After removing the tube from the magnetic stand, the pelleted beads are resuspended in 30 µl of nuclease-free water, incubated at room temperature for 2 minutes, and pelleted again by placing the tube on the magnetic stand. The supernatant containing size-selected cDNA molecules is then collected (and, optionally, re-amplified for 3–5 cycles). The resulting size-selected library is quantified with the Qubit dsDNA high-sensitivity assay (#Q33231, Invitrogen) using the Qubit 4 Fluorometer (Invitrogen). It is recommended that 8 samples (12 samples) be processed together using a PCR 8-tube (12-tube) strip and multichannel pipettes for better efficiency.

Since PCR-bubbles are generated during the annealing stage of a PCR cycle, especially when the ratio of primer to template is low (e.g. when over-amplification occurs), the following PCR conditions were used throughout the size-selection process to minimize generation of PCR-bubble artifacts in the amplified product: up to 40 ng of an input cDNA library in each 100 µl PCR mixture (using Q5® Hot Start High-Fidelity 2X Master Mix, #M0494S, New England Biolabs) for 5 cycles of amplification (for a different number of cycles, the maximum amount of input is modified accordingly; e.g., up to 10 ng and 20 ng of an input library for 6 and 7 cycles of amplification, respectively); for the initial amplification, 0.4 µM (final concentration) of the primers “part-R1-PBK004F” and “part-TSO-PBK004R” (**Extended Data Table 3**) were utilized for tagging cDNA molecules with additional PCR handles (initial denaturation: 98 °C for 3 minutes; cycles: 98 °C for 30 seconds, 63 °C for 30 seconds, 72 °C for 3 minutes; final elongation: 72 °C for 3 minutes); all subsequent amplification utilized 0.4 µM (final concentration) of the primers “PBK004F” and “PBK004R” (**Extended Data Table 3**) (initial denaturation: 98 °C for 3 minutes; cycles: 98 °C for 30 seconds, 63 °C for 30 seconds, 72 °C for 3 minutes; final elongation: 72 °C for 3 minutes).

After all the sequential size-selection steps of Ouro-SizeSelect are completed, the size-selection results are evaluated using a gel-electrophoresis-based method, such as the High Sensitivity DNA assay (#5067-4626, Agilent) using the Bioanalyzer 2100 (Agilent) or the agarose gel-electrophoresis using SYBR™ Safe DNA Gel Stain (#S33102, Invitrogen). Finally, for a long-read sequencing experiment, the size-selected libraries are pooled with the following weight ratio (the numbers are in parentheses, followed by the expected size range): the original artifact-depleted library (10 ng, 0.25–1.5 kbp), the size-selected libraries after 0.5x SPRIselect (40 ng, 0.75–2 kbp), 0.9x (40 ng, 1.25–2.5 kbp), 0.8x (40 ng, 1.75–3 kbp), 0.75x (20 ng, 2.25–3.5 kbp), 0.725x (20 ng, 2.75–4 kbp), 0.7x (20 ng, 3.25–4.5 kbp), 0.69x (20 ng, 3.75–5.25 kbp), and 0.68x (20 ng, 4.5–6 kbp) size-select solutions (**Extended Data Figure 1c**).

After applying Ouro-SizeSelect, the proportion of uninformative cDNAs from mtDNA and rRNA often decreases to negligible levels (below 2%) since these uninformative cDNAs are often smaller than 1 kbp and over-represented in a single-cell cDNA library before size-selection due to the PCR amplification bias.

*Ouro-Tools*

The Ouro-Tools pipeline comprises five main modules, allowing seamless integration with existing bulk and single-cell long-read RNA-seq pipelines and tools (**Supplementary Figure 1**). Every main module of Ouro-Tools utilizes efficient parallelization for compute-intensive tasks to facilitate the processing of large datasets. Additionally, each Ouro-Tools module employs filesystem-based locks for parallel processing of a large number of samples across multiple machines for scalability. This detailed description of the Ouro-Tools pipeline is based on the Ouro-Tools version v0.1.1, which was utilized for building the mouse single-cell long-read splicing atlas (v202405) and the benchmarking of previously published long-read scRNA-seq datasets.

***The raw long-read pre-processing module***

As the first module of the Ouro-Tools pipeline (implemented as the “LongFilterNSplit” workflow, **Supplementary Figure 2**), the module has a dual function for (1) providing comprehensive quality control metrics of a long-read scRNA-seq experiment and (2) pre-processing of raw long-read sequencing data for the downstream analysis. The raw long-read pre-processing module efficiently identifies cell barcode-free and cell barcode-doublet artifacts, chimeric cDNA molecules, molecules generated by mispriming events, and uninformative cDNA molecules (e.g., mtDNA-derived and rRNA-derived cDNA molecules), by aligning individual reads (and parts of each read) against the reference genome and the predefined sequences of uninformative cDNA molecules. First, each long-read sequence is searched against the uninformative cDNA sequences to detect and exclude an unwanted cDNA molecule from the subsequent analysis. Next, the long-read sequence is searched against the reference genome, which is more computationally intensive than searching the uninformative cDNA sequences, to detect and exclude a molecule that does not originate from the genome (e.g., adapter concatemers, such as adapter dimers). For the construction of the mouse splicing atlas and the benchmarking of various long-read scRNA-seq methods, the following sequences were utilized: GRCh38, GRCm38, TAIR10, PvP01^65^ (downloaded from the NCBI using the GenBank accession ‘GCA_900093555.2’), Macaca_fascicularis_6.0 were utilized as the reference genomes for *Homo sapiens*, *Mus musculus*, *Arabidopsis thaliana*, *Plasmodium vivax*, and *Macaca fascicularis*, respectively; ‘NR_046235.3’ and ‘NR_046233.2’ (GenBank accessions) sequences were utilized for detecting the cDNAs originated from rRNAs for *Homo sapiens* and *Mus musculus*, respectively; the mitochondria genome and (if applicable) the chloroplast genome sequences that are included in the reference genome were utilized for detecting the cDNAs originated from the chloroplast and mitochondrial genomes.

Subsequently, the read aligned to the reference genome is further analyzed to identify and split a composite molecule generated by *in vitro* and *in silico* concatenation events^66^ of two or more cDNA molecules with intact adapters (hence, the name of the workflow “LongFilterNSplit”). Internally, Ouro-Tools employs the Minimap2 aligner^67^ (using mappy, the Python interface to Minimap2) for the splice-aware alignment of a long-read sequence to the reference genome. After aligning a read against the reference genome, Minimap2 returns more than one confident alignment if individual parts of the read can be confidently aligned to different regions or strands (e.g., + and - strands) across the reference genome. If the length of an intervening sequence between the two adjacent independently aligned parts of the read is longer than the threshold (the default setting is 150 bp, considering the length of a nanopore sequencing adapter), the two adjacent parts are split into separate cDNA molecules (the intervening sequence is copied and attached to each molecule), both of which are classified as a non-chimeric cDNA molecule. If the length of the intervening sequence is shorter than the threshold or there is no intervening sequence between the two adjacent independently aligned parts of the read, both parts of the read are collectively flagged as belonging to the same chimeric molecule. However, we noticed that the Minimap2 aligner sometimes reports two or more fragmented alignments for a non-chimeric cDNA molecule when the transcript contains an extremely large intron (> 50–100 kbp). Therefore, when the two adjacent “independently” aligned parts of a presumed chimeric molecule are aligned to the same strand of the same chromosome of the reference genome without an intervening sequence and the distance between the two aligned regions on the reference genome is shorter than the given threshold (the default setting is 200 kbp), the two alignments are combined into a single alignment, flagged as a non-chimeric cDNA molecule.

For a long-read scRNA-seq experiment that utilized Ouro-Enrich for targeted enrichment of the transcripts of interest, the raw long-read pre-processing module searches each read aligned to the reference genome for the sign of a topological modification introduced by Ouro-Enrich (after an Ouro-Enrich experiment, an inverted transposition occurs around the target site, see **Figure 2a**). With the default setting, when the two adjacent “independent” alignments of a read are situated next to each other in the reference genome (the distance between the alignment ends < 30 bp) and satisfy the conditions of a topological modification specific to Ouro-Enrich, the original cDNA molecule is reconstructed from the read by removing the adapter sequences ligated to the cut site and reverse the topological modification from the Ouro-Enrich experiment.

After each unique cDNA molecule is identified and classified into a chimeric or non-chimeric cDNA molecule, the soft-clipped sequences and the short sequences at the ends of the genome-aligned portion of the molecule (the default setting is 36 bp) are collected and analyzed for identification of cell barcode-free and cell barcode-doublet artifacts and cDNA molecules generated by mispriming events. First, for each end of the cDNA molecule, the presence of an enzymatically added poly(A) tail (hereinafter “external poly(A)”) or a genome-encoded internal poly(A) tract is detected. Specifically, with the default setting, a sliding window of 16 bp is utilized to identify a region with a percentage of A (adenosine) > 75%. Next, for each end of the cDNA, the number of consecutive G (guanosine) nucleotides in the soft-clipped sequence, starting from the end of the genome-aligned portion of the cDNA, is collected. Based on the collected metrics, the cDNA molecule (either non-chimeric or chimeric) is classified into one of the following classes: “external poly(A) and external-Gs,” “external poly(A) and no external-Gs,” “internal poly(A) and external-Gs,” “internal poly(A) and no external-Gs,” “poly(A) at both ends,” and “no poly(A).” Only the non-chimeric cDNA molecules classified as the “external poly(A) and external-Gs” class are considered as the cDNA molecules that are not generated by mispriming (i.e., the *in vitro* full-length cDNA molecules) (**Figure 1g**). The non-chimeric cDNA molecules classified as “no poly(A)” and “poly(A) at both ends” classes are considered cell barcode-free and cell barcode-doublet artifacts, respectively.

Lastly, according to the classification results, all the cDNA molecules are organized into separate output FASTQ files (**Supplementary Figure 2**), each of which can be aligned to the reference genome using a long-read aligner of choice with the preferred setting (e.g., Minimap2 for annotation-guided alignment based on the transcript annotations prepared by the researcher). Particularly, for each cDNA molecule that contains a single (external or internal) poly(A) tail, the read is re-oriented so that it has the same orientation as its original mRNA transcript, with the poly(A) tail at its 3’ end; the resulting long-reads of cDNAs can be utilized for strand-specific long-read RNA-seq analysis. Additionally, for each class, the lengths of the cDNA molecules are collected as a histogram, providing a comprehensive overview and useful quality-control metrics of a long-read scRNA-seq experiment (**Extended Data Figure 2-3**).

***The barcode extraction module***

In the standard Ouro-Tools pipeline, the strand-specific long-reads of valid cDNA molecules (with the default setting, the “aligned_to_genome__non_chimeric_poly_A__plus_strand” FASTQ file from the previous module) are aligned to the reference genome using Minimap2 with gene annotations, which is shown to be highly effective for reducing the alignment errors around the known splice junctions in a recent benchmarking of various long-read RNA-seq alignment frameworks^68^. For the construction of the mouse splicing atlas and the benchmarking of various long-read scRNA-seq methods, the annotation-guided alignments were performed using the gene annotations from the Ensembl release 105, the Ensembl release 102, the Ensembl Plants release 56, and PvP01^65^ (the GenBank accession ‘GCA_900093555.2’) for *Homo sapiens*, *Mus musculus*, *Arabidopsis thaliana*, and *Plasmodium vivax.*

After directional long-reads of cDNAs (long-read cDNA sequences that are oriented in the same direction as their original mRNA transcripts, with poly(A) tails at their 3’ ends) are aligned to the reference genome, the BAM file is further processed by the barcode extraction module (implemented as the “LongExtractBarcodeFromBAM” workflow), which identifies cell barcode (CB) and unique molecular identifier (UMI) sequences for each read and exports the results as a “barcoded” BAM file, a BAM file containing corrected CB and UMI sequences for each read using the predefined SAM tags (**Supplementary Figure 3**). The workflow is primarily composed of two phases. In the first phase, the raw (uncorrected) CB and UMI sequences are extracted from the soft-clipped sequences of the reads. Subsequently, all raw CB sequences of the sample are collectively analyzed to prepare the mappings to correct cell barcode sequences. In the second phase, the raw CB sequences are corrected using the mappings. For the correction of UMI sequences, the raw UMI sequences are grouped for each barcode attachment site (i.e., polyadenylation site) on the reference genome for each corrected CB sequence and clustered using a linear-time UMI clustering algorithm inspired by a linear-time protein sequence clustering algorithm^69^ (mmseq2).

The first phase of the workflow starts with searching the 3’ and 5’ PCR handle sequences in soft-clipped sequences of individual reads. With the default setting, the workflow searches for the 3’ and 5’ PCR handles of cDNA molecules from 10x Genomics, which are partial-Read-1 (R1, “CTACACGACGCTCTTCCGATCT”) and partial-Template-Switch-Oligonucleotide (TSO, “AAGCAGTGGTATCAACGCAGAG”) primer sequences. For long-read scRNA-seq libraries prepared using Ouro-Cell-Enrich, the following 3’ PCR handle sequence should be used instead of R1, which contains parts of R1 and “Y-shaped-adapter” utilized in Ouro-Cell-Enrich): “CAGCACATCCCTTTCTCACACTAGCCTTCTCGTT.” The search results of 3’ and 5’ PCR handles for each read are stored using the ‘XR’ and ‘XT’ tags, which indicate the number of errors (nucleotide mismatches, insertions, and deletions combined) for the identified 3’ and 5’ PCR handles, respectively (-1 indicates that the PCR handle could not be identified using the maximum error rate of 20% with the default setting) (see **Supplementary Table 4** for the complete list of SAM tags utilized by the Ouro-Tools pipeline). Next, for the scRNA-seq chemistries utilizing the 3’ barcoding strategy, raw CB and UMI sequences are extracted from the soft-clipped sequence at the 3’ end of the read (where barcodes are attached) based on the identified location of the 3’ PCR handle (vice versa for the scRNA-seq chemistries utilizing 5’ barcoding strategy), which are stored using the ‘CU’ tag (uncorrected raw CB-UMI sequence). For parallelization (faster processing), since searching the PCR handles can be performed independently for each read, reads are randomly distributed across worker processes, each exporting the search results as a temporary BAM file. Once all worker processes complete the tasks, the temporary BAM files from the worker processes are sorted and combined into a single temporary BAM file for the next phase of the workflow.

Subsequently, the raw CB-UMI sequences are collected from all workers to identify a list of valid cell barcode sequences for the sample. First, the raw CB sequences are filtered using the barcode whitelist (the barcode whitelist for the 10x Genomics 3’ v3.1 scRNA-seq product), the minimum count (observed in at least 3 different cDNA molecules), and the minimum rank (up to 20,000 whitelist barcodes with highest counts are allowed for the downstream processing) with the default setting. Next, two kinds of mappings are constructed for efficient correction of raw cell barcode sequences. The primary mapping accounts for a single error in one of the valid cell barcodes (a mismatch of a single nucleotide, a single insertion, or a single deletion), enabling efficient correction of a raw cell barcode sequence that harbors a single sequencing error, which accounts for the majority of raw cell barcode sequences that require error correction. During the construction of the primary mapping, a collision between two or more valid cell barcodes is recorded and eventually excluded from the mapping. The secondary mapping utilizes varying lengths of k-mers that are unique to each valid cell barcode for the correction of raw cell barcode sequences containing a higher number of errors. With the default setting, the secondary mapping is constructed by generating k-mers from each valid cell barcode sequence (k = 8–15) while marking the collided k-mers that could be generated from more than one valid cell barcode.

In the second phase of the workflow, the temporary BAM file containing PCR handle search results is further analyzed. For efficient clustering of raw UMI sequences of the cDNA molecules sharing the same barcode attachment site on the reference genome, the tasks are split into batches for individual chromosomes for parallelization (all reads aligned to the same chromosome are analyzed together by a single worker process). First, a raw cell barcode sequence is compared with the barcode whitelist to check whether the raw CB contains any sequencing error. If the raw CB contains at least one sequencing error, the primary mapping is utilized to check whether only a single sequencing error is present in the raw CB. If the raw CB appears to contain more than one sequencing error, the secondary mapping is then utilized to correct the raw CB; longer k-mers are utilized first to correct smaller numbers of errors; if the error correction process fails even using the shortest k-mers (k=8, with the default setting), the raw CB (and the cDNA molecule containing the barcode) is discarded and excluded from the downstream analysis. The resulting error-corrected cell barcode sequence is stored using the ‘CB’ tag.

Next, the raw UMI sequences are grouped for each barcode attachment site on the reference genome for each corrected CB sequence. The grouped raw UMI sequences are then clustered using the following k-mer-based linear UMI clustering algorithm. First, with the default setting (optimal for the length of the UMI sequence (16 bp) of the 10x Genomics 3’ v3.1 scRNA-seq product), k-mers (k=7) are generated for each raw UMI sequence. Next, for each raw UMI sequence, the following operations are performed: (1) for each existing UMI cluster, check whether the raw UMI shares at least one k-mer with the representative k-mer set of the UMI cluster; (2) if such UMI cluster can be identified, assign the raw UMI to the UMI cluster and update the representative k-mer set of the UMI cluster by selecting k-mers that are shared by at least 75% of all the raw UMI sequences belonging to the UMI cluster; (3) if such UMI cluster cannot be identified, a new UMI cluster is formed with the raw UMI as its sole member (the k-mers of the raw UMI becomes the representative k-mer set). Once all the grouped raw UMI sequences are assigned to UMI clusters, each UMI cluster’s most frequent UMI sequence is chosen as a representative UMI sequence (most likely the error-corrected UMI sequence of the raw UMIs). The UMI clustering results are stored using the ‘UB’ (corrected UMI sequence) and ‘UR’ (raw UMI sequence) tags. Furthermore, the total number of base pairs of the read covering the reference genome and the genome-encoded internal poly(A) tract length at the 3’ end of the read are stored using the ‘LE’ and ‘IA’ tags, respectively.

Finally, the workflow exports a barcoded BAM file containing CB and UMI correction results. Also, additional quality-control metrics of a long-read scRNA-seq experiment are collected and exported, including various histograms of the lengths (genome-aligned lengths, the values stored using the ‘LE’ tag) of the cDNA molecules after UMI deduplication, the cDNA molecules with both 3’ and 5’ PCR handles, the cDNA molecules with valid CB, and the cDNA molecules with different UMI duplication rates (e.g., cDNA molecules with no UMI duplication, cDNA molecules with a UMI duplication rate between 64-127, .) (**Supplementary Figure 3**).

***The biological full-length identification module***

A mature mRNA transcript (synthesized by RNA polymerase II) can be characterized by the presence of the 7-methylguanosine (m7G) cap and the enzymatically added poly(A) tail at its 5’ and 3’ ends, respectively. The next module of Ouro-Tools, the biological full-length identification module, identifies an “*in vivo*” full-length cDNA molecule containing the reverse transcription signatures of the m7G cap and the enzymatically added poly(A) tail, representing an intact mRNA *in vivo* (**Figure 3a**). Consequently, the cDNA molecules representing fragmented mRNA transcripts are excluded from downstream analysis.

The unique template-switching property of Moloney murine leukemia virus (M-MLV) reverse transcriptase (RT) generates a distinctive pattern in the resulting cDNA when the m7G cap is present at the 5’ end of an mRNA. The template switching property of M-MLV reverse transcriptase has allowed the efficient construction of a full-length cDNA library from very long mRNA molecules (>5 kbp); currently, M-MLV RT and the enzymes derived from M-MLV RT are utilized widely in the biotechnology industry, including every single-cell RNA-Seq platform that generates a full-length cDNA library (as an intermediate product). Recent studies revealed that the presence of m7G cap at the 5’ end of mRNA not only increases the template switching efficiency at the 5’ end^70,71^ but also results in precisely one “non-templated” cytidine addition^71^ before template switching occurs, possibly templated by the m7G cap itself^71^. Interestingly, changing the capping nucleotide from 7-methylguanosine to other nucleotides also changed the first nucleotide added before the template switching (C for the m7G cap) to the complementary nucleotide (T, G, and A) to the capping nucleotide (A, C, and U, respectively)^70^, further validating the proposed mechanism that the capping nucleotide itself templates the “non-templated” addition. Hence, it has been repeatedly observed that the location of the genuine transcription start site (TSS) strongly correlates with the presence of an unencoded guanosine nucleotide (i.e., an unencoded-G, which base pairs with the “non-templated” cytidine) found at a junction between the 3’ end of a template-switching-oligo-encoded sequence and the 5’ end of genome-encoded cDNA sequence, which has been utilized by few bioinformatics toolkits for identifying locations of *de novo* TSSs with significantly increased accuracy at bulk (CapFilter^72^) or single-cell level (SCAFE^73^) using short-read sequencing. However, there is currently no such bioinformatic method developed for Nanopore long-read sequencing, which has notably higher deletion error rates for G/C homopolymers^74^, which can cause significant problems during the analysis of soft-clipped G/C homopolymer sequences for the identification of an unencoded-G.

Since template-switching-oligos (TSOs) invariably ends with -rGrGrG-3’ (located at their 3’ ends) for efficient template switching by M-MLV RT, the “non-templated” cytosine addition due to the presence of the m7G cap at the 5’ end of a mature mRNA increases the total number of the unreferenced (not derived from the genome) guanosine nucleotides at the 5’ end of cDNA (oriented in the same direction as the original mRNA) from three (derived from the -rGrGrG-3’ sequence of TSOs) to four. Therefore, the biological full-length identification module collects the lengths of guanosine homopolymers at the 5’ ends of cDNAs to identify genuine TSSs that produce capped mRNAs. The module is implemented as a pipeline consisting of several separate workflows: the “LongSurvey5pSiteFromBAM”, “LongClassify5pSiteProfiles”, “LongAdd5pSiteClassificationResultToBAM”, and “FilterArtifactReadFromBAM” workflows (**Supplementary Figure 4**).

The “LongSurvey5pSiteFromBAM” workflow, as the first workflow of the module, collects the lengths of G (guanosine) and C (cytosine) homopolymers in the soft-clipped sequences starting from the 5’ ends of cDNAs aligned to positive and negative strands of the reference genome, respectively, for each TSS candidate (i.e., a 5’ site) for each sample; the lengths of G (or C) homopolymers are averaged and collected for each group of cDNAs sharing the same 5’ site with identical CB and UMI sequences; with the default setting, internal poly(A)-primed cDNAs (cDNAs lacking valid 3’ end signature) are excluded from the analysis. Additionally, for each 5’ site, the lengths of G (or C) homopolymers in the genome-aligned sequences at the 5’ ends of cDNAs are collected for accurate inference of the number of the unreferenced guanosine nucleotides aligned to the genome due to the chance occurrence of consecutive G (or C) nucleotides next to the 5’ site. Consequently, the workflow exports a histogram of the lengths of soft-clipped and genome-aligned G/C homopolymers (i.e., a 5’ site profile) for each TSS candidate for the sample. For efficient clustering and UMI-deduplication of reads based on the alignment locations of their 5’ ends on the reference genome, the tasks are split into batches for individual chromosomes for parallelization (all reads aligned to the same chromosome are analyzed together by a single worker process).

Next, the “LongClassify5pSiteProfiles” workflow combines the 5’ site profiles collected from individual samples and detects the signature of the m7G cap for each TSS candidate, identifying genuine TSSs. Again, the tasks are split into batches for individual chromosomes for parallelization (all the 5’ site profiles of TSS candidates of the same chromosome are analyzed together by a single worker process). First, histograms of the lengths of soft-clipped and genome-aligned G/C homopolymers are combined across the samples and converted into proportions using the total UMI-deduplicated read count for each TSS candidate. Next, with the default setting, the predefined score matrix (alternatively, the score matrix can be trained using the list of ground-truth TSSs) is utilized to classify each TSS candidate. Consequently, each 5’ site profile is classified into one of the following classes, which are ‘0_GGGG’, ‘0_GGG’, ‘-1_GGGG’, ‘-1_GGG’, ‘-2_GGGG’, ‘-2_GGG’, and ‘no_unrefG’; ‘GGGG’ and ‘GGG’ indicate the unreferenced G homopolymers at the 5’ ends of cDNAs from mRNAs with and without the m7G cap, respectively; ‘0’, ‘-1’, and ‘-2’ indicate the number of the unreferenced G nucleotides aligned to the genome coincidentally, displayed using the minus sign; when the unreferenced G homopolymers are not present at the 5’ ends of cDNAs, the 5’ site profile is classified into the ‘no_unrefG’ class, which represents an artifactual TSS generated by mispriming events. After classification, the locations and directions of the identified TSSs (the 5’ site profiles classified into one of the ‘0_GGGG’, ‘-1_GGGG’, and ‘-2_GGGG’ classes) are exported as an output of the workflow. Additionally, the classification results of the TSS candidates are visualized using interactive graphs to evaluate the classification process.

Subsequently, the “LongAdd5pSiteClassificationResultToBAM” workflow applies the TSS identification results of the previous workflow to the barcoded BAM file of each sample, generating a TSS-identified barcoded BAM file as an output of the workflow. Similarly, the tasks are split into batches for individual chromosomes for parallelization. If a cDNA starts from one of the identified TSSs, the read is marked with the flag indicating a valid TSS, which is stored using the ‘VS’ tag in the output TSS-identified barcoded BAM file; the lengths of soft-clipped and genome-aligned G/C homopolymers are stored using the ‘AG’ and ‘UG’ tags, respectively; the number of nucleotides in the genome-aligned G/C homopolymer that has not been originated from the genome (i.e., the length of the unreferenced G/C homopolymer that is coincidentally aligned to the genome) is stored using the ‘AU’ tag.

Lastly, the “FilterArtifactReadFromBAM” workflow classifies cDNAs based on the presence of the reverse transcription signatures of the m7G cap (valid 5’ end) and the enzymatically added poly(A) tail (valid 3’ end) and identifies the *in vivo* full-length cDNA molecules. With the default setting, a cDNA molecule is marked with a flag indicating a valid 3’ end when the poly(A) tail extends to the soft-clipped region at the 3’ end of the cDNA more than 8 bp. Also, a cDNA molecule is marked with another flag indicating a valid 5’ end when the cDNA starts from one of the identified TSSs or an unencoded-G is detected at the 5’ end of the cDNA. Consequently, the TSS-identified barcoded BAM file of each sample is split into the four BAM files, according to the flags indicating valid 3’ and 5’ ends for each cDNA molecule: the ‘valid_3p_valid_5p’ (*in vivo* full-length mRNAs), ‘valid_3p_invalid_5p’ (5’-degraded mRNAs and partial cDNA fragments generated by 5’ mispriming events), ‘invalid_3p_valid_5p’ (partial cDNA fragments generated by 3’ mispriming events), and ‘invalid_3p_invalid_5p’ (other partial cDNA fragments) BAM files. As a result, the module successfully depleted truncated cDNA molecules *in silico* (**Figure 3d**), which could significantly improve the sensitivity and accuracy of full-length transcriptome analysis of individual single cells. Particularly, Ouro-Tools utilizes the locations of the identified TSSs for downstream analysis, such as estimating the mRNA size distribution of a sample and assigning individual cDNA molecules to specific transcript isoforms accurately.

***The size distribution normalization module***

Since short-read RNA-Seq was widely adopted in the research community for quantitative transcriptome analysis, a plethora of normalization techniques have been developed to reduce the technical variability inherent in short-read RNA-Seq and increase the sensitivity for detecting actual biological variations between samples. Consequently, the normalization methods developed for bulk short-read RNA-Seq have been readily incorporated into single-cell RNA-Seq analysis frameworks, including the size factor normalization method that adjusts for the sequencing depth of each cell with the assumptions that most genes are not differentially expressed and that every cell has the similar number of mRNA molecules^75^. Since then, even though more sophisticated normalization algorithms have been developed to account for the technical variability unique to single-cell RNA-Seq, the size factor normalization method, despite its simplicity, currently remains one of the most widely used normalization methods for single-cell RNA-Seq data due to its robustness and consistency^76^.

The recent advent of long-read sequencing technologies has enabled comprehensive transcriptome analysis at the resolution of individual full-length mRNA and cDNA molecules. However, it also revealed a previously underappreciated type of technical variation inherent in all RNA-Seq methods: technical variations in the full-length mRNA and cDNA size distributions. For example, even if the cDNA size distributions of samples are almost identical and the same long-read sequencing technology is used for the samples, the sequencing process itself can introduce and exacerbate the differences in cDNA size distributions between the samples. Due to the much slower diffusion rate of larger cDNA molecules, we repeatedly observed that the read length distribution of a sample continuously changes as a Nanopore long-read sequencing run progresses, reaching a steady state ~20 hours after the run has started. However, a Nanopore sequencing run is often terminated before the steady state has reached to recover blocked pores with DNA nuclease treatment for increasing the total sequencing throughput for the flow cell (subsequently, the flow cell is reloaded with the same library or a library from a different sample for another run). Similarly, PacBio’s SMRT sequencing system also has a known preference for sequencing shorter molecules due to the much faster loading of the shorter SMRTbell templates to the SMRT sequencing system, often requiring a stringent removal of shorter cDNA molecules in order to characterize transcripts longer than 2 kbp with sufficient sequencing coverages (Pacific Biosciences Document #PN 100-378-900-01). Altogether, generating single-cell (or bulk) long-read RNA-Seq datasets with an identical size distribution of cDNA molecules is often not feasible.

Given that the size factor normalization method remains one of the most widely used methods for scRNA-seq data^76^, we sought to develop a normalization technique for long-read scRNA-seq that seamlessly integrates with the size factor normalization method. When we examined the 17,382 short-read RNA-Seq datasets covering 54 organs and tissues from the GTEx Consortium^77^, generated with the standardized RNA-Seq protocol, mRNA size distributions of most organs and tissues were virtually identical after removal of organ-specific “peaks” (**Extended Data Figure 5e**), representing highly abundant organ-specific mRNA transcripts (such as transcripts encoding β-Amylases for pancreas). Therefore, with the assumptions that most transcripts are not differentially expressed^78^ and that the underlying mRNA size distribution of each cell is nearly identical, we reasoned that the technical variability originating from the differences in the *in vivo* full-length cDNA size distributions between samples can be effectively reduced by constructing a “reference” mRNA size distribution that can represent all mRNA molecules in all the samples and adjusting the differences to the reference mRNA size distribution for each sample (**Figure 3a**).

The size distribution normalization module of Ouro-Tools is implemented using the “LongSummarizeSizeDistributions” and “LongCreateReferenceSizeDistribution” workflows. First, using the “LongSummarizeSizeDistributions” workflow, a full-length, UMI-deduplicated cDNA size distribution is obtained from the ‘valid_3p_valid_5p’ barcoded BAM file (representing *in vivo* full-length mRNAs) for each sample, one of the output files from the biological full-length identification module. Next, the reference mRNA size distribution is constructed for all the samples using the “LongCreateReferenceSizeDistribution” workflow (**Supplementary Figure 5**).

To accurately capture technical variations between the experiments, excluding the effects of highly abundant mRNA transcripts specific to certain cell types and experimental conditions is necessary. Consequently, the first part of the “LongCreateReferenceSizeDistribution” workflow is identifying underlying mRNA size distributions of individual samples by iteratively removing highly abundant mRNA species that manifests as a “peak” from the UMI-deduplicated, full-length cDNA size distribution of each sample (**Figure 3f**). Each iteration of the peak removal process performs the following steps. First, with the default setting, every “peak” that satisfies the following criteria is identified from the size distribution: at the 50% height of the peak, the width of the peak should be in the range from 0.5 bp to 50 bp; the number of mRNAs (UMI-deduplicated count) that is included in the area above the 50% height of the peak (i.e., the read count in the peak) should be at least 50 counts and should be larger than 5% of that of the area below the 50% height of the peak (i.e., the read count of the baseline); the height of the peak from to the baseline should be larger than the height of the baseline. Next, each identified peak is removed using the linear interpolation between the start and end positions of the peak on the baseline; to reduce noise, the heights at the start and end of the peak are replaced with the geometric means of 4 bp (the default setting) before and after the peak, respectively, before the interpolation. The iterative peak removal process is repeated up to 10 times until no new peak can be detected.

Next, the adaptive smoothing algorithm is applied to estimate each sample’s underlying mRNA size distribution (**Figure 3f**). The Gaussian smoothing process utilizes a Gaussian filter with a fixed standard deviation value to effectively reduce the localized noise of an image (including 1D images such as histograms). However, the Gaussian filter with a constant standard deviation does not account for the mean-variance relationship in the typical peak-removed mRNA size distribution (the region of higher sequencing coverages shows a higher local variation of sequencing coverages). We observed that applying a filter with a fixed standard deviation creates significant discrepancies between the smoothened and the original distributions in the low-coverage regions, especially when the filter with a large standard deviation is utilized to reduce the noise in the high-coverage regions. To account for the mean-variance relationship, with the default setting, a total of 25 Gaussian filters with standard deviation (σ) values between 4 and 64 (evenly spaced on a linear scale) are independently applied to the peak-removed mRNA size distributions. Subsequently, for each position of the size distribution (i.e., each length of mRNAs), the Gaussian filter with the largest standard deviation value that does not result in the skewed distribution deviating more than 50% from the minimally smoothened size distribution using the Gaussian filter with σ = 4 is selected. The resulting mRNA size distribution is cleaned using a Gaussian filter with σ = 3 to obtain an underlying mRNA size distribution for the sample, which is subsequently used to construct the reference mRNA size distribution.

Lastly, the reference mRNA size distribution of the samples is constructed as the geometric mean of the underlying mRNA size distributions of the samples to adequately represent all mRNA transcripts profiled in the samples (**Figure 3a**). Consequently, the optimal correction ratios are determined for each sample using a grid search algorithm; with the default setting, the grid search algorithm tries to make the relative ratios of the mRNA size distribution of the sample to the reference mRNA size distribution closer to 1 (1 indicates no correction is needed) for the size range of 1-3 kbp by searching the optimal scale factor in a range from 0 to 100.

***The single-cell long-read count module***

Previous studies have shown that long-read scRNA-seq enables accurate quantification of isoforms and splice junctions^79^ and locus-specific quantification of transposable-element-derived transcripts^22^ and regulatory element-derived transcripts. The last module of the standard Ouro-Tools pipeline, the single-cell long-read count module, not only enables accurate quantification of isoforms by utilizing the reverse transcription signatures of the 3’ and 5’ ends of full-length mRNA molecules but also supports single-cell analysis of insufficiently characterized genomic loci using long-read sequencing, including transcribed cis-regulatory elements (tCREs) and transcripts derived from transposable elements (TEs), while simultaneously reducing unwanted technical variations in read length distributions for accurate integration of multiple datasets (**Figure 3a**). Specifically, the module normalizes the mRNA size distribution of every cell in a sample using global scale factors calculated for the sample (i.e., the optimal correction ratios that were calculated by the previous module, the size distribution normalization module) by comparing the overall mRNA size distribution of the sample to the reference mRNA size distribution, before applying the size factor normalization method (or any other scRNA-seq normalization method) for individual cells in the sample.

The single-cell long-read count module is implemented as the “LongExportNormalizedCountMatrix” workflow (**Supplementary Figure 6**). The workflow is largely composed of three parts: constructing an index (only required once for each set of gene, transcript, repeat elements, and regulatory element annotations and the reference genome), assigning each read to various “buckets” (including genes, transcripts, exons, splice junctions, TE, tCRE, and individual genomic tiles), and exporting a size distribution-normalized count matrix for each “bucket” (later these count matrixes are combined into a single size distribution-normalized count matrix as an output). In the standard Ouro-Tools pipeline, the workflow is applied to the ‘valid_3p_valid_5p’ barcoded BAM file for each sample, representing *in vivo* full-length mRNAs of the sample.

Initially, an Ouro-Tools index is constructed using the given reference genome and annotations of various elements in the genome. For the construction of the mouse splicing atlas and the benchmarking of various long-read scRNA-seq methods, the following annotations were utilized. First, the reference genome and gene annotations from Ensembl^80^ were utilized (for *Homo sapiens*, GRCh38 and the Ensembl release 105 were utilized; for *Mus musculus*, GRCm38 and the Ensembl release 102 were utilized; for *Arabidopsis thaliana*, TAIR10 and the Ensembl Plants release 56 were utilized). For gene annotations, gene ID, gene name, transcript ID, and transcript name should be present in the attributes of the input GTF/GFF file (the gene annotations from Ensembl have such information). With the default setting, each group of genes with an identical gene name, an identical chromosome name, and an identical strand orientation are merged into a single gene. After the merge operation, the remaining groups of genes with identical gene names are renamed so that each gene name becomes unique across the genome (by adding a suffix like “_dup1”, “_dup2”, … ). Additionally, promoters are defined using the transcription start sites (TSSs) of different isoforms of each gene with a predefined length of 2,000 bp. For convenience, the Ensembl version number is dropped (with the default setting) from gene and transcript IDs. Next, for transposable element (TE) annotations, the repeat elements identified by the RepeatMasker program^81^ using the Repbase Update^82^ were utilized, which was downloaded from the UCSC Table Browser^83^ for *Homo sapiens* and *Mus musculus*. With the default setting, repeat elements smaller than 100 bp are excluded, and only the repeat elements of “SINE,” “LINE,” “LTR,” “DNA,” and “Retroposon” classes are selected and included in the resulting index. Lastly, the regulatory element annotations from the Ensembl Regulatory Build^84^ were utilized for *Homo sapiens* and *Mus musculus*. Regulatory element annotations should include regulatory element ID as an attribute in the input GTF/GFF file. Even though the transcription of an enhancer-derived RNA (eRNA) is initiated in both directions within the enhancer, it extends well beyond the enhancer element^85^. Therefore, the provided regulatory element annotations are expanded in both directions by 2,000 bp (with the default setting). Once the Ouro-Tools index is constructed for the organism (requires 0.9–15 GB of storage, depending on the organism), all subsequent runs can utilize the stored Ouro-Tools index without repeating the construction process.

Next, the module analyzes individual reads in the (TSS-identified) barcoded BAM file and assigns each read to “buckets” representing various genomic and transcriptomic features, such as genes, exons, splice junctions, intron retention events, transcripts, TEs, regulatory elements, and genomic bins. For parallelization, the tasks are split into batches for individual chromosomes (all reads aligned to the same chromosome are analyzed together by a single worker process). The assignment process hierarchically classifies each cDNA molecule into one of the following classes: transcripts of known genes, TE-derived transcripts, regulatory element-derived transcripts, and transcripts derived from previously uncharacterized genomic loci. First, for each read, all overlapping genes are identified and further analyzed to assign a single gene to the read at most. If the read overlaps with precisely a single gene or does not overlap with any gene, the gene assignment step ends here. However, when the read overlaps with more than one gene, first, the strand orientation of each gene is checked with the orientation of the cDNA aligned to the reference genome, and the genes with the mismatched strand orientation are excluded. If two or more genes overlap with the read after the filtering step, all exons overlapping with the read are retrieved, and the total number of overlapping base pairs is counted for each overlapping gene with the matched strand orientation. As a result, the gene with the largest overlapping base pairs at the exon level is identified and assigned to the cDNA. If the numbers of overlapping base pairs of the genes are the same and the decision cannot be made, the cDNA is marked using the bitwise flag “0x2” to indicate that the gene assignment was ambiguous (see **Supplementary Table 5** for the complete list of the bitwise flags utilized by the Ouro-Tools pipeline). Next, if the cDNA does not overlap with any gene (i.e., “intergenic”) or the cDNA is assigned to a single gene but only covers the introns of the gene (does not overlap with any exon of the gene), the cDNA is searched against the repeat elements. Similarly, if more than one repeat element overlaps with the cDNA, the total number of overlapping base pairs is counted for each overlapping repeat element, and the repeat element with the largest overlap is assigned to the cDNA (if a decision cannot be made, the cDNA is marked using the bitwise flag “0x80”, **Supplementary Table 5**). If the cDNA does not overlap with the repeat elements or the total proportion of the cDNA overlapping with the repeat element(s) is below a predefined threshold (with the default setting, 10%), the cDNA is searched against the regulatory elements. If more than one regulatory element overlaps with the cDNA, the regulatory element with the largest overlap is assigned to the cDNA, and if the decision cannot be made, the bitwise flag “0x1000” is utilized to mark the cDNA (**Supplementary Table 5**). Lastly, if the cDNA does not overlap with genes, repeat elements, and regulatory elements, the read is assigned to one of the genomic bins (as a “catch-all” category), which are non-overlapping regions of the reference genome with a predefine width of 100bp (with the default setting).

If a cDNA is successfully assigned to a single gene, its alignments to the reference genome and the isoform sequences of the gene are further analyzed to assign the cDNA to additional “buckets” representing various transcriptomic features. First, a bucket for the gene is initialized (if it has not been initialized) by building a gene-specific Minimap2 index (Minimap2 is one of the popular aligners for long-read RNA-seq) using the transcript sequences of the gene. Since the enzymatically added poly(A) tail is commonly not included in the transcript sequences, with the default setting, 50 bp of a poly(A) tail sequence is appended to the 3’ ends of the transcripts before constructing the index. Next, with the gene-specific Minimap2 index, the cDNA is aligned to the predefined transcripts of the gene; before the re-alignment process, the soft-clipped sequences of the read are removed, and the read is re-oriented so that its poly(A) tail is situated at the 3’ end like the transcript sequences. Subsequently, with the default setting, the alignments to the transcripts are filtered using the following criteria: the alignment is excluded when the read is aligned to the transcript in the anti-sense orientation; the alignment is excluded when soft-clipping(s), insertions, or deletions larger than 10 bp are present in the alignment; the alignment is excluded if the differences of transcript start site (TSS) and transcript end site (TES, or polyadenylation site) are larger than 25 bp and 100 bp, respectively (we utilized a more lenient threshold for the mismatch of polyadenylation sites since we often observed significant mismatches of the polyadenylation sites between the reference transcripts and the actual full-length cDNA molecules). After filtering, the transcript with the largest overlap is assigned to the cDNA. Once the cDNA is assigned to the specific transcript of the gene, the alignment of the cDNA to the transcript is further analyzed to identify overlapping exons and splice junctions of the transcript. Additionally, the alignment of the cDNA to the reference genome is independently analyzed to identify overlapping exons and splice junctions of the gene; the buckets representing the exons and splice junctions identified from the alignments to the transcripts of the gene can be differentiated from those identified from the alignments to the reference genome using the presence of the suffix “|realigned” in the bucket names, indicating the transcriptomic features are identified using the re-alignment process. All the classification results of individual reads are stored in the output BAM file using the predefined SAM tags (see **Supplementary Table 4** for the completed list of the SAM tags utilized by the module).

Finally, individual “buckets” are converted to count matrices, combined, and exported as a single-cell long-read count matrix. Since each bucket is processed independently from other buckets, the tasks are split into batches for individual buckets for parallelization. A group of reads in each “bucket” (multiple buckets can contain the same read) are UMI-deduplicated, size-distribution-normalized, and converted to a single-cell count matrix. The process is repeated two more times using a subset of the reads based on the strand orientation with an additional suffix marking the strandedness of the cDNAs; the groups of reads aligned to the forward and reverse strands with “|strand=+” and “|strand=-” suffixes, respectively; for a gene bucket, “|strand=sense” and “|strand=antisense” suffixes are used, according to the relative orientation of the cDNA molecules to the gene. During UMI deduplication, the read with the largest number of base pairs aligned to the genome is chosen for each group of reads with identical cell barcodes and UMI sequences. After UMI-deduplication, the mRNA size distribution normalization is performed by adjusting a weight that each UMI-deduplicated read contributes to the final single-cell count matrix based on the genome-aligned length of the read (excluding adapter sequences), using the optimal correction ratios for the sample that were calculated from the previous module. The resulting single-cell count matrices from all the buckets are combined into a combined count matrix as an output of the module. Additionally, for the overall evaluation of the module, the summary metrics (e.g., the total number of reads before UMI-deduplication, the total number of CPU time utilized for processing of the bucket, and the proportion of *in vivo* full-length cDNA molecules) are collected for each bucket and exported as an output file. Furthermore, a list of mRNA size ranges of interest can be given as one of the inputs of the workflow to export a separate count matrix using the subset of cDNA molecules for each mRNA size range of interest, which can be useful for the integration of multiple datasets with highly variable read length distributions.

***The differential transcript usage detection module***

For robust detection of alternative splicing events, we designed a bioinformatic algorithm that could flexibly accommodate an even higher degree of sparsity in transcript-level count data from single-cell long-read RNA-seq, compared to gene-level count data from conventional single-cell short-read RNA-seq. Without accounting for the effects of the high dropout rate of scRNA-seq, bimodal splicing patterns (e.g., each group of cells exclusively expresses a unique isoform) are often detected from cells with unimodal splicing pattern^86^ (e.g., all cells express several isoforms simultaneously at nearly constant ratios). Since pseudo-bulk approaches have been effective for accurate differential expression analysis in single-cell RNA-seq datasets with a high dropout rate^87^, we reasoned that a pseudobulk-based algorithm would be ideal for robust detection of cell type-specific alternative splicing patterns.

In the standard Ouro-Tools pipeline, the (mRNA size distribution-normalized) single-cell long-read count matrix from the single-cell long-read count module is further processed using the size factor normalization method that utilizes the total (size distribution-normalized) counts of each cell as a size factor for the cell, including the cDNAs aligned to intergenic regions (e.g., repeat elements, regulatory elements, and previously uncharacterized genomic loci). Next, the normalized count matrix (before log-transformation) is split into gene-specific normalized count matrices for individual groups of transcriptomic features belonging to the same gene (i.e., a gene, the transcripts, the exons, and the splice junctions of the gene) for parallelization. Consequently, the tasks are split into batches for individual genes for parallelization. Each gene-specific subset of the normalized count matrix is then converted to a gene-specific pseudobulk-level count matrix by randomly grouping each subset of the cells belonging to the same cell type into a pseudobulk sample based on their gene expression levels. With the default setting, for each cell type, a group of cells is randomly picked (with the predefined random seed, 42, for reproducibility) until the total normalized count of the gene exceeds 30 for the selected cells, which is subsequently aggregated into a pseudobulk sample; the aggregation process is repeated until all the cells belonging to the cell type are aggregated to pseudobulk samples. Subsequently, the normalized counts for each “sub-gene” transcriptomic feature of the gene (i.e., the transcripts, the exons, and the splice junctions of the gene) are converted to proportions relative to the normalized count of the gene for each pseudobulk sample (for a transcript feature, the proportion represents the transcript usage), which accounts for all the cDNA molecules in the pseudobulk sample that contributes to the counts of every “sub-gene” transcriptomic feature of the gene. Lastly, the proportions of each “sub-gene” transcriptomic feature of each cell type are compared to those of the rest of the cell types at the pseudobulk-level, using the various metrics and statistical tests, including the area under the receiver operating characteristic (AUROC), the Wilcoxon signed-rank test, a Chi-squared test, and a fold change between the average proportions of the pseudobulk samples of the cell type and the rest of pseudobulk samples.

***Other miscellaneous workflows***

Ouro-Tools includes other workflows for further processing and visualization of the TSS-identified barcoded BAM files, including the “DeduplicateBAM” and “SplitBAMs” workflows. The barcoded BAM files were processed using the following steps for the mouse splicing atlas datasets. First, the “DeduplicateBAM” workflow performs UMI-deduplication of cDNA molecules for each barcoded BAM file by selecting the largest cDNA molecule for each group of cDNA molecules with the identical cell barcode and UMI sequences for each barcode-attachment site on the reference genome (i.e., each polyadenylation site for the 3’ barcoding strategy-based scRNA-seq methods). Next, the “SplitBAMs” workflow splits the UMI-deduplicated barcoded BAM files of the samples of each organ into BAM files of individual cell types in the organ. Additionally, the “SplitBAMs” workflow splits the UMI-deduplicated barcoded BAM files of each sample into BAM files of individual cells in the sample, which are subsequently utilized for the analysis of coverage patterns at the single-cell level (**Figure 4c**).

*Tissue dissociation and sample preparation*

**Cell culture and preparation for SPLiT-Seq**

Mouse fibroblasts (NIH/3T3) and macrophages (RAW 264.7) were cultured in high-glucose DMEM medium (#SH30243.01, Cytiva) supplemented with 10% Fetal Bovine Serum (FBS) (#SH30919.03, HyClone) and penicillin-streptomycin (#SV30010, HyClone). Human immortalized T cells (Jurkat, Clone E6-1) were cultured in RPMI medium (#11875093, Gibco) supplemented with 10% FBS and penicillin-streptomycin. Cells were grown in a 37°C incubator with 5% CO_2_. Cells were harvested using trypsinization (#25-200-072, Gibco) and scraping for NIH/3T3 and RAW 264.7, respectively. Harvested cells were washed, resuspended in 0.04% BSA (#130-091-376, Miltenyi Biotec) in DPBS (#14190144, Gibco), and processed using the SPLiT-Seq protocol^3^.

**Mice**

C57BL/6J mice were obtained from the ORIENT BIO (Gyeonggi-do, Republic of Korea). Mice were provided food and water *ad libitum*, maintained on a 12-hour light/dark cycle, and housed under controlled temperature (22 °C) and humidity (55%) conditions in the Laboratory Animal Resource Center of the Gwangju Institute of Science and Technology. Female and male C57BL/6J mice were sacrificed at 8 weeks to collect samples for constructing the Mouse Single-Cell Long-Read Splicing Atlas. All animal experiments were approved (“GIST-2020-097”) by the Institutional Animal Care and Use Committee (IACUC) of the Gwangju Institute of Science and Technology.

**Leukocyte depletion based on differential centrifugation**

We utilized the Percoll gradient-based depletion method^88^ to deplete immune cells from single-cell suspensions from the mouse skin and mammary gland, intestine, and lung (**Supplementary Table 1**). First, 100% isotonic Percoll solution was prepared using pre-chilled Percoll (#17-0891-01, Cytiva) and 10X Hanks' Balanced Salt Solution (HBSS) (without Ca^2+^, Mg^2+^, and phenol red, #14185052, Gibco). Next, 42% isotonic Percoll solution^88^ was prepared by diluting the 100% isotonic Percoll solution with pre-chilled DMEM with red phenol (#SH30243.01, Cytiva). Additionally, 15% and 50% isotonic Percoll solutions were prepared by diluting the 100% isotonic Percoll solution with pre-chilled HBSS (without Ca^2+^, Mg^2+^, and phenol red). The chilled isotonic Percoll solutions were stacked below a single-cell suspension in a conical tube, starting from the lowest concentration (15%); from the bottom of the tube, 50%, 42%, and 15% isotonic Percoll layers, and the single-cell suspension layer are stacked on top of each other. The resulting Percoll gradients were centrifuged at 1,000 RCF at 4 ℃ for 10 minutes with no brake. Since leukocytes are denser^89^ (1.06–1.11 g/ml) than the 42% isotonic Percoll layer (~1.056 g/ml), cells between the 42% and 15% isotonic Percoll layers are depleted of immune cells, which were collected for further processing.

**Debris removal**

Debris was removed from the single-cell suspension using the Debris Removal Solution (#130-109-398, Miltenyi Biotec) before depleting red blood cells, following the manufacturer’s instructions.

**Lysis of red blood cells (RBCs)**

The cell pellet was resuspended in 1 ml of chilled 1X RBC Lysis Buffer (#420301, BioLegend) and incubated on ice for 10 minutes. Next, 10 mL of chilled single-cell wash buffer (0.04% BSA in DPBS) was added and centrifuged at 300 RCF at 4 ℃ for 10 minutes. The supernatant was removed, and the cell pellet was resuspended in 5 mL of the chilled single-cell wash buffer.

**Thymus**

Single cells from mouse thymus were provided by the Immune Synapse & Cell Therapy Research Center of Korea, Gwangju Institute of Science and Technology. After RBC lysis, the single-cell suspension was filtered using a 40 µm Flowmi cell strainer.

**Ovary**

Mouse ovary (8–10 weeks) was dissociated using the papain-based dissociation method described by Frost, E. R. et al.^90^.

**Testis**

Mouse testis (8–10 weeks) was dissociated using the method described by Jung, M. et al. ^91^.

**Skin and mammary gland**

Mouse skin and mammary gland (8–10 weeks) were dissociated using the Mouse Epidermis Dissociation Kit (#130-095-928, Miltenyi Biotec). Optionally, immune cells were depleted using the Percoll gradient-based method, enriching various keratinocytes and epithelial cells from the single-cell suspension.

**Intestine**

The small intestine of mice (8–10 weeks) was dissociated using the Mouse Lamina Propria Dissociation Kit (#130-097-410, Miltenyi Biotec). Optionally, immune cells were depleted using the Percoll gradient-based method, enriching enterocytes and other parenchymal cells (i.e., tuft cells) from the single-cell suspension.

**Lung**

Mouse lung (8–10 weeks) was dissociated with the Multi Tissue Dissociation Kit 2 (#130-110-203, Miltenyi Biotec) using the “Dissociation of rat lung using the Multi Tissue Dissociation Kit 2” protocol (Miltenyi Biotec). Also, another single-cell suspension was prepared from the mouse lung perfused with PBS to deplete circulating immune cells. Optionally, all leukocytes were depleted using the Percoll gradient-based method, enriching pulmonary endothelial cells and other parenchymal cells from the single-cell suspension.

**Kidney**

Mouse kidney (8–10 weeks) was dissociated with the Multi Tissue Dissociation Kit 2 (#130-110-203, Miltenyi Biotec) using the “Dissociation of mouse kidney using the Multi Tissue Dissociation Kit 2” protocol (Miltenyi Biotec).

**Kidney Glomeruli**

The glomeruli cells were enriched using magnetic beads^92^ (250 µl Dynabeads M-450 Tosylactivated, #65501, Invitrogen). Mice were perfused through transcardial puncture using PBS containing magnetic beads. The kidneys were then minced into 1 mm^3^ pieces and incubated in digestion buffer (1 mg/ml collagenase A (#COLLA-RO, Roche) in HBSS (#14025092, Gibco)) at 37 ℃ for 20 minutes. The tissue debris was removed using a 100 µm cell strainer, and the glomeruli were isolated through a magnetic separator for 10 minutes. The separated glomeruli were resuspended in digestion buffer (1 mg/ml collagenase A, 1 mg/ml Pronase E (#7433, Sigma-Aldrich), 50 µg/ml DNase I (#10104159001, Roche) in prewarmed PBS) and incubated at 37 ℃ for 30 minutes. Every 10 minutes, a 27-gauge needle was used to shear the sample during digestion. The suspension was centrifuged at 500 RCF at 4 ℃ for 5 minutes after being sieved through a 40 µm Flowmi cell strainer.

**Heart**

Mouse heart (8–10 weeks) was dissociated with the Multi Tissue Dissociation Kit 2 (#130-110-203, Miltenyi Biotec) using the “Dissociation of adult mouse heart using the Multi Tissue Dissociation Kit 2” protocol (Miltenyi Biotec).

**Heart Nuclei**

First, mouse heart (8–10 weeks) was dissociated with the Multi Tissue Dissociation Kit 2 (#130-110-203, Miltenyi Biotec) using a modified version of the “Dissociation of adult mouse heart using the Multi Tissue Dissociation Kit 2” protocol (Miltenyi Biotec) before nuclei isolation. A mouse heart was transferred into a 10 cm Petri dish containing 10 ml of PBS to remove blood from the organ. The heart was then minced into 1–2 mm^3^ pieces and transferred to a Gentle MACS C Tube (#130-093-237, Miltenyi Biotec) containing the prewarmed enzyme mix (#130-110-203, Miltenyi Biotec). The sample was dissociated using the Gentle MACS Octo Dissociator with Heaters (#130-096-427, Miltenyi Biotec) by running the program “37C_Multi_G” twice. The dissociation reaction was terminated by adding 7.5 ml of high-glucose DMEM medium (#SH30243.01, Cytiva) supplemented with 20% Fetal Bovine Serum (#SH30919.03, HyClone) to the sample. The sample was then centrifuged at 600 RCF for 5 minutes. The supernatant was removed, and the cell pellet was washed twice with single-cell wash buffer (0.04% BSA in DPBS). After applying the debris removal and RBC lysis processes described in the “Dissociation of adult mouse heart using the Multi Tissue Dissociation Kit 2” protocol (Miltenyi Biotec) to the sample, spontaneously beating, live cardiomyocytes (~100 µm in size) were observed under the light microscope, indicating successful dissociation of the mouse heart.

Next, nuclei were isolated from the isolated cells (Document #CG000365 Rev A, 10x Genomics). First, after removing the supernatant, 100 µl of the chilled lysis buffer (10 mM Tris-HCl [pH 7.4], 10 mM NaCl, 3 mM MgCl_2_, 0.1% Tween-20 (#1662404, Bio-Rad), 0.1% Nonidet P40 Substitute (NP40) (#74385, Sigma-Aldrich), 0.01% Digitonin (#BN2006, Invitrogen), 1% BSA, 1 mM DTT (#646563, Supelco), and 1 U/µl Protector RNase Inhibitor (#3335402001, Roche) in nuclease-free water (#AM9937, Invitrogen)) was added to the sample and mixed with the cell pellet by pipetting 10 times. The cell lysis reaction mixture was incubated on ice for 3 minutes. Subsequently, the lysis reaction was terminated by adding 1 ml of the chilled nuclei wash buffer (10 mM Tris-HCl [pH 7.4], 10 mM NaCl, 3 mM MgCl_2_, 1% BSA, 0.1% Tween-20, 1 mM DTT, and 1 U/µl Protector RNase Inhibitor in nuclease-free water) to the lysed cells, and the sample was centrifuged at 500 RCF at 4 ℃ for 5 minutes. The supernatant was removed, and the nuclei pellet was washed with 1 ml of the nuclei wash buffer 3 times. Lastly, the nuclei suspension was filtered using a 40 µm Flowmi cell strainer.

**Pancreas islet cell**

Mouse pancreatic islet cells were isolated using the previously described protocol^93^.

**Pancreas nuclei**

First, a mouse pancreas was transferred into a 10 cm Petri dish and rinsed with 10 ml of DPBS. Subsequently, the sample was minced into 1–2 mm^3^ pieces and transferred to a DNA LoBind tube (#0030108051, Eppendorf) containing 400 µl of the chilled (diluted) lysis buffer (a mixture of 100 µl of the lysis buffer (described above) and 300 µl of PBS) (Document #CG000124 Rev F, 10x Genomics). The sample was homogenized and incubated on ice for 2 minutes. Next, after adding 800 µl of the chilled nuclei wash buffer (described above), the homogenate was centrifuged at 500 RCF at 4 ℃ for 10 minutes. Subsequently, the supernatant was removed, and the nuclei pellet was washed twice with 1 ml of the nuclei wash buffer. The resulting nuclei suspension was filtered with a 40 µm Flowmi cell strainer.

**Brain**

The cerebral cortex and hippocampus of the mouse brain (8–10 weeks) were dissociated using the Mouse Adult Brain Dissociation Kit (#130-107-677, Miltenyi Biotec).

**Brain Nuclei**

Through dissection, the cerebral cortex and hippocampus of the mouse brain were transferred into a DNA LoBind tube (#0030108051, Eppendorf) containing 400 µl of the chilled (diluted) lysis buffer (a mixture of 100 µl of the lysis buffer (described above) and 300 µl of PBS) (Document #CG000124 Rev F, 10x Genomics). The sample was homogenized and incubated on ice for 4 minutes. Next, after adding 800 µl of the chilled nuclei wash buffer (described above), the homogenate was centrifuged at 500 RCF at 4 ℃ for 10 minutes. Subsequently, the supernatant was removed, and the nuclei pellet was washed twice with 1 ml of the nuclei wash buffer. The nuclei suspension was filtered with a 40 µm Flowmi cell strainer and stained with 7-Aminoactinomycin D (7-ADD) (#A1310, Invitrogen). Subsequently, debris was removed using fluorescence-activated cell sorting (BD FACSAria III, BD Biosciences), obtaining a single-nuclei suspension containing ~20,000 nuclei.

**Liver**

Mouse liver (8–10 weeks) was dissociated using the Mouse Liver Dissociation Kit (#130-105-807, Miltenyi Biotec). Optionally, hepatic non-parenchymal cells (NPCs) were enriched using the Percoll gradient-based method^94^, including hepatic stellate cells (HSCs), Kupffer cells (KCs), and liver sinusoidal endothelial cells (LSECs).

**Liver Nuclei**

First, a mouse liver was transferred into a 10 cm Petri dish and rinsed with 10 ml of DPBS. Next, the sample was minced^95^ into 1–2 mm^3^ pieces, which were transferred to a 15 ml conical tube containing dissociation solution (1 mg/ml collagenase type I (#17018029, Gibco) and 10 µM Rho-associated kinase inhibitor (#M1817, Abmole Bioscience, Houston, TX, USA) in DPBS). The tissue was dissociated at 37 ℃ for 10 minutes. Subsequently, the sample was centrifuged at 500 RCF for 5 minutes. After removing the supernatant, the cell pellet was washed twice with wash buffer (0.04% BSA and 10 µM Rho-associated kinase inhibitor in DPBS). Next, nuclei were isolated from the dissociated cells (Document #CG000365 Rev A, 10x Genomics); after the chilled lysis buffer (described above) was mixed with the cell pellet, the cell lysis reaction mixture was incubated on ice for 3 minutes. The resulting nuclei suspension was filtered with a 40 µm Flowmi cell strainer.

### SUPPLEMENTARY TABLES AND FIGURES

(Supplementary_Table_1.xlsx)

**Supplementary Table 1: Comprehensive summary of the 22 Ouro-Seq-applied long-read scRNA-seq datasets for the Mouse Single-Cell Long-Read Splicing Atlas.**

(Supplementary_Table_2.xlsx)

**Supplementary Table 2: Comprehensive summary of benchmarking results of various long-read scRNA-seq methods.**

(Supplementary_Table_3.xlsx)

**Supplementary Table 3: List of DNA oligo sequences utilized for sgRNA synthesis for Ouro-Deplete and Ouro-Enrich (Ouro-Cell-Enrich) experiments.**

(Supplementary_Table_4.xlsx)

**Supplementary Table 4: Proportions of long-reads representing full-length cDNAs across genes for the Ouro-Seq libraries prepared from *Mus musculus*.**

(Supplementary_Table_5.xlsx)

**Supplementary Table 5: Numbers of detected genes, isoforms, splice junctions, exons, transcribed regulatory elements, and transcribed transposable elements for each cell type.**

| SAM tag name | data type | Description | Module name |
| --- | --- | --- | --- |
| CB | Z | the corrected cell barcode sequence | Barcode Extraction |
| UB | Z | the corrected UMI sequence after the UMI clustering process | Barcode Extraction |
| UR | Z | the uncorrected UMI sequence before the UMI clustering process | Barcode Extraction |
| XR | i | the number of errors for identification of R1 adapter (marks the 3’ end of cDNA). -1 indicates that the adapter was not identified | Barcode Extraction |
| XT | i | the number of errors for identification of TSO adapter (marks the 5’ end of cDNA). -1 indicates that the adapter was not identified | Barcode Extraction |
| CU | Z | the uncorrected raw CB-UMI sequence | Barcode Extraction |
| IA | i | the length of detected internal poly(A) tract on the genome | Barcode Extraction |
| LE | i | the total number of genome-aligned base pairs | Barcode Extraction |
| AG | i | the number of consecutive G nucleotides, starting from the 5’ site in the aligned region of the read | Full-Length Identification |
| UG | i | the number of consecutive G nucleotides, starting from the 5’ site in the unaligned region of the read (soft-clipped sequence) | Full-Length Identification |
| VS | i | “1” if the 5’ site is identified as a valid transcript start site (TSS), “0” if the 5’ site is identified as an invalid TSS, representing 5’ sites of the PCR/RT artifacts and degraded mRNAs | Full-Length Identification |
| AU | i | the inferred number of unreferenced G nucleotides aligned to the genome | Full-Length Identification |
| XC | i | the bitwise flags (see **Supplementary Table 7** for more details) | Single-Cell Count |
| XR | Z | the repeat element ID | Single-Cell Count |
| YR | i | the total number of base pairs overlapping with the repeat element to which the read is confidently assigned | Single-Cell Count |
| XG | Z | the gene ID | Single-Cell Count |
| YG | i | the total number of base pairs overlapping with the exons of the gene to which the read is confidently assigned | Single-Cell Count |
| XP | Z | the promoter ID | Single-Cell Count |
| YX | i | the total number of base pairs overlapping with any exons that overlap with the read | Single-Cell Count |
| YF | i | the total number of base pairs overlapping with any repeat elements that overlap with the read (considering only filtered repeat elements) | Single-Cell Count |
| XE | Z | the regulatory element ID | Single-Cell Count |
| YU | i | the total number of base pairs overlapping with any repeat elements that overlap with the read (considering all repeat elements) | Single-Cell Count |
| YE | i | the total number of base pairs overlapping with any regulatory elements that overlap with the read | Single-Cell Count |
| XT | Z | the transcript ID that is uniquely assigned to the read using the re-alignment process | Single-Cell Count |
| ZF | i | the flag that indicates the read represents a full-length cDNA with valid 3’ and 5’ ends | Single-Cell Count |

**Supplementary Table 6. The list of SAM tags utilized by Ouro-Tools.**

| Binary flag | Feature type | Description |
| --- | --- | --- |
| 0x1 | gene | overlaps with gene(s) |
| 0x2 | gene | gene assignment is ambiguous |
| 0x4 | gene | completely intronic reads |
| 0x8 | gene | exonic reads |
| 0x10 | promoter | overlaps with promoter region(s) |
| 0x20 | promoter | promoter assignment is ambiguous |
| 0x40 | repeats | overlaps with repeat element(s) |
| 0x80 | repeats | ambiguous assignment to two or more number of repeat elements |
| 0x100 | repeats | the entire length of a read overlaps with a single repeat element |
| 0x200 | regulatory | overlaps with regulatory element(s) |
| 0x400 | regulatory | overlaps with both repeat element(s) and regulatory element(s) |
| 0x800 | regulatory | overlaps exclusively with regulatory element(s) (no overlaps with repeat element) |
| 0x1000 | regulatory | ambiguous assignment to two or more number of regulatory elements |
| 0x2000 | regulatory | the entire length of a read overlaps with a single regulatory element |

**Supplementary Table 7. The list of bitwise flags utilized by the single-cell long-read count module of Ouro-Tools for indicating the classification results of individual reads.**

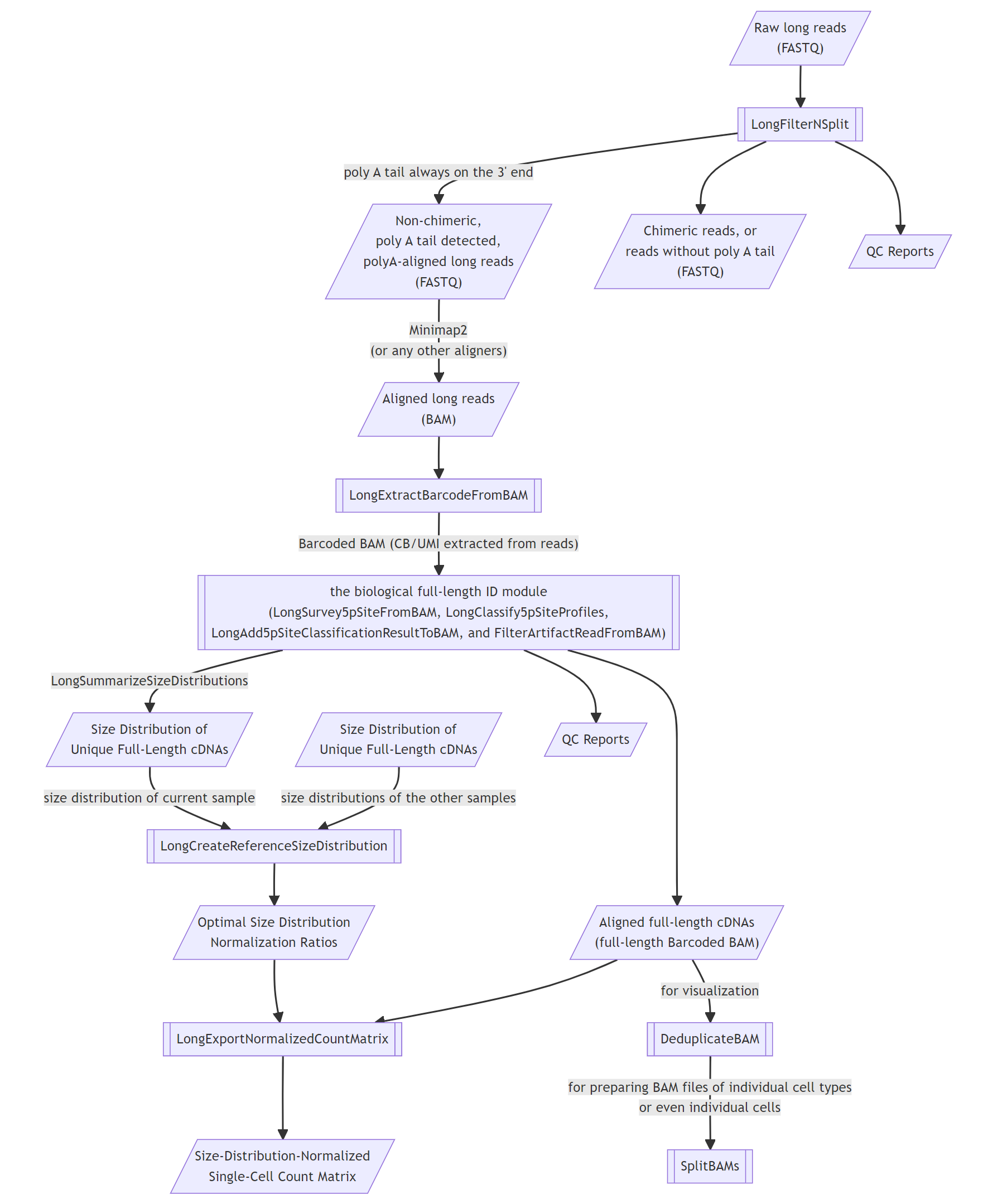

**Supplementary Figure 1.** An overview of the Ouro-Tools pipeline.

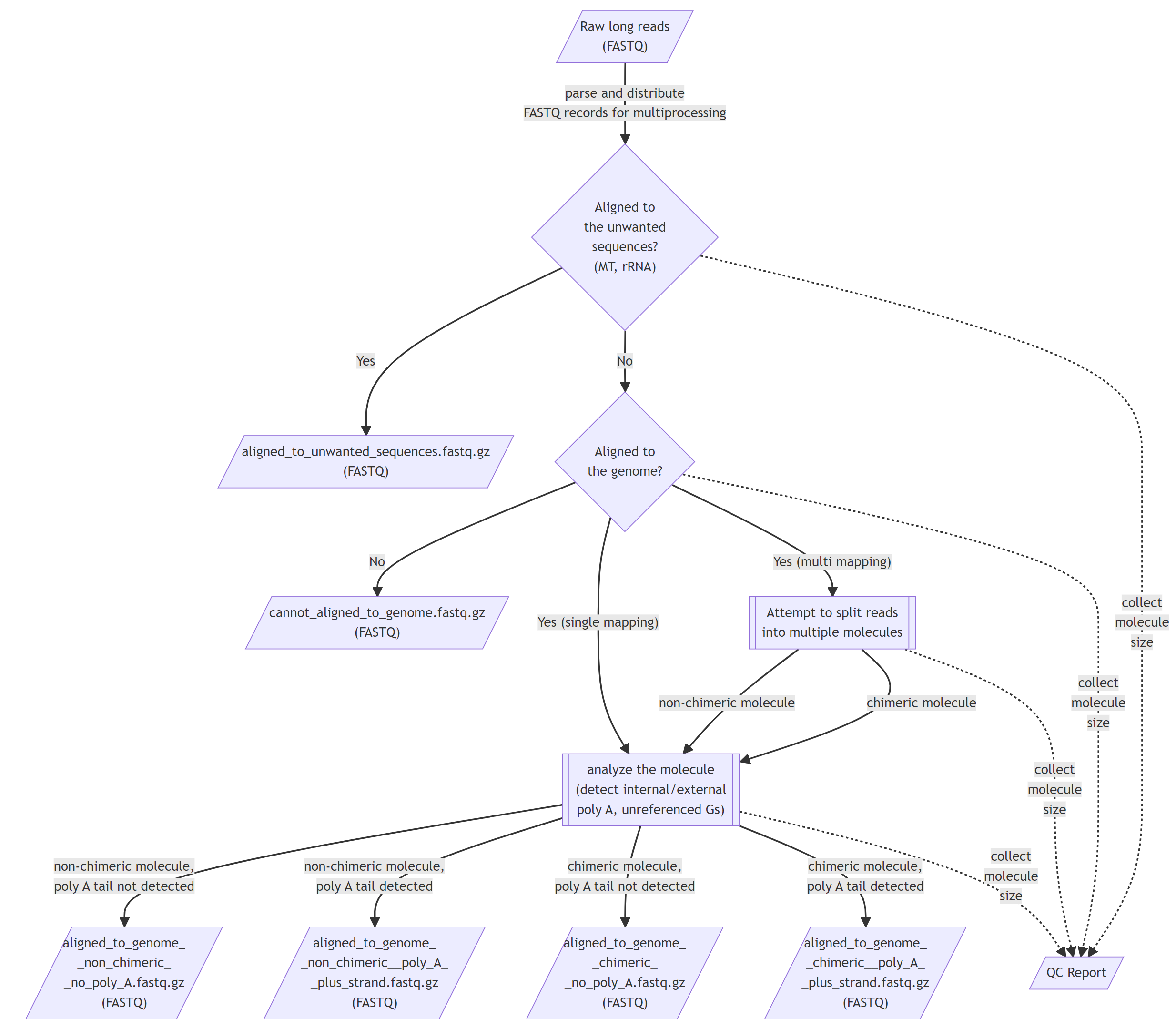

**Supplementary Figure 2.** A flowchart representing the “LongFilterNSplit” workflow of the Ouro-Tools pipeline.

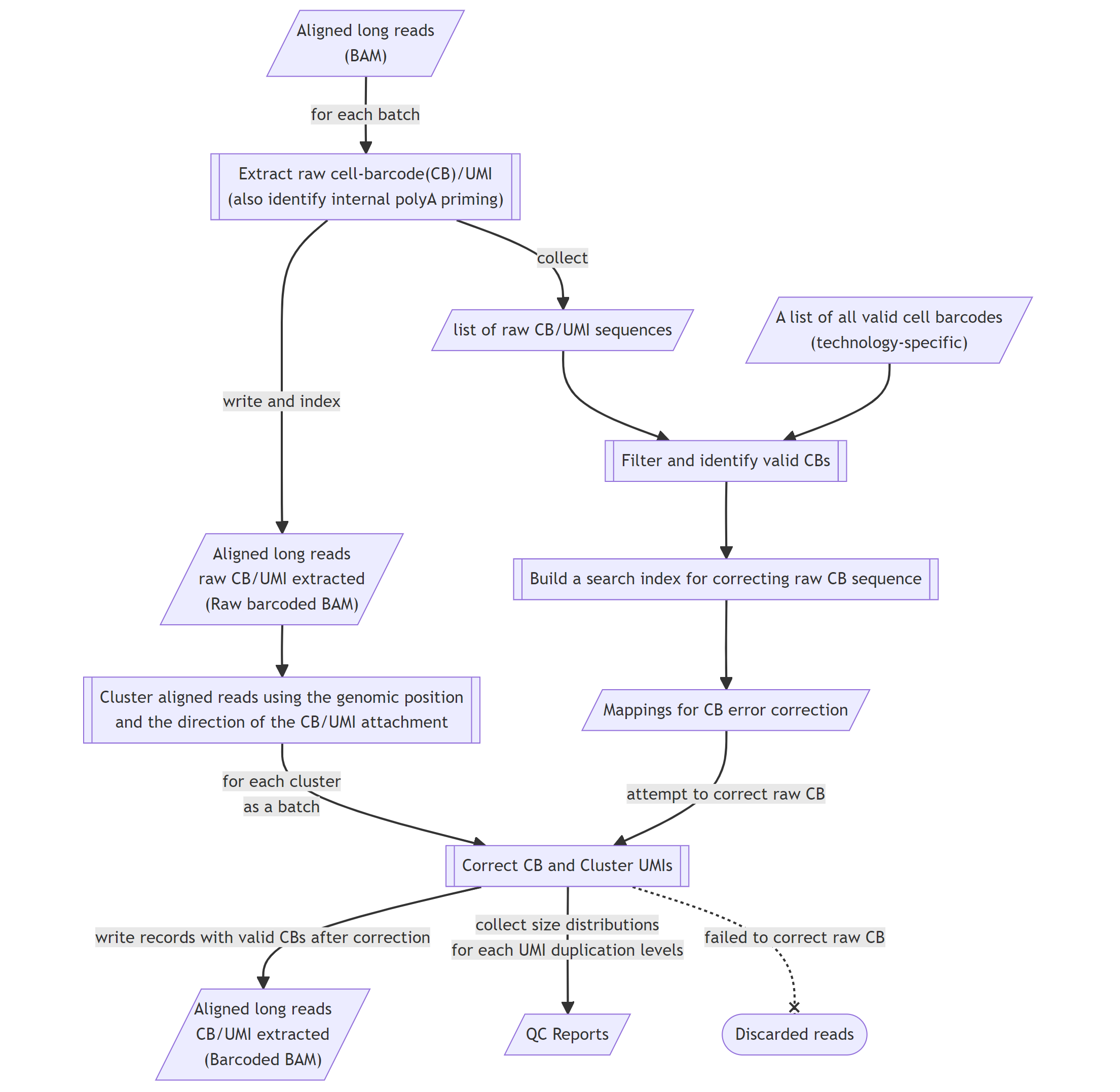

**Supplementary Figure 3.** A flowchart representing the “LongExtractBarcodeFromBAM” workflow of the Ouro-Tools pipeline.

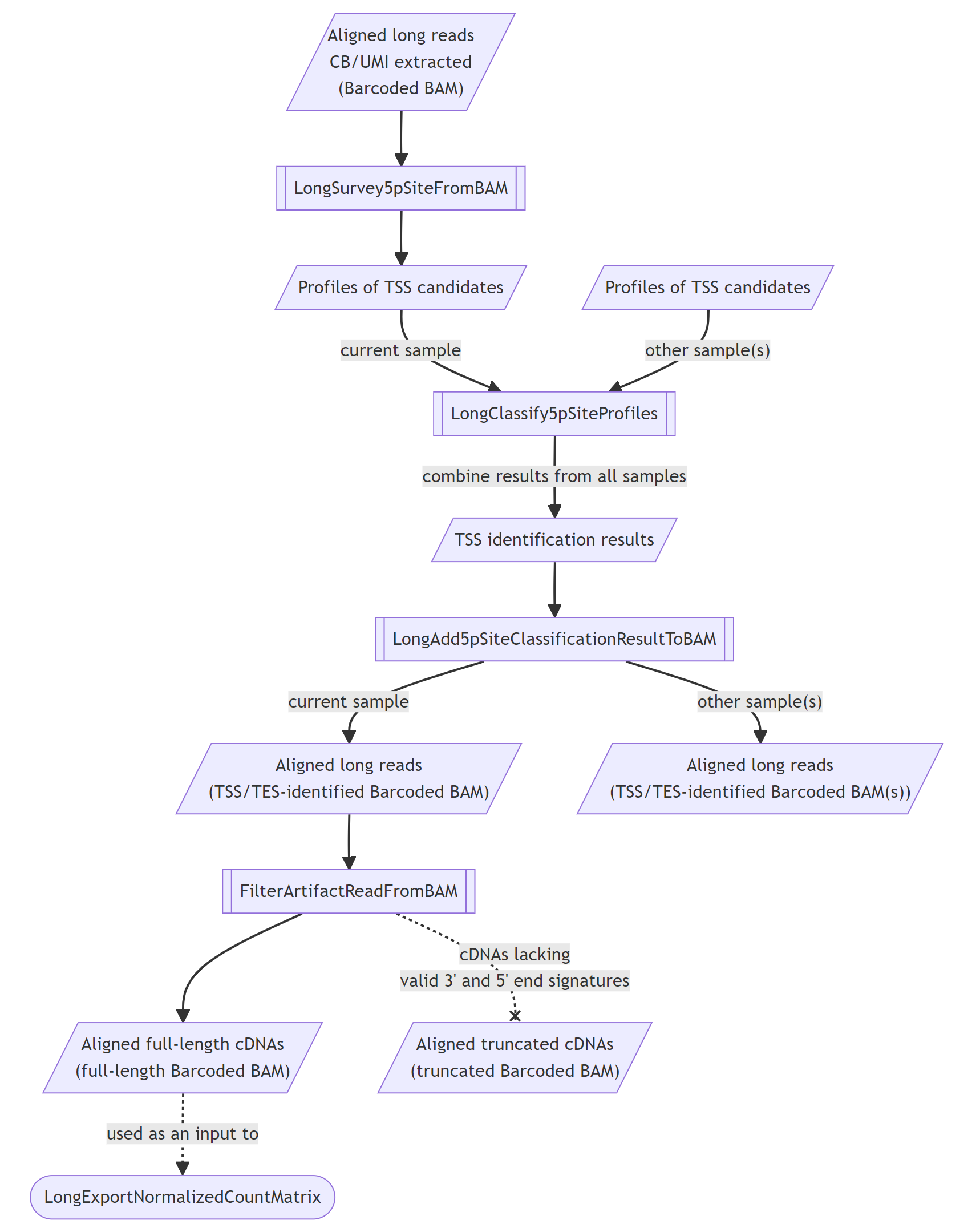

**Supplementary Figure 4.** A flowchart representing the biological full-length identification module of the Ouro-Tools pipeline.

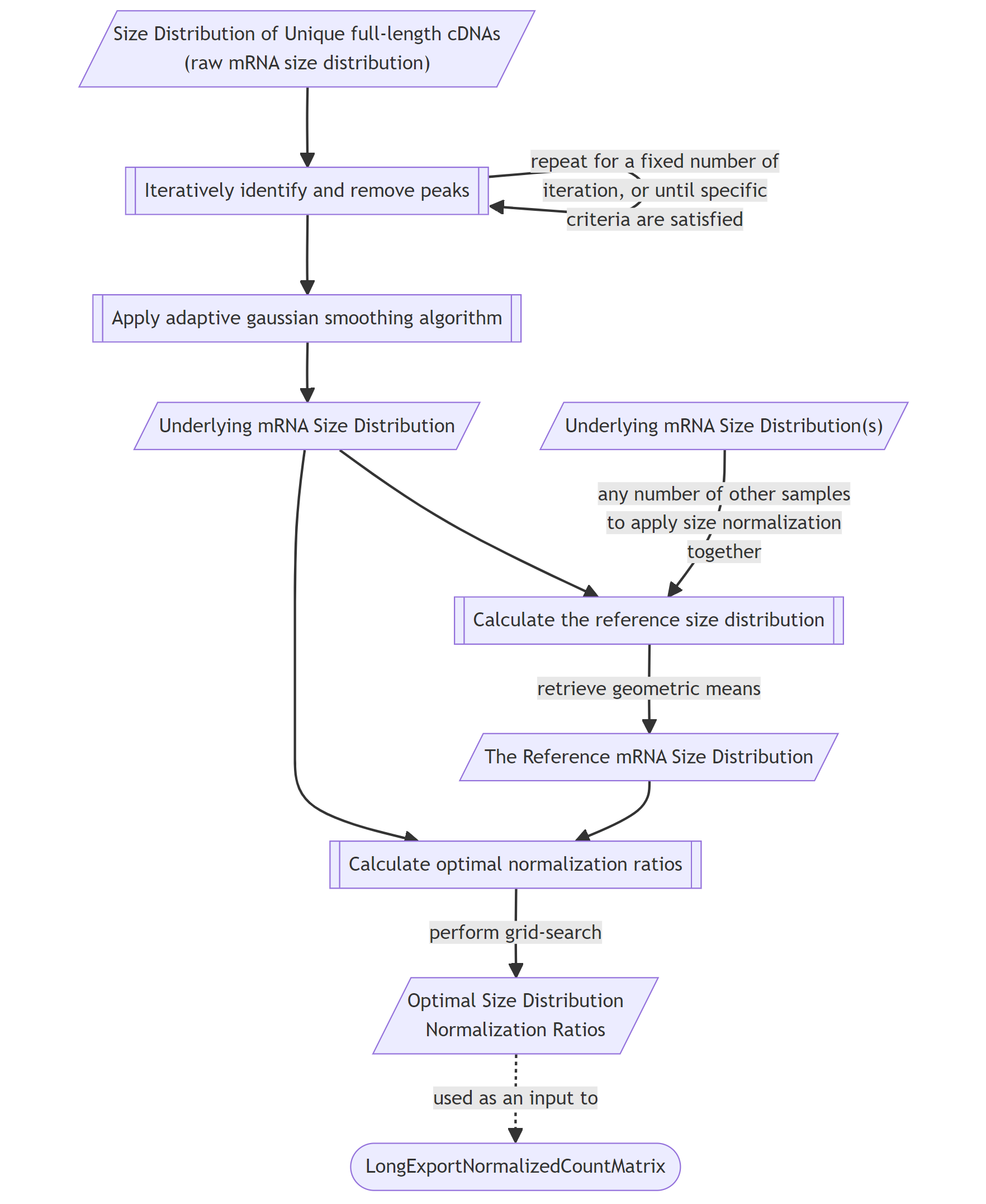

**Supplementary Figure 5.** A flowchart representing the “LongCreateReferenceSizeDistribution” workflow of the Ouro-Tools pipeline.

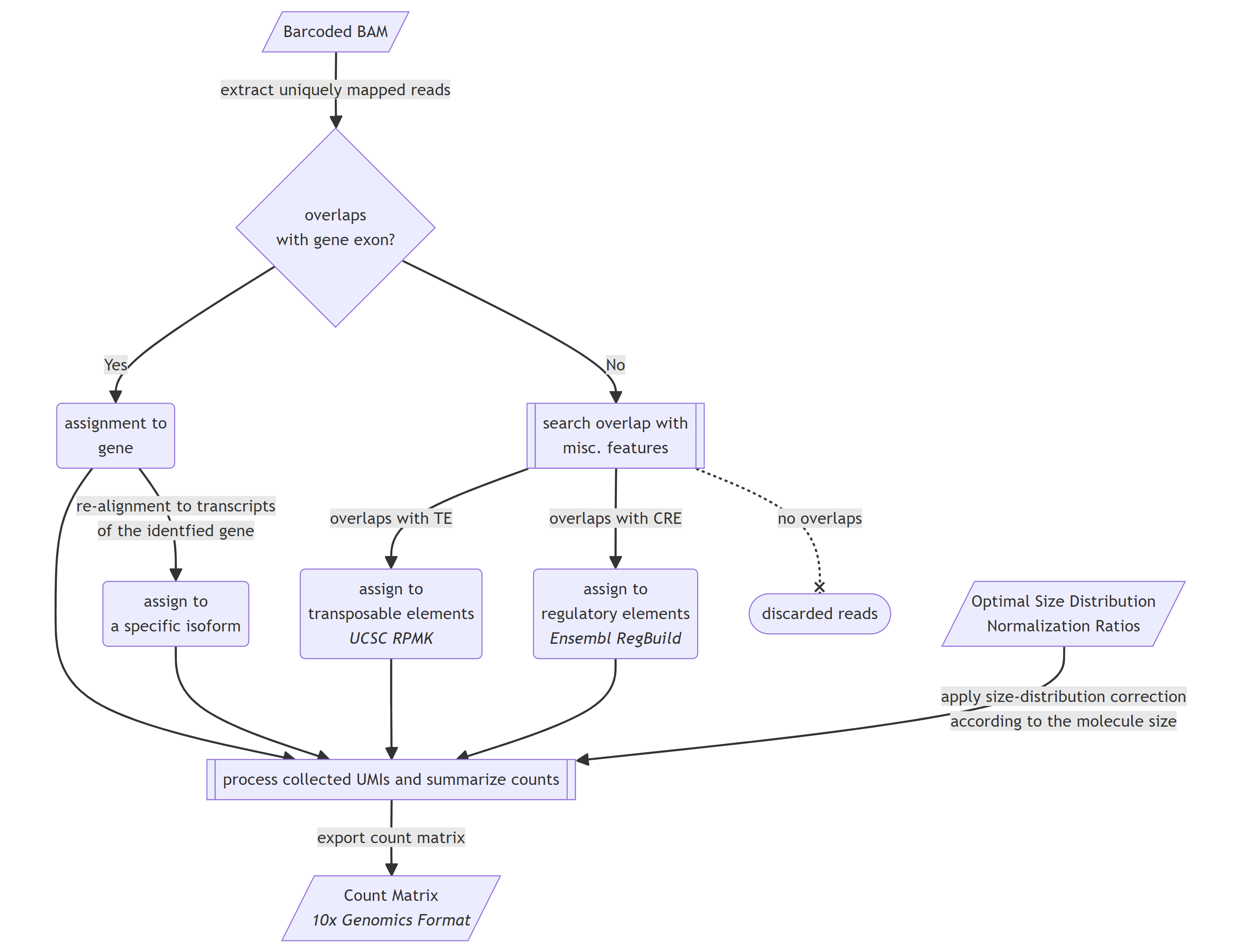

**Supplementary Figure 6.** A flowchart representing the “LongExportNormalizedCountMatrix” workflow of the Ouro-Tools pipeline.
